# Nano3P-seq: transcriptome-wide analysis of gene expression and tail dynamics using end-capture nanopore cDNA sequencing

**DOI:** 10.1101/2021.09.22.461331

**Authors:** Oguzhan Begik, Gregor Diensthuber, Huanle Liu, Anna Delgado-Tejedor, Cassandra Kontur, Adnan Muhammad Niazi, Eivind Valen, Antonio J. Giraldez, Jean-Denis Beaudoin, John S. Mattick, Eva Maria Novoa

## Abstract

RNA polyadenylation plays a central role in RNA maturation, fate, and stability. In response to developmental cues, polyA tail lengths can vary, affecting the translation efficiency and stability of mRNAs. Here, we develop Nanopore 3’ end-capture sequencing (Nano3P-seq), a novel method that relies on nanopore cDNA sequencing to simultaneously quantify RNA abundance, tail composition and tail length dynamics at per-read resolution. By employing a template switching-based sequencing protocol, Nano3P-seq can sequence any given RNA molecule from its 3’ end, regardless of its polyadenylation status, without the need for PCR amplification or ligation of RNA adapters. We demonstrate that Nano3P-seq captures a wide diversity of RNA biotypes, providing quantitative estimates of RNA abundance and tail lengths in mRNA, lncRNA, sn/snoRNA, scaRNA, and rRNA molecules. We find that, in addition to mRNA and lncRNA, polyA tails can be identified in 16S mitochondrial rRNA in both mouse and zebrafish models. Moreover, we show that mRNA tail lengths are dynamically regulated during vertebrate embryogenesis at an isoform-specific level, correlating with mRNA decay. Finally, we identify non-A bases within polyA tails of various lengths and reveal their distribution during vertebrate embryogenesis. Overall, Nano3P-seq is a simple and robust method for accurately estimating transcript levels, tail lengths, and tail composition heterogeneity in individual reads, with minimal library preparation biases, both in the coding and non-coding transcriptome.

## Introduction

RNA molecules are subject to multiple co- and post-transcriptional modifications, shaping them to their final mature form ^1^. Polyadenylation of RNA is one such modification, which is known to affect the stability and translation efficiency of the RNA molecule ^2–4^ and to play an essential role in determining the fate of RNA molecules in a wide range of biological processes ^5,6^.

One context in which polyadenylation has been shown to play a major role in determining RNA fate and decay is vertebrate embryogenesis ^6^. Indeed, in the first hours post-fertilization, vertebrate embryos undergo major cellular reprogramming, a process known as the maternal-to-zygotic transition (MZT) ^7^. During the MZT, maternally inherited RNA and proteins are responsible for activation of the zygotic genome and are later replaced by the zygotic program ^8,9^. Because the MZT begins in a transcriptionally silent embryo, this transition relies heavily on post-transcriptional regulatory mechanisms ^7^, including modulation of the polyadenylation status of the RNA molecules ^6,10^. Therefore, characterizing the dynamics of RNA polyadenylation is key to understanding how these modifications regulate the fate and function of RNA molecules.

In the last few years, several transcriptome-wide methods have become available for studying the dynamics of polyadenylated tails (polyA tails) based on next-generation sequencing (NGS), such as PAL-seq or TAIL-seq ^10,11^. While these methods have been successfully employed to characterize the dynamics of polyA tail lengths in various contexts, they have several important caveats: (i) they provide a limited perspective on isoform–tail relationships due to the short-read-length nature of NGS-based technologies; (ii) they do not provide single-molecule resolution; (iii) they are severely affected by polymerase chain reaction (PCR) amplification biases; and (iv) they can only measure tail lengths that are shorter than the read length.

To overcome these limitations, the direct RNA sequencing (dRNA-seq) platform offered by Oxford Nanopore Technologies (ONT) has been proposed as a means to study both the transcriptome and polyA tail lengths simultaneously ^12,13^. To sequence native RNAs using dRNA-seq, polyA-tailed RNA molecules are ligated to a 3’ adapter that contains an oligo(dT) overhang (**Figure 1a**). Consequently, dRNA-seq libraries capture the full-length polyA tail; however, ligation occurs only on RNA molecules that anneal to the oligo(dT) overhang, thus exclusively capturing polyA transcripts with tail lengths greater than 10 nt. A variation of this method consisting of *in vitro* poly(G/I)-tailing the transcriptome prior to library preparation has been proposed for studying nascent RNAs using dRNA-seq ^14,15^, thus capturing both polyadenylated and non-polyadenylated mRNAs. However, major limitations to this variation include low sequencing yields compared with standard dRNA-seq (10%–30%) ^14^ and a lack of tools to distinguish poly(I) and poly(A) signals; therefore, polyA tail length information is lost in these datasets ^14,15^. An alternative approach for studying the transcriptome using nanopore technologies is direct cDNA sequencing (dcDNA-seq), but this approach cannot sequence the polyA (-) transcriptome, nor can it capture polyA tail length information (**Figure 1a**). Overall, both dRNA-seq and dcDNA-seq nanopore library preparation protocols are limited to the sequencing of polyadenylated transcripts and thus cannot provide a comprehensive view of both polyadenylated and deadenylated RNA molecules, in addition to being unable to capture RNA molecules with other types of RNA tails (e.g., polyuridine).

**Figure 1.**
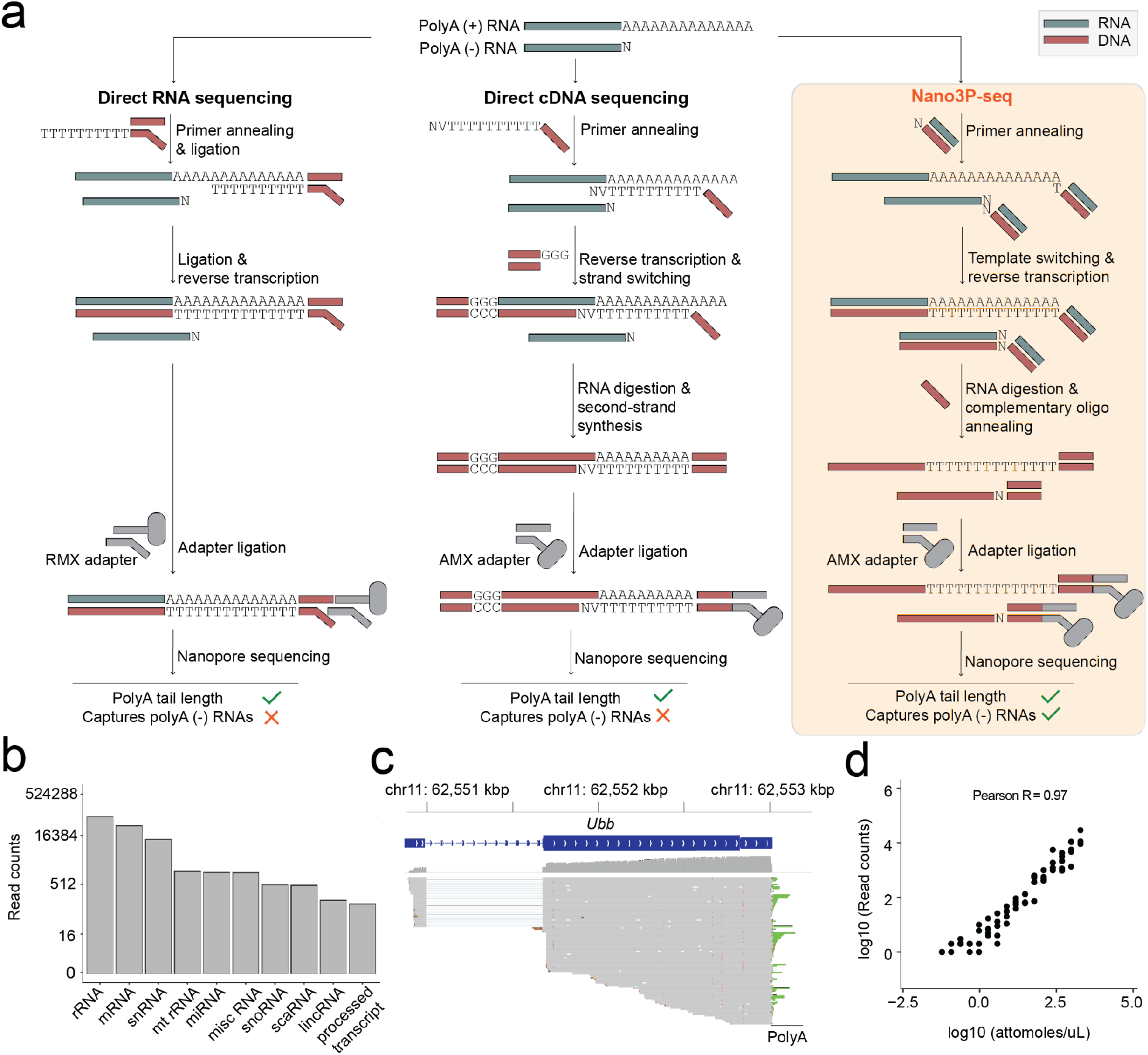
Nano3P-seq captures polyadenylated and non-polyadenylated RNAs, while retaining polyA tail length information. **(a)** Schematic overview comparing three different library preparation methods for studying the transcriptome using nanopore sequencing: i) standard direct RNA nanopore sequencing (left panel), ii) standard direct cDNA nanopore sequencing (middle panel) and iii) Nano3P-seq (right panel). **(b)** Nano3P-seq captures a wide range of RNA biotypes in a mouse brain nuclear/mitochondrial RNA sample. **(c)** Integrative Genome Viewer (IGV) snapshot of reads generated with Nano3P-seq, mapped to the *Ubb* gene, illustrating the diversity of polyA tail lengths captured across different reads. The polyA tail region is shown in green. **(d)** Scatter plot of log-transformed concentrations (attomoles/µL) and read counts of sequin genes (Pearson’s R : 0.93, slope: 0.93). Each dot represents a sequin. See also Figure S1h–i. Abbreviations: RMX: RNA adapter mix (provided with the SQK-RNA002 direct RNA sequencing library preparation kit); AMX: adapter mix (provided with the SQK-DCS109 direct cDNA sequencing library preparation kit); rRNA: ribosomal RNA; mRNA: messenger RNA; snRNA: small-nuclear RNA; mt rRNA: mitochondrial ribosomal RNA; miRNA: microRNA; misc RNA: miscellaneous RNA; snoRNA: small-nucleolar RNA; scaRNA: small Cajal body-specific RNA; lincRNA: long intergenic non-coding RNAs.

Here, we present a novel method that employs nanopore sequencing to simultaneously obtain per-isoform transcriptome abundance and tail lengths in full-length individual reads, with minimal library preparation steps, which we term **<u>Nano</u>**pore **<u>3 P</u>**rime end-capture **<u>seq</u>**uencing (Nano3P-seq) (**Figure 1a**). Notably, Nano3P-seq uses template switching to initiate reverse transcription and, therefore, does not require 3’ end adapter ligation steps, PCR amplification, or second-strand cDNA synthesis. We demonstrate that Nano3P-seq can capture any type of RNA molecule regardless of its 3’ sequence, including polyA-tailed and non-tailed RNA. Moreover, we show that Nano3P-seq can accurately quantify RNA abundances in both the coding and non-coding transcriptome, as well as quantify the polyA tail lengths and tail composition of individual RNA molecules in a highly reproducible manner.

## Results

### Nano3P-seq captures both polyadenylated and non-polyadenylated RNA molecules in a quantitative and reproducible manner

Because nanopore sequencing is typically limited to the analysis of polyA (+) RNA molecules (**Figure 1a**), previous efforts have opted to perform *in vitro* polyadenylation reactions of the total RNA to capture non-polyadenylated RNA in the sequencing run^16^. While this option can capture any given transcript present in the sample, it also leads to a loss of polyA tail length information. Therefore, we reasoned that by coupling template switching to cDNA nanopore sequencing, we would simultaneously capture the polyA (+) and polyA (-) transcriptome, while retaining polyA tail length information from each individual RNA molecule (**Figure 1a**).

To assess the ability of Nano3P-seq to sequence both polyA (+) and polyA (-) RNA, we first sequenced two synthetic RNAs, one lacking a polyA tail and one that had been *in vitro* polyadenylated (see *Methods*) (**Figure S1a**–**c**). Our results show that Nano3P-seq efficiently captures both polyadenylated and non-polyadenylated RNA molecules, as well as the diversity of polyA tail lengths in individual RNAs (**Figure S1c**). We then examined the performance of Nano3P-seq in *in vivo* samples, and sequenced total RNA samples from mouse brain, previously enriched in nuclear and mitochondrial content via subcellular fractionation to increase the content of non-coding RNAs ^17^ (see *Methods*). We confirmed that Nano3P-seq captured RNA biotypes that are typically polyadenylated (i.e., mRNA, lincRNA, processed transcript) as well as non-polyadenylated (i.e. rRNA, miscRNA, snRNA, snoRNA), the majority of them being rRNA, mRNA, and snRNA (**Figure 1b,** see also **Figure S1d and Table S1**). In addition, our results confirmed that polyA tail length information was retained in individual reads. Specifically, the majority of reads corresponding to mRNAs had polyA tails (**Figure 1c,** see also **Figure S1e,f**), whereas non-coding RNAs such as snoRNAs (**Figure S1f)** and snRNAs did not have polyA tails (**Figure S1g**), as expected.

To assess the accuracy and reproducibility of Nano3P-seq in quantifying RNA abundances, we examined the performance of Nano3P-seq in synthetic RNA mixes (sequins) ^18^ that had been spiked into samples in independent flow cells (**Table S2**). Our results showed that Nano3P-seq provided accurate estimates of RNA abundances both at the per-gene (Pearson’s R: 0.93, slope: 0.93) (**Figure 1d**) and per-isoform (Pearson’s R: 0.89, slope: 0.92) **(Figure S1h)** level. These quantifications were highly reproducible across biological replicates, both at the per-gene (Pearson’s R: 0.98) and per-isoform (Pearson’s R: 0.97) (**Figure S1i**) level. We should note that previous works using Illumina RNA-seq reported a global correlation of 0.89 and 0.86 between observed and expected sequin counts at per-gene and per-transcript level, respectively ^18^.

### Nano3P-seq recapitulates the dynamics of coding and non-coding RNAs during vertebrate embryogenesis

Next, we employed Nano3P-seq to examine the RNA dynamics that occur during the MZT at single-molecule resolution (**Figure 2a**). To this end, we isolated total RNA from zebrafish embryos at 2, 4, and 6 hours post-fertilization (hpf) in biological duplicates, ribodepleted the samples, and sequenced them using the Nano3P-seq protocol **(Figure S2a,b,** see also *Methods*). Quantification of the mRNA abundances in three independent biological replicates showed that per-gene abundances (reads per million, RPM) obtained using Nano3P-seq were highly reproducible (**Figure 2b,** see also **Figure S2c,d**).

**Figure 2.**
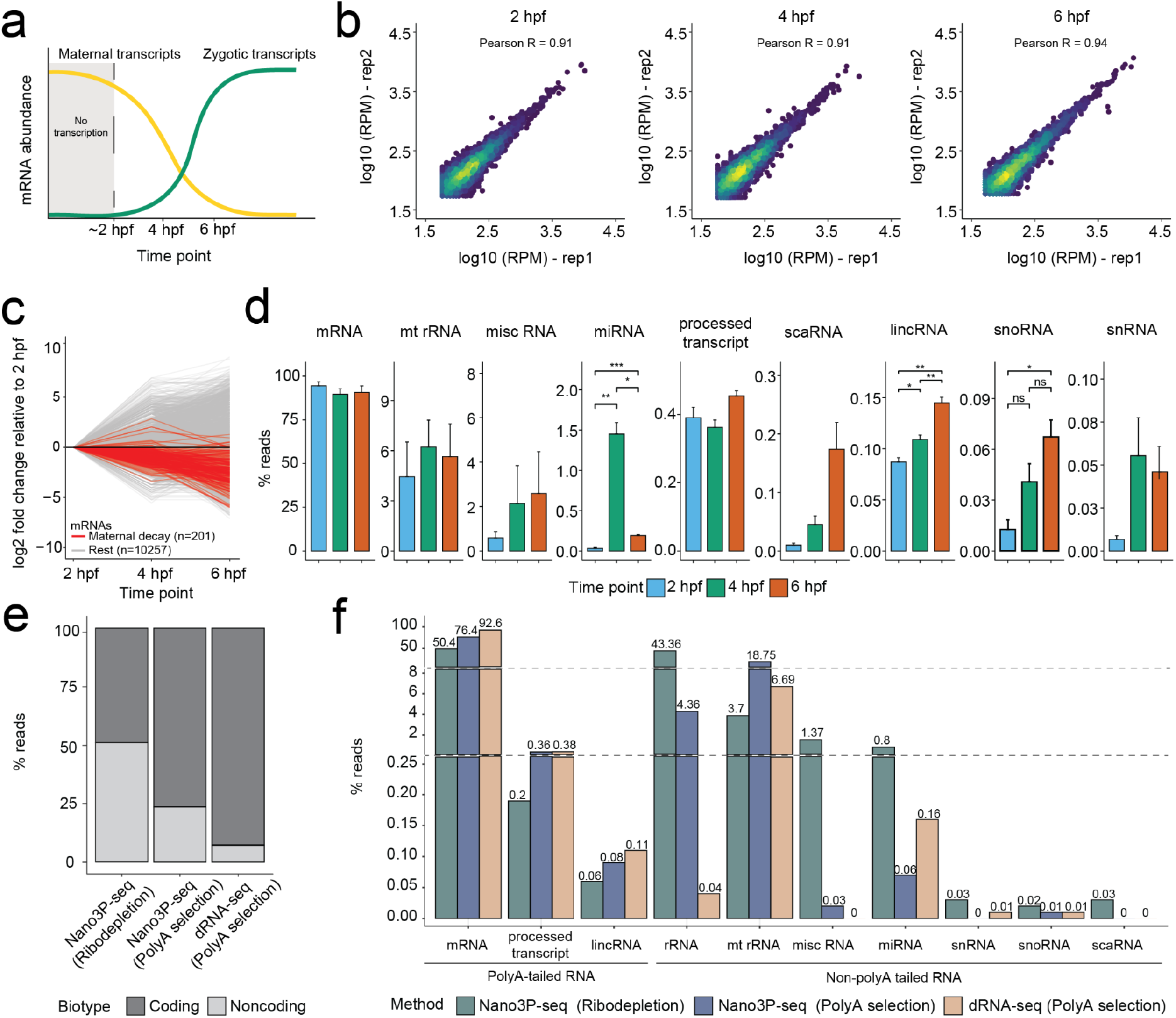
Nano3P-seq captures a wide diversity of coding and non-coding RNAs and their expression dynamics during the maternal-to-zygotic transition (MZT). **(a)** Schematic overview of the transcriptional change that occurs during the MZT in zebrafish. **(b)** Scatter plots depicting the correlation of mRNA log-transformed reads per million (RPM) between biological replicates at three different time points during the MZT. **(c)** Changes in mRNA abundance during the MZT (t = 2, 4, and 6 hours post-fertilization [hpf]), relative to 2 hpf. Genes previously reported to have a “maternal decay mode” are depicted in red. **(d)** Barplots depicting the abundance of different RNA biotypes captured by Nano3P-seq during the MZT (2, 4, and 6 hpf, shown in blue, green, and red, respectively). Statistical analyses were performed using the Kruskal–Wallis test (*n* = 3 biological replicates, error bars indicate the standard deviation). **(e)** Relative proportion of coding and non-coding RNAs captured using dRNA-seq (on polyA-selected samples), Nano3P-seq (on polyA-selected samples), and Nano3P-seq (on ribodepleted samples). **(f)** Percentage of reads mapping to distinct biotypes captured using Nano3P-seq (on ribodepleted samples) (green), Nano3P-seq (on polyA-selected samples) (blue), and dRNA-seq (on polyA-selected samples) (light brown). See also Figure S2 (P > 0.05:ns, p ≤ 0.05:*, p ≤ 0.01:**, p ≤ 0.001:***, p ≤ 0.0001:****).

A comparative analysis of mRNA population dynamics across time points showed that Nano3P-seq recapitulated the transcriptomic switch that occurs during the MZT ^7,8^, with a decay of mRNA genes previously reported to have a “maternal decay mode” (**Figure 2c, Table S3**), in agreement with previous studies ^19,20^. Notably, in addition to polyadenylated RNAs, Nano3P-seq captured a wide variety of RNA biotypes without polyA tails that are also present in early embryo stages. We observed a significant increase in the abundance of lincRNAs and snoRNAs during the MZT, as well as a sharp increase in miRNA expression at 4 hpf, followed by a decrease at 6 hpf (**Figure 2d**). Analysis of individual miRNA gene expression patterns during the MZT revealed that the sharp increase observed at 4 hpf was primarily caused by increased expression of genes belonging to the miR-430 family, which is known to be essential in mediating the decay of a group of maternal mRNAs during the MZT (mirR-430-dependent decay) ^21^ (**Figure S2e**). By contrast, much fewer non-coding RNA populations were globally captured when dRNA-seq was applied to the same samples (**Figure 2e**).

We noted, however, that mitochondrial rRNAs were not enriched in Nano3P-seq datasets relative to dRNA-seq datasets (**Figure 2f**). Indeed, per-read analysis of zebrafish mitochondrial rRNA reads revealed that a significant proportion of 16S mitochondrial rRNA contained a polyA tail, which explains the lack of enrichment of mitochondrial rRNAs in Nano3P-seq datasets relative to dRNA-seq datasets (**Figure S2f**). In agreement with this observation, we found that polyA-tailed 16S mitochondrial rRNAs were present not only in zebrafish (**Figure S2g**), but also in mouse (**Figure S2h,i**), suggesting that this feature is conserved across species and not a sequencing artifact, in agreement with previous reports ^22,23^. The presence of polyA tails in mouse 16S mitochondrial rRNAs was further validated using an orthogonal method (polyA tail length assay coupled to Sanger sequencing, see *Methods*), which confirmed that 16S mitochondrial rRNA is polyadenylated (**Figure S2j**).

### PolyA tail lengths can be accurately estimated using Nano3P-seq

We then examined whether Nano3P-seq accurately estimated polyA tail lengths. We note that algorithms for detecting polyA tails in native RNA nanopore sequencing reads are well established and benchmarked ^12,24–26^, but their applicability to cDNA reads, such as those from Nano3P-seq, remains unclear. To this end, we first examined whether the *tailfindR* polyA tail prediction software ^25^ would capture the presence or absence of polyA tails on synthetic RNAs that were either polyadenylated or non-polyadenylated and had been sequenced using Nano3P-seq. We found that *tailfindR* can identify both polyadenylated and non-polyadenylated Nano3P-seq reads (**Figure 3a, S3a)**. Then, we assessed the accuracy of the polyA tail length predictions of *tailfindR* in Nano3P-seq datasets that included a battery of synthetic RNAs (sequins) ^18^ or synthetic cDNA sequences with known polyA tail lengths (**Figure 3b**). Our results showed that polyA tail length estimations of sequins in Nano3P-seq data were highly reproducible across replicates (R: 0.993, **Figure S3b**), with an accuracy similar to that observed when performing polyA tail length estimations in sequins that had been sequenced using dRNA-seq (**Figure 3c,** see also **Figure S3c** and **Table S4**). Moreover, the variance of tail length estimates across reads belonging to the same transcript was smaller in Nano3P-seq datasets than in dRNA-seq datasets (**Figure S3d,e**). Similar results were obtained when we assessed the performance of the algorithm in a set of synthetic cDNA sequences that spanned a broader range of polyA tail lengths (0, 15, 30, 60, 90, 120 nt) (**Figure 3b,c,** see also **Figure S3f**).

**Figure 3.**
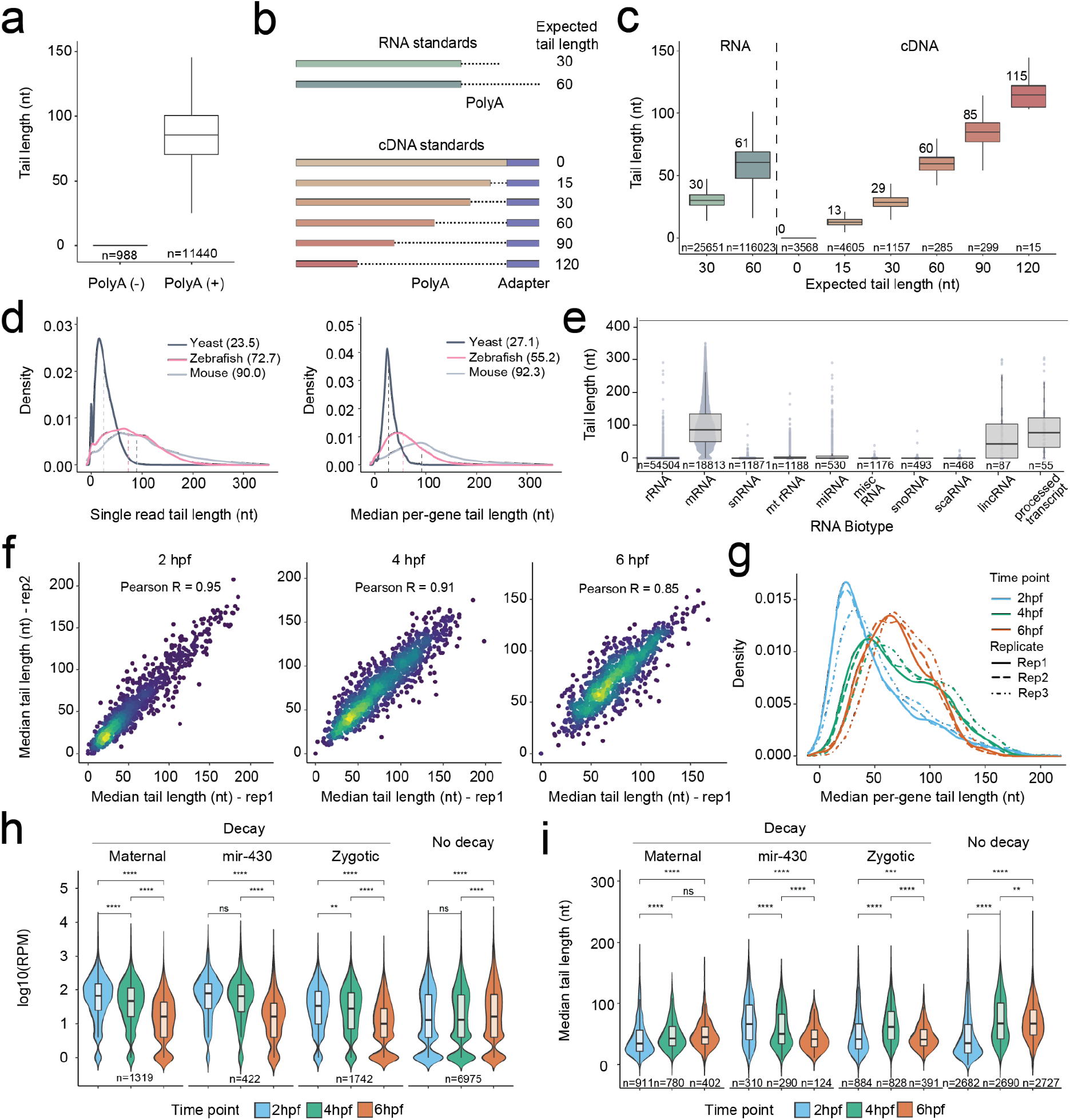
Nano3P-seq can be used to accurately estimate polyA tail lengths in individual molecules. **(a)** PolyA tail length estimates of non-polyadenylated (curlcake 1) and polyadenylated (curlcake 2) synthetic RNAs sequenced with Nano3P-seq. See also Figure S1a–c. **(b)** Schematic overview of the standards used to assess the tail length estimation accuracy of Nano3P-seq. **(c)** Boxplots depicting tail length estimations of RNA and cDNA standards sequenced with Nano3P-seq. Values on boxplots indicate the median polyA tail length estimation for each standard. The number of reads included in the analysis is shown below each boxplot. **(d)** PolyA tail length distribution of yeast, zebrafish, and mouse mRNAs represented as single-transcript values (left panel) and per-gene medians (right panel). **(e)** PolyA tail length estimates across different RNA biotypes from mouse brain total RNA enriched in nuclear/mitochondrial RNA. Each dot represents a read. The number of reads included in the analysis is shown below each boxplot. **(f)** Replicability of median per-gene polyA tail length estimations of zebrafish embryonic mRNAs between two biological replicates for three different time points (2, 4, and 6 hpf). **(g)** Median per-gene polyA tail length distribution of zebrafish embryonic mRNAs across zebrafish developmental stages (2, 4, and 6 hpf, shown in blue, green, and red, respectively) in three biological replicates (shown as full lines, dashed lines, and dotted/dashed lines, respectively). **(h)** Comparative analysis of mRNA abundances (shown as log10 RPM counts) of zebrafish mRNAs binned according to their annotated decay mode (maternal decay, zygotic activation-dependent decay, miR-430-dependent decay, and no decay) during early embryogenesis (t = 2, 4, and 6 hpf). Statistical comparison of means was performed using the Kruskal–Wallis test. The number of genes included in the analysis is shown below each violin plot. **(i)** Median per-gene polyA tail length estimations of zebrafish mRNAs binned according to their decay mode (maternal, miR-430, zygotic, and no decay) at 2, 4, and 6 hpf. Statistical analyses were performed using the Kruskal–Wallis test. The number of genes included in the analysis is shown below each violin plot. Boxplot limits are defined by lower (bottom) and upper (top) quartiles. The bar indicates the median, and whiskers indicate +/- 1.5X interquartile range. (P > 0.05:ns, p ≤ 0.05:*, p ≤ 0.01:**, p ≤ 0.001:***, p ≤ 0.0001:****). See also Figure S3.

Finally, we performed a comparative analysis of mRNA polyA tail lengths from mouse, yeast, and zebrafish. We observed that mouse mRNAs had the longest mRNA tails among the three species, with a median polyA tail length of 90 nt, whereas the shortest polyA tail lengths were observed in yeast, with a median polyA tail length of 23.5 nt (**Figure 3d**), in agreement with previous studies ^10^.

### Charting polyA tail length dynamics *in vivo* with Nano3P-seq

We explored whether Nano3P-seq could be used to investigate polyA tail length dynamics *in vivo*. To this end, we first examined the ability of Nano3P-seq to identify which RNA biotypes were polyadenylated in mouse brain total RNA samples that had been previously enriched in nuclear/mitochondrial content to increase the proportion of ncRNAs. We found that polyA tails were mainly predicted on mRNAs, but also in lincRNAs and processed transcripts, which are also known to be polyadenylated ^27,28^ (**Figure 3e,** see also **Table S1**).

We next analyzed the polyA tail length dynamics across developmental stages of zebrafish mRNAs during the MZT (t = 2, 4, and 6 hpf). PolyA tail length estimates were highly reproducible across independent biological replicates sequenced in independent flow cells for all three time points studied (R: 0.85–0.95) (**Figure 3f**). We observed an overall increase in the mean mRNA polyA tail length during the MZT (**Figure 3g**, see also **Figure S3g**), in agreement with previous reports^10^. All mRNAs examined were found to be polyadenylated, with the exception of histone mRNAs, which had a median polyA tail length of zero (**Figure S3h,** see also **Table S3**), in agreement with previous works reporting their non-polyadenylated status ^29^. These findings show that Nano3P-seq can capture RNA molecules with structured 3’ ends, such as those found in histones ^30^. Finally, we note that per-gene polyA tail length estimates obtained by Nano3P-seq during the MZT showed a good correlation with those obtained using orthogonal methods such as PAL-seq ^10^ (Pearson’s R=0.71-0.85) (**Figure S3i,j**)..

Next, we examined the correlation between polyA tail length dynamics and mRNA decay. To this end, mRNA transcripts were binned depending on their decay mode (maternal decay, zygotic activation-dependent decay, miR-430-dependent decay, and no decay), as previously described ^21^. We observed that the three groups of mRNAs that are known to decay (maternal, zygotic, and miR-430) showed a significant decrease in mRNA abundance (**Figure 3h, Table S3**), as expected. However, the patterns of polyA tail length dynamics strongly varied depending on the decay mode of the transcript (**Figure 3i, Table S3**). Specifically, we observed that transcripts that decayed in an miR-430-dependent manner showed a significant decrease in polyA tail length during the MZT, in agreement with previous studies^19,21^. By contrast, for mRNAs with the zygotic genome activation-dependent decay mode, this shortening only occurred after 4 hpf, and maternal mRNAs did not present a decrease in polyA tail length, but instead showed a consistent increase in tail length throughout the MZT. These observations are consistent with the reanalysis of PAL-seq data (**Figure S3k,l**). Overall, these results show that not all decay modes are associated with a reduction in transcript polyA tail lengths and demonstrate the applicability of Nano3P-seq to identify polyadenylated RNA populations, study their RNA abundance, and estimate their polyA tail length dynamics, at both the global level and the level of individual transcripts, and thus provide mechanistic insights into different gene regulatory programs.

### Nano3P-seq captures isoform-specific differences in polyA tail dynamics during the MZT

A major feature that distinguishes nanopore sequencing from NGS is its ability to produce long reads, allowing RNA polyadenylation dynamics to be studied at the isoform level. Therefore, we wondered whether Nano3P-seq could identify differentially polyadenylated transcript isoforms during the MZT.

To this end, we first examined whether polyA tail lengths significantly diverged across time points at the per-isoform level. We note that only reads mapping to genes encoding for at least two annotated isoforms, and with mapping coverage greater than 10 reads per isoform, were maintained for further analyses (**Table S5**). Our analyses revealed that 55.3% (± 8.62%) of analyzed transcripts varied significantly in polyA tail length during the MZT (**Table S6**). Notably, we observed that analyses at the per-gene level often revealed a different picture compared with analyses at per-isoform level. For example, in *khdrbs1a* and *syncrip*, per-isoform analysis revealed opposite tail length dynamics among isoforms during the MZT, with one isoform decreasing and another isoform increasing in polyA tail length as the MZT progressed (**Figure 4a,b**, see also **Figure S4a,b**).

**Figure 4.**
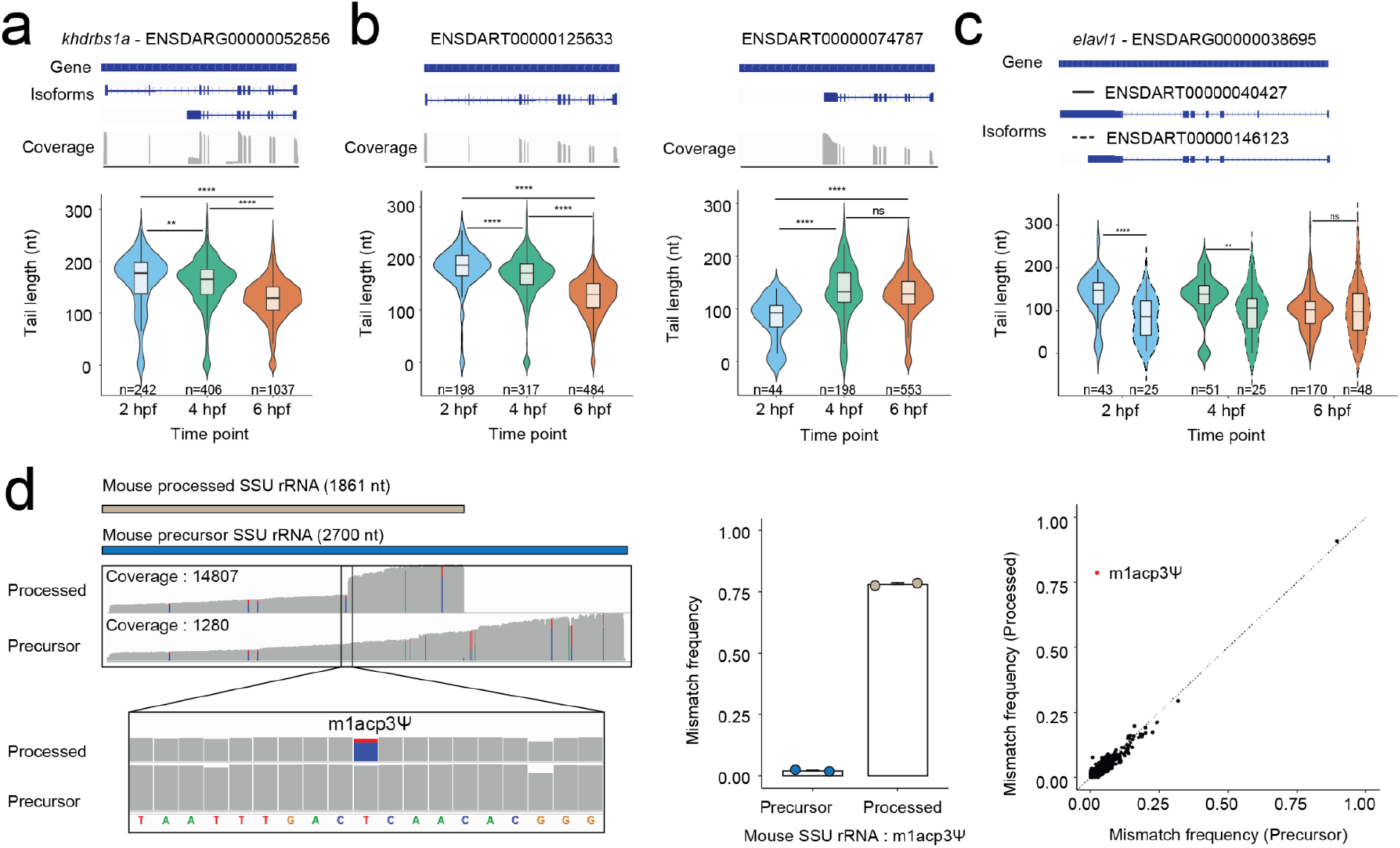
Isoform-specific polyA tail and modification dynamics can be captured using Nano3P-seq. **(a)** Comparison of polyA tail length distributions of reads mapping to *khdrbs1a*, illustrated at the per-gene level, measured at three time points during the zebrafish MZT. Annotations of the gene and two main isoforms are shown at the top of the panels, along with an IGV coverage track of the reads mapping to the gene. **(b)** Comparison of polyA tail length distributions of reads mapping to distinct isoforms of *khdrbs1a* measured at three time points during the zebrafish MZT. Annotations of the isoforms and IGV coverage track of the reads mapping to each isoform are shown at the top of the panels. Only isoforms with more than 10 reads are shown. The number of reads included in the analysis is shown below each violin plot. P-values have been computed using the Kruskal–Wallis test and corrected for multiple testing using the Benjamini–Hochberg method. **(c)** Comparison of polyA tail length distributions of reads mapping to two distinct isoforms (full and dashed outline) of *elavl1* measured at three time points during the zebrafish MZT. Annotations of the gene and two main isoforms are shown at the top of the panels. Only isoforms with more than 10 reads are shown. The number of reads included in the analysis is shown below each violin plot. P-values have been computed using the Kruskal–Wallis test and corrected for multiple testing using the Benjamini–Hochberg procedure (P > 0.05:ns, p ≤ 0.05:*, p ≤ 0.01:**, p ≤ 0.001:***, p ≤ 0.0001:****). **(d)** IGV coverage tracks of reads mapping to mouse processed small subunit (SSU) rRNA (upper track) and precursor SSU rRNA (lower track), including a magnified image at the position known to be modified with m^1^acp^3^Ψ (left panel). Positions with a mismatch frequency lower than 0.1 are shown in gray. In the middle panel, the mismatch frequency values in mouse precursor and processed SSU rRNA at the position known to be modified with m^1^acp^3^Ψ (*n* = 2 biological replicates) are shown. In the right panel, the per-site mismatch frequencies observed in reads mapping to mouse precursor SSU rRNA and mouse processed SSU rRNA are compared, showing that the only outlier is m^1^acp^3^Ψ. See also Figure S4.

Next, we compared isoform-specific polyA tail lengths across isoforms encoded by the same gene and found that 17.3% (± 6.7%) of analyzed genes presented significant differences in their polyA tail lengths across isoforms (**Figure 4c**, see also **Figure S4c and Table S7**). Altogether, these results show that polyA tail length dynamics are not only dependent on the gene and embryogenesis stage, but are also specific to individual transcript isoforms. Moreover, these findings demonstrate that Nano3P-seq can provide transcriptome-wide measurements of the polyadenylation status of diverse biological samples with both single-read and single-isoform resolution.

### Detection of isoform-specific RNA modifications using Nano3P-seq

RNA molecules are decorated with chemical modifications, which are essential for the stability, maturation, fate, and function of the RNA ^31–34^. Some modifications occur in base positions that are involved in Watson-–Crick (WC) base-pairing, causing a disruption during reverse transcription. Consequently, these modifications can be seen as increased “errors’’ and drop-off rates in RNA-seq datasets ^35–38^. One example is the hypermodified base m^1^acp^3^Ψ, which is present in the eukaryotic small subunit (SSU) rRNA ^39^ crucial for the final processing steps of precursor rRNA (pre-rRNA) into mature SSU rRNA ^40,41^.

Therefore, we examined whether Nano3P-seq could capture RNA modification differences across precursor and mature rRNA molecules from distinct maturation stages. To this end, we assigned reads mapping to SSU rRNAs as either “precursor” or “processed” (**Figure 4d,** left panel) and analyzed the sequencing error patterns in the two populations. We observed that the mismatch frequency at the m^1^acp^3^Ψ-modified site was very high in mature rRNAs but not present in pre-rRNAs, suggesting that this modification is only present in mature rRNA populations. The presence of the hypermodification was also accompanied by a drop-off in sequencing coverage at the m^1^acp^3^Ψ-modified site, in agreement with its ability to disrupt the WC base-pairing. Analysis of the “error” signatures along the SSU transcripts showed that this position was the only one position to change between precursor and processed rRNA molecules (**Figure 4d**). These results were also orthogonally confirmed in dRNA-seq datasets, although the difference between pre-rRNA and mature rRNA error patterns were less evident than in Nano3P-seq datasets, likely due to the presence of m^1^Ψ modification (precursor of m^1^acp^3^Ψ) at this site in the pre-rRNAs, which also causes increased “errors” in dRNA-seq datasets, but not in Nano3P-seq datasets (**Figure S4d**). Finally, we noted that the same pattern of m^1^acp^3^Ψ modification was observed in yeast SSU rRNA (**Figure S4e**), in agreement with previous studies ^41^. This analysis could not be performed in the zebrafish Nano3P-seq datasets due to the absence of *de novo* transcription of zygotic rRNAs in early embryo stages ^42^. Altogether, our results demonstrate that Nano3P-seq can identify isoform-specific and/or maturation-dependent RNA modification in the form of altered mismatch frequencies and/or reverse transcription drop-offs.

### Analysis of tail composition using Nano3P-seq

Recent works using TAIL-seq have reported that a number of terminal modifications in polyA tails, such as polyuridine stretches, play a role in mRNA decay ^11,20,43,44^. However, these methods cannot be used to detect tail modifications among the vast majority of tail nucleotides, as Illumina sequencing quality strongly deteriorates in homopolymeric stretches and with increased read length. In contrast, Pacific BioSciences (PacBio)-based approaches such as FLAM-seq can sequence through the entire tail, however, they cannot unambiguously identify 3′ terminal modifications ^45^.

Therefore, we explored whether Nano3P-seq could accurately identify nucleotide composition variations (either internal or terminal) within polyA tails. To this end, we designed synthetic cDNA molecules with polyA tails ending with A (30A), U (1U), UUU (3U), UUUUU (5U), CCCCC (5C), and GGGGG (5G), as well as a polyA tail that contained several internal G nucleotides at fixed positions (IntG) **(Figure 5a**, see also *Methods*). Synthetic molecules were sequenced using Nano3P-seq, and for each read, the “tail” was defined as the set of nucleotides found between the nanopore adapter and the last nucleotide mapped to the cDNA standard reference (**Figure 5b,** see also **Figure S5a**). Nucleotide composition analysis of the last 20 nt of each synthetic tail revealed that Nano3P-seq accurately estimated the non-A base content in the tails (**Figure S5b**) and accurately identified the position in which these non-A bases were found (**Figure 5c**).

**Figure 5.**
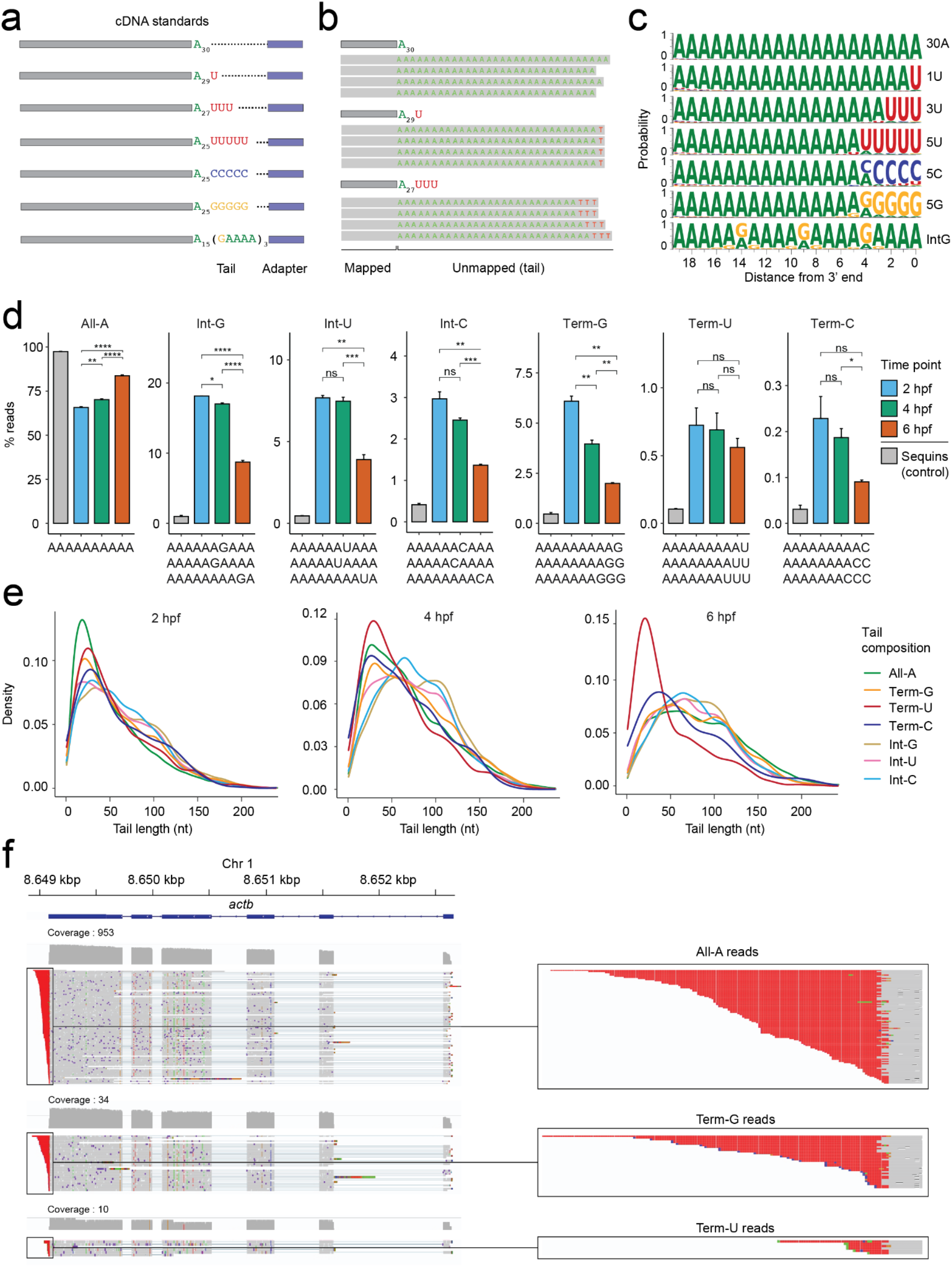
Analysis of tail composition using Nano3P-seq. **(a)** Schematic overview of the standards used to assess the ability of Nano3P-seq to accurately quantify the base content of polyA tails. **(b)** IGV snapshots of nucleotide composition in cDNA standard tails sequenced using Nano3P-seq. Grey regions indicate the mapped part of the reads, whereas colored letters indicate soft-clipped bases (unmapped) which are the base-called tails, after trimming the adapter. **(c)** Probability of base composition (A: green, G: orange, C: blue, U: red) per position in the last 20 nt of the cDNA standard tails. **(d)** Percentage of reads belonging to groups classified based on their polyA tail base composition. Some sequence examples belonging to different groups are illustrated below the bar plots. Samples in this analysis are embryonic mRNAs across zebrafish developmental stages (2, 4, and 6 hpf, shown in blue, green, and red, respectively) in three biological replicates and a control that includes sequin R1 and R2 groups of RNAs (gray). Statistical comparison of means was performed using the Kruskal–Wallis test (*n* = 3 biological replicates, error bars indicate the standard deviation). (P > 0.05:ns, p ≤ 0.05:*, p ≤ 0.01:**, p ≤ 0.001:***, p ≤ 0.0001:****). **(e)** PolyA tail length estimation distributions of mRNA reads belonging to groups classified based on their polyA tail base composition across zebrafish development stages (2, 4, and 6 hpf). **(f)** IGV snapshots of reads mapping to zebrafish *actb* mRNA (left panel). In the right panel, zoomed images of individual reads with different terminal bases (all-A reads: top, Term-G reads: middle, Term-U reads: bottom) are shown.

Next, we analyzed the mRNA tail composition of zebrafish embryos from different developmental stages (2, 4, and 6 hpf). Analysis of tail base abundance revealed that G was the most common non-A base in zebrafish mRNAs and that there was a significant decrease in non-A bases with progression of the MZT (**Figure S6a,b**). As a control, we examined the composition of bases in synthetic molecules (‘sequins’) sequenced with Nano3P-seq and found that the frequency of non-A bases in biological sequences was significantly higher (6-15–fold) than in synthetic molecules (**Figure S6a,** see also **Table S8)**, suggesting that the majority of non-A bases observed in Nano3P-seq biological datasets are not base-calling artifacts.

We then binned the tails based on the composition of the last 30 nt (All-A: contains only A, Int-G: contains internal Gs, Int-U: contains internal Us, Int-C: contains internal Cs, Term-G: contains terminal Gs, Term-U: contains terminal Us, Term-C: contains terminal Cs) (**Figure 5d**, see also **Figure S6c).** Our analysis showed that non-A internal bases were more abundant than non-A terminal bases, with internal Gs (Int-G) being the most abundant type of non-A bases in polyA tails (**Figure 5d,** see also **Table S9**). Moreover, we found that the abundance of transcripts with non-A base tails (internal or terminal) typically decreased with progression of the MZT, except for transcripts with terminal Us, for which the observed decrease was not statistically significant.

Finally, we examined the relationship between tail length and the presence of non-A bases in the tails. Although the median tail length increases during the MZT (**Figure 5e**, see also **Figure 3g**), we found that this increase did not occur for all transcripts, as it depended on tail composition. Specifically, terminally uridylated tails (Term-U) were short (median length = 42, 50, 32 nt at 2, 4, 6 hpf, respectively) regardless of the time point examined (**Figure 5e-f,** see also **Table S10**). An equivalent analysis in mouse Nano3P-seq datasets revealed that Term-U tails were also significantly shorter than the other tail types examined (**Figure S6d**); by contrast, this trend was not observed in yeast Nano3P-seq datasets (**Figure S6e**). Our results are in agreement with previous studies in mouse and human cell lines showing that G and U bases are common in polyA tails and that tails ending with U bases are more frequent in shorter polyA tails ^11,46^.

## Discussion

In the last few years, a variety of NGS-based high-throughput methods have been developed to characterize the 3’ ends of RNA molecules at a transcriptome-wide scale, including methods to reveal polyA tail sites (e.g., 3P-seq ^6^, PAS-seq ^47^, PAT-seq ^48^) and to estimate polyA tail lengths (e.g., PAL-seq ^10^, TAIL-seq ^11^, mTAIL-seq ^49^). However, a major limitation of NGS-based methods is their inability to assign a given polyA tail length to a specific transcript isoform, which causes a loss of isoform-specific tail length information. In addition, NGS-based methods cannot measure tail lengths greater than the read length, thus biasing our view of polyA tail dynamics to those transcripts that display shorter tail lengths.

More recently, novel methods for estimating polyA tail lengths using PacBio long-read sequencing technologies have been developed, such as FLAM-seq ^45^ and PAIso-seq ^46^. In contrast to NGS, these methods can capture isoform–tail relationships; however, they are still affected by PCR amplification and ligation biases, in addition to producing relatively modest outputs in terms of the number of reads ^50–52^. Moreover, PacBio typically requires expensive sequencing instruments that are not widely available. Nanopore dRNA-seq has been proposed as an alternative long-read sequencing technology for studying polyA tail lengths ^12,13^; however, the standard dRNA-seq approach (**Figure 1a**) cannot capture deadenylated RNAs, molecules with non-canonical tailings (e.g., polyuridine), or molecules with polyA tails shorter than 10 nt, thus biasing the view of the transcriptome toward polyadenylated molecules. Customized dRNA-seq methods involving *in vitro* poly(G/I)-tailing have been developed to overcome some of these limitations, but a lack of bioinformatic tools to distinguish polyI and polyA signals limits their applicability to study polyA tail length differences across transcripts in these datasets ^14,15^. In addition, dRNA-seq requires 500 ng of RNA as input, whereas Nano3P-seq requires as little as 50 ng, thus decreasing the required input material by 10-fold. Nano3P-seq addresses the current limitations by offering a simple and robust solution for studying the coding and non-coding transcriptome simultaneously regardless of the presence or absence of polyA tails or 3’ tail composition, without PCR or ligation biases, and with single-read and single-isoform resolution. Moreover, the use of thermostable group II intron reverse transcriptase (TGIRT) in the Nano3P-seq protocol not only maximizes the production of full-length cDNAs, but also ensures the inclusion of RNA molecules that are highly structured and/or modified, which would often not be captured (or their representation would be significantly biased) using standard viral reverse transcriptases ^53,54^.

Nano3P-seq provides quantitative measurements of RNA abundances (**Figure 1d**) and captures diverse RNA biotypes regardless of their tail end composition (**Figure 2d**). We have shown that Nano3P-seq can be applied to diverse species with a wide range of polyA tail lengths (**Figure 3d,e**) and can be used to study the dynamics of polyadenylation (**Figure 3f,g,i**). Specifically, we have demonstrated that Nano3P-seq provides per-read-resolution transcriptome-wide maps of RNA abundance and polyadenylation dynamics during the zebrafish MZT. Our results show that transcripts targeted by miR-430 decay in a deadenylation-dependent manner, whereas those targeted by maternal and zygotic decay have distinct polyA tail length dynamics during the MZT (**Figure 3h,i**). Moreover, we identified isoform-specific regulation of polyadenylation, demonstrating that analyses at the per-gene level are insufficient to capture the dynamics of polyadenylation during the zebrafish MZT (**Figure 4a**–**c**, see also **Figure S4a**–**c**). Overall, we have demonstrated that Nano3P-seq can identify polyadenylation changes across time points, mRNA decay programs, and isoforms, providing mechanistic insights into different gene regulatory programs.

Using Nano3P-seq, we compared the zebrafish transcriptome from both ribodepleted and polyA-selected transcriptomes during the zebrafish MZT. Because the vast majority of cellular RNA is composed of rRNA, transcriptomic studies typically remove a significant portion of rRNA molecules to sequence a wider diversity of RNA biotypes. This removal can be achieved by i) ribodepletion of the sample using biotinylated oligos that are complementary to rRNAs or ii) selective enrichment of polyA(+) transcripts using oligo(dT) beads. Although these two approaches are often used interchangeably, their effects on the transcriptome composition are not equal. Nano3P-seq allows us to compare the effects of these two approaches on both the transcriptome composition and polyA tail length distribution. In terms of its effects on transcriptome composition, we found that ribodepletion captures a larger variety of RNA biotypes compared with polyA-selection, including several non-polyA-tailed RNA biotypes, as expected (**Figure 2f**). However, we did not observe a significant difference in the distribution of mRNA polyA tail lengths between the two methods (**Figure S7a–c**), suggesting that oligo(dT) enrichments do not significantly bias the polyA(+) mRNA populations by preferentially enriching for those with longer polyA tails.

In addition, we performed a comparative analysis of zebrafish polyA tail length, read length, and per-read quality in libraries sequenced using either Nano3P-seq or dRNA-seq. We found that Nano3P-seq captures RNA molecules regardless of their tail ends, resulting in the capture of diverse RNA biotypes (**Figure 1b** and **S1c–g**) including deadenylated mRNA molecules (**Figure S3h** and **S7**) and molecules terminating with non-A bases in their 3’ ends (**Figure 5** and **S5,6**). By contrast, dRNA-seq only captured longer polyadenylated transcripts, as this method relies on the presence of polyA tail lengths greater than 10 nt. Indeed, when comparing the distribution of per-read polyA tail length estimations of mRNAs, we observed that Nano3P-seq captured mRNAs with predicted tail lengths of zero, whereas dRNA-seq only captured reads with longer tails (**Figure S7d,e**). Moreover, we find that Nano3P-seq produces reads with significantly longer lengths than dRNA-seq (**Figure S7f**), whereas the per-read qualities are similar between the two methods (**Figure S7g**), in agreement with previous works ^12^. In terms of sequencing output, the yields of Nano3P-seq runs are similar -or slightly better- to those observed in dRNA-seq runs, producing ∼100-200K reads in Flongle devices and ∼0.5-2M in MinION devices, depending on the RNA input type and quality of the flowcell (**Table S11**).

Finally, we examined the ability of Nano3P-seq to capture tail composition variations in individual RNA molecules during the zebrafish MZT. To ensure that the observed base variations within reads were not caused by base-calling or sequencing artifacts, we also analyzed the tail composition of synthetic sequences (“sequins”), which were used to define the “background” error of our method. We found that 2.4% of the sequin tails contained non-A bases in some form (TermU, TermG, TermC, IntU, IntG, IntC), whereas 35%, 31%, and 16% of the zebrafish reads (at 2, 4, and 6 hpf, respectively) contained non-A bases in their tails (**Table S10**). These results outrule sequencing or base-calling artifacts as a major source of the base composition variation observed in *in vivo* zebrafish samples using Nano3P-seq. Thus, we conclude that Nano3P-seq is a robust approach for characterizing base composition dynamics in short and long tails at per-isoform resolution.

Overall, our work demonstrates that Nano3P-seq can simultaneously capture both non-polyA-tailed and polyA-tailed transcriptomes, making it possible to accurately quantify the RNA abundances and polyA tail lengths at per-read and per-isoform levels, while minimizing the amount of biases introduced during library preparation. These features set Nano3P-seq as a potent, low-cost ^55^ method that can provide mechanistic insights into the regulation of RNA molecules and improve our understanding of mRNA tailing processes and post-transcriptional control.

## Methods

### *In vitro* transcription of RNAs

The synthetic ‘curlcake’ sequences ^56^ (Curlcake 1, 2244 bp and Curlcake 2, 2459 bp) were *in vitro* transcribed using Ampliscribe™ T7-Flash™ Transcription Kit (Lucigen-ASF3507). Curlcake 2 was polyadenylated using *E. coli* polyA Polymerase (NEB-M0276S). polyA-tailed RNAs were purified using RNAClean XP beads. The quality of the *in vitro* transcribed (IVT) products as well as the addition of polyA tail to the synthetic constructs was assessed using Agilent 4200 Tapestation **(Figure S1a**). Concentration was determined using Qubit Fluorometric Quantitation. Purity of the IVT products was measured with NanoDrop 2000 Spectrophotometer.

### Poly(A) Tail-Length Assay and Sanger sequencing

Poly(A) Tail-Length Assay Kit (Thermo Fisher, 764551KT) was used following the manufacturer’s instructions. PCR products corresponding to the tail- and gene-specific primer combinations of mouse 16S rRNA (universal forward primer: 5’- GGTCGGTTTCTATCTATTTACGATTTCTC -3’, gene-specific reverse primer: 5’- TTCTCTAGGTTAGAGGGTGTACGTATAT -3’) and human ACTB (assay control, primer composition not disclosed by manufacturer) were loaded on a 2.5% agarose gel (Lonza, 50010) and stained with GelRed® (Merck, SCT123). Product sizes determined using the GeneRuler 50bp DNA ladder (Thermo Scientific, SM0371). Subsequently, tail- and gene-specific PCR products of mouse 16S rRNA were purified by gel-elution (Cytiva Life Sciences™, 28903470) and sent for Sanger sequencing with the shared forward primer (5’- GGTCGGTTTCTATCTATTTACGATTTCTC -3’). Resulting chromatograms were analyzed using SnapGene (v.6.0.2). After confirming alignment to the reference sequence, the unclipped chromatograms were used to visualize 3’-ends.

### Yeast culturing and total RNA extraction

*Saccharomyces cerevisiae* (strain BY4741) was grown at 30°C in standard YPD medium (1% yeast extract, 2% Bacto Peptone and 2% dextrose). Cells were then quickly transferred into 50 mL pre-chilled falcon tubes, and centrifuged for 5 minutes at 3,000 g in a 4°C pre-chilled centrifuge. Supernatant was discarded, and cells were flash frozen. Flash frozen pellets were resuspended in 700 µL Trizol with 350 µL acid washed and autoclaved glass beads (425-600 µm, Sigma G8772). The cells were disrupted using a vortex on top speed for 7 cycles of 15 seconds (the samples were chilled on ice for 30 seconds between cycles). Afterwards, the samples were incubated at room temperature for 5 minutes and 200 µL chloroform was added. After briefly vortexing the suspension, the samples were incubated for 5 minutes at room temperature. Then they were centrifuged at 14,000 g for 15 minutes at 4°C and the upper aqueous phase was transferred to a new tube. RNA was precipitated with 2X volume Molecular Grade Absolute ethanol and 0.1X volume Sodium Acetate (3M, pH 5.2). The samples were then incubated for 1 hour at -20°C and centrifuged at 14,000 g for 15 minutes at 4°C. The pellet was then washed with 70% ethanol and resuspended with nuclease-free water after air drying for 5 minutes on the benchtop. Purity of the total RNA was measured with the NanoDrop 2000 Spectrophotometer. Total RNA was then treated with Turbo DNase (Thermo, #AM2238) (2 ul enzyme for 50 ul reaction of 200 ng/ul RNA) at 37°C for 15 minutes, with a subsequent RNAClean XP bead cleanup.

### Mice breeding

Experiments were performed with male mice aged between 8 and 10 weeks. All mice were euthanized using CO_2_ and tissues were snap frozen in liquid nitrogen. Animals were kept on a 12:12h light:dark cycle and provided with water and food *ad libitum*.

### RNA isolation from mouse brain

In order to isolate nuclear/mitochondrial-enriched RNA from the mouse (*Mus musculus*) brain, we followed previously published protocols ^57^ with minor changes. A quarter of a C57BL6/J mouse brain was used for this protocol, and all samples and reagents were kept on ice during the protocol. Brain tissue was mined with a razor blade into smaller pieces. Cold Nuclei EZ Lysis Buffer (0.01 M Tris-Cl,pH7.5, 0.06M KCl, 0.001M EDTA, 0.5% NP40, 1X Protease Inhibitor (complete™ Protease Inhibitor Cocktail Tablets, #11697498001, Roche)) was added to the tissue in 1.5 mL eppendorf tube. The sample was homogenized using a 1 mL dounce (stroking ∼10-20 times), and the homogenate was transferred into a 2mL eppendorf tube. 1 mL of cold Nuclei EZ Lysis Buffer was added and mixed, followed by 4 minutes incubation on ice. During the incubation, the sample was gently mixed a couple of times using a pipette. Homogenate was filtered using a 70 um strainer mesh, and the flowthrough was collected in a polystyrene round-bottom FACS tube and subsequently transferred into a new 2 mL tube. The sample was centrifuged at 500g for 5 minutes at 4°C and the supernatant was removed. The nuclei/mitochondria enriched sample was resuspended in another 1.5 mL EZ Lysis buffer and incubated for 5 minutes on ice. The sample was centrifuged at 500 g for 5 minutes 4°C and the supernatant was discarded (cytoplasm). 500 uL Nuclei Wash and Resuspension Buffer (1M PBS, 1%BSA, SUPERase In RNase Inhibitor (Thermo Fisher, AM2694)) was added to the sample and incubated for 5 minutes without resuspending to allow buffer interchange. After incubation, 1 mL of Nuclei Wash and Resuspension Buffer was added and the sample was resuspended. The sample was centrifuged at 500g for 5 minutes at 4°C. The supernatant was removed and only ∼50 ul was left. Using 1.4 mL Nuclei Wash and Resuspension Buffer, the sample was resuspended and transferred to a 1.5 mL eppendorf tube. The last washing step was repeated and the pellet was resuspended in 300 uL Nuclei Wash and Resuspension Buffer. RNA was extracted using Trizol.

### Zebrafish breeding

Wild-type zebrafish (*Danio rerio*) embryos were obtained through natural mating of the TU-AB strain of mixed ages (5–18 months). Mating pairs were randomly chosen from a pool of 60 males and 60 females allocated for each day of the month. Embryos and adult fish were maintained at 28 °C.

### Zebrafish total RNA extraction and polyA selection

For RNA samples, 25 embryos per developmental stage and per replicate were collected and flash frozen in liquid nitrogen. Frozen embryos were thawed and lysed in 1 mL TRIzol (Life Technologies) and total RNA was extracted using the manufacturer’s protocol. Total RNA concentration was calculated by nanodrop.

For polyA-selected RNA samples, polyadenylated RNAs were isolated with oligo (dT) magnetic beads (New England BioLabs, S1419S) according to the manufacturer’s protocol and eluted in 30 µL prior to nanodrop quantification.

### Zebrafish total RNA ribodepletion

Ribodepletion was performed on zebrafish total RNA using riboPOOL oligos (siTOOLs, cat #055) following the manufacturer’s protocol. Briefly, 5 ug total RNA in 14 uL was mixed with 1 uL resuspended riboPOOL oligos, 5 uL hybridization buffer and 0.5 uL SUPERase•In RNase Inhibitor (Thermo Fisher, AM2694). The mix was incubated for 10 minutes at 68°C, followed by a slow cool down (3°C/min) to 37°C for hybridization. In the meantime, Dynabeads MyOne Streptavidin C1 (Thermo Fisher, 65001) beads were resuspended by carefully vortexing at medium speed. 80 uL of Dynabeads bead resuspension (10 mg/mL) was transferred into a tube, which then was placed on a magnetic rack. After aspirating the supernatant, 100 uL of bead resuspension buffer (0.1 M NaOH, 0.05 M NaCl) was added to the sample and beads were resuspended in this buffer by agitating the tube. Sample was placed on a magnet and the supernatant was aspirated. This step was performed twice. Beads were then resuspended in 100 uL of bead wash buffer (0.1 M NaCl) and placed on magnet in order to aspirate the supernatant. Beads were then resuspended in a 160 uL depletion buffer (5 mM Tris-HCl pH 7.5, 0.5 mM EDTA, 1 M NaCl). This suspension was then divided into two tubes of 80 uL, which will be used consecutively. 20 uL of hybridized riboPOOL and total RNA was briefly centrifuged to spin down droplets and it was pipetted into the tube containing 80 uL of beads in depletion buffer (10 mM Tris-HCl pH 7.5, 1 mM EDTA, 2 M NaCl). The tube containing the mix was agitated to resuspend the solution well. Then the mix was incubated at 37°C for 15 minutes, followed by a 50°C incubation for 5 minutes. Immediately before use, the second tube containing 80 ul of beads was placed on a magnetic rack and the supernatant was aspirated. After the incubation at 50°C, the first depletion reaction was placed on a magnet and the supernatant was transferred into the tube containing the other set of beads. The mix was incubated again at 37°C for 15 minutes, followed by a 50°C incubation for 5 minutes. After briefly spinning down the droplets, the mix was placed on a magnet for 2 minutes. The supernatant was transferred into a different tube and cleaned up using RNA Clean & Concentrator-5 (Zymo, R1013).

### Nano3P-Seq library preparation

The protocol is based on the direct cDNA Sequencing ONT protocol (DCB_9091_v109_revC_04Feb2019), with several modifications to be able to perform TGIRT template switching. Before starting the library preparation, 1 µL of 100 µM R_RNA (Oligo: 5’ rGrArArGrArUrArGrArGrCrGrArCrArGrGrCrArArGrUrGrArUrCrGrGrArArG/3SpC3/ 3’) and 1 µL of 100 µM D_DNA (5’ /5Phos/CTTCCGATCACTTGCCTGTCGCTCTATCTTCN 3’) were mixed with 1 µL 0.1 M Tris pH 7.5, 1 µL 0.5 M NaCl, 0.5 ul RNAse Inhibitor Murine (NEB, M0314S) and 5.5 ul RNase-free water. The mix was incubated at 94°C for 1 minute and the temperature was ramped down to 25°C (- 0.1°C/s) in order to pre-anneal the oligos. Then, 50-100 ng RNA was mixed with 1 µL pre-annealed R_RNA+D_DNA oligo, 1uL 100 mM DTT, 4 µL 5X TGIRT Buffer (2.25 M NaCl, 25 mM MgCl2, 100 mM Tris-HCl, pH 7.5), 1 µL RNasin® Ribonuclease Inhibitor (Promega, N2511), 1-2 µL TGIRT (InGex) and nuclease-free water up to 19 µL. We should note that if 50 ng are used as input, only 1 µL of TGIRT is needed, whereas if 100 ng is used as input, 2µL of TGIRT enzyme is needed. The reverse-transcription mix was initially incubated at RT for 30 minutes before adding 1 µL 10 mM dNTP mix. Then the mix was incubated at 60°C for 60 minutes and inactivated by heating at 75°C for 15 minutes before moving to ice. RNAse Cocktail (Thermo Scientific, AM2286) was added to the mix in order to digest the RNA, and the mix was incubated at 37°C for 10 minutes. The reaction was then cleaned up using 0.8X AMPure XP Beads (Agencourt, A63881). In order to be able to ligate the sequencing adapters to the first cDNA strand, 1 µL 100 µM CompA_DNA (5’ GAAGATAGAGCGACAGGCAAGTGATCGGAAGA 3’) was annealed to the 15 µL cDNA in a tube with 2.25 µL 0.1 M Tris pH 7.5, 2.25 µL 0.5 M NaCl and 2 µL nuclease-free water. The mix was incubated at 94°C for 1 minute and the temperature was ramped down to 25 °C (-0.1°C/s) in order to anneal the complementary to the first strand cDNA. Then, 22.5 µL first strand cDNA was mixed with 5 µL Adapter Mix (AMX), 22.5 µL Rnase-free water and 50 µL Blunt/TA Ligase Mix (NEB, M0367S) and incubated in room temperature for 10 minutes. The reaction was cleaned up using 0.8X AMPure XP beads, using ABB Buffer for washing. The sample was then eluted in Elution Buffer (EB) and mixed with Sequencing Buffer (SQB) and Loading Beads (LB) prior to loading onto a primed R9.4.1 flowcell. Libraries were run on either Flongle or MinION flowcells with MinKNOW acquisition software version v.3.5.5. A detailed step-by-step Nano3P-seq protocol is provided as an additional file (**File S1**).

### Annealing based direct cDNA-Sequencing library preparation with TGIRT

Some adjustments were made to the original Direct cDNA-Sequencing ONT protocol (SQK-DCS109), in order to be able to use TGIRT (InGex) as reverse transcription enzyme for nanopore sequencing, as this enzyme does not produce CCC overhang that is typically exploited by the direct cDNA sequencing library preparation protocol (**Figure 1a**). Briefly, 1 µL of 100 µM Reverse transcription primer VNP (5’ /5Phos/ACTTGCCTGTCGCTCTATCTTCTTTTTTTTTTTTTTTTTTTTVN 3’) and 1 µL of 100 µM of *in-house* designed complementary oligo (CompA: 5’ GAAGATAGAGCGACAGGCAAGTA 3’) were mixed with 1 µL 0.2 M Tris pH 7.5, 1 µL 1 M NaCl and 16ul RNase-free water. The mix was incubated at 94°C for 1 minute and the temperature was ramped down to 25°C (-0.1°C/s) in order to pre-anneal the oligos. Then, 50 ng polyA-tailed RNA was mixed with 1 µL pre-annealed VNP+CompA,, 1uL 100 mM DTT, 4 µL 5X TGIRT Buffer (2.25 M NaCl, 25 mM MgCl2, 100 mM Tris-HCl, pH 7.5), 1 µL RNasin® Ribonuclease Inhibitor (Promega, N2511), 1 µL TGIRT and nuclease-free water up to 19 µL. The reverse-transcription mix was initially incubated at RT for 30 minutes before adding 1 µL 10 mM dNTP mix. Then the mix was incubated at 60°C for 60 minutes and inactivated by heating at 75°C for 15 minutes before moving on to ice. Furthermore, RNAse Cocktail (Thermo Scientific, AM2286) was added to the mix in order to digest the RNA and the mix was incubated at 37°C for 10 minutes. Then the reaction was cleaned up using 0.8X AMPure XP Beads (Agencourt, A63881). In order to be able to ligate the sequencing adapters the the first strand, 1 µL 100 µM CompA was again annealed to the 15 µL cDNA in a tube with 2.25 µL 0.1 M Tris pH 7.5, 2.25 µL 0.5 M NaCl and 2 µL nuclease-free water. The mix was incubated at 94°C for 1 minute and the temperature was ramped down to 25 °C (-0.1°C/s) in order to anneal the complementary to the first strand cDNA. Furthermore, 22.5 µL first strand cDNA was mixed with 5 µL Adapter Mix (AMX), 22.5 uL Rnase-free water and 50 µL Blunt/TA Ligase Mix (NEB, M0367S) and incubated in room temperature for 10 minutes. The reaction was cleaned up using 0.8X AMPure XP beads, using ABB Buffer for washing. The sample was then eluted in Elution Buffer (EB) and mixed with Sequencing Buffer (SQB) and Loading Beads (LB) prior to loading onto a primed R9.4.1 flowcell and ran on a MinION sequencer with MinKNOW acquisition software version v.3.5.5.

### Synthetic cDNA standards

A total of 12 synthetic cDNA standards were synthesized as ultramers by IDT (Integrated DNA Technologies) to assess the tail length estimation and tail composition quantification accuracy of Nano3P-seq:

Synthetic cDNA standards designed to assess accuracy in tail length estimation:

cDNA_pA_standard_0:

/5Phos/CTTCCGATCACTTGCCTGTCGCTCTATCTTCGTAAATAGAAATAGACTAGCTCCACTTTTAAGAATTATTTATGCAATTAAATACATGGGTGACCAAAAGAGCGGGCGGATACACGCGTCACCACAAGCAGAATAAAAGGTAAACCTGAAATTGTTTTAACATAAAATGAAAAATGCTTGTTTGCAACCCTATATAGAA

cDNA_pA_standard_15:

/5Phos/CTTCCGATCACTTGCCTGTCGCTCTATCTTCTTTTTTTTTTTTTTTGTAAATAGAAATAGACTAGCTCCACTTTTAAGAATTATTTATGCAATTAAATACATGGGTGACCAAAAGAGCGGGCGGATACACGCGTCACCACAAGCAGAATAAAAGGTAAACCTGAAATTGTTTTAACATAAAATGAAAAATGCT TGTTTG

cDNA_pA_standard_30:

/5Phos/CTTCCGATCACTTGCCTGTCGCTCTATCTTCTTTTTTTTTTTTTTTTTTTTTTTTTTTTTTGTAAATAGAAATAGACTAGCTCCACTTTTAAGAATTATTTATGCAATTAAATACATGGGTGACCAAAAGAGCGGGCGGATACACGCGTCACCACAAGCAGAATAAAAGGTAAACCTGAAATTGTTTTAACATAA AATG

cDNA_pA_standard_60:

/5Phos/CTTCCGATCACTTGCCTGTCGCTCTATCTTCTTTTTTTTTTTTTTTTTTTTTTTTTTTTTTTTTTTTTTTTTTTTTTTTTTTTTTTTTTTTGTAAATAGAAATAGACTAGCTCCACTTTTAAGAATTATTTATGCAATTAAATACATGGGTGACCAAAAGAGCGGGCGGATACACGCGTCACCACAAGCAGAATAAA AG

cDNA_pA_standard_90:

/5Phos/CTTCCGATCACTTGCCTGTCGCTCTATCTTCTTTTTTTTTTTTTTTTTTTTTTTTTTTTTTTTTTTTTTTTTTTTTTTTTTTTTTTTTTTTTTTTTTTTTTTTTTTTTTTTTTTTTTTTTTGTAAATAGAAATAGACTAGCTCCACTTTTAAGAATTATTTATGCAATTAAATACATGGGTGACCAAAAGAGCGGGCGG

cDNA_pA_standard_120:

/5Phos/CTTCCGATCACTTGCCTGTCGCTCTATCTTCTTTTTTTTTTTTTTTTTTTTTTTTTTTTTTTTTTTTTTTTTTTTTTTTTTTTTTTTTTTTTTTTTTTTTTTTTTTTTTTTTTTTTTTTTTTTTTTTTTTTTTTTTTTTTTTTTTTTTTTTGTAAATAGAAATAGACTAGCTCCACTTTTAAGAATTATTTATGCAATT

Synthetic cDNA standards designed to assess accuracy in tail composition analyses:

cDNA_p29A_pU1_standard_30:

/5Phos/CTTCCGATCACTTGCCTGTCGCTCTATCTTCATTTTTTTTTTTTTTTTTTTTTTTTTTTTTGTAAATAGAAATAGACTAGCTCCACTTTTAAGAATTATTTATGCAATTAAATACATGGGTGACCAAAAGAGCGGGCGGATACACGCGTCACCACAAGCAGAATAAAAGGTAAACCTGAAATTGTTTTAACATAA AATG

cDNA_p27A_pU3_standard_30:

/5Phos/CTTCCGATCACTTGCCTGTCGCTCTATCTTCAAATTTTTTTTTTTTTTTTTTTTTTTTTTTGTAAATAGAAATAGACTAGCTCCACTTTTAAGAATTATTTATGCAATTAAATACATGGGTGACCAAAAGAGCGGGCGGATACACGCGTCACCACAAGCAGAATAAAAGGTAAACCTGAAATTGTTTTAACATAA AATG

cDNA_p25A_pU5_standard_30:

/5Phos/CTTCCGATCACTTGCCTGTCGCTCTATCTTCAAAAATTTTTTTTTTTTTTTTTTTTTTTTTGTAAATAGAAATAGACTAGCTCCACTTTTAAGAATTATTTATGCAATTAAATACATGGGTGACCAAAAGAGCGGGCGGATACACGCGTCACCACAAGCAGAATAAAAGGTAAACCTGAAATTGTTTTAACATA AAATG

cDNA_p25A_pC5_standard_30:

/5Phos/CTTCCGATCACTTGCCTGTCGCTCTATCTTCGGGGGTTTTTTTTTTTTTTTTTTTTTTTTTGTAAATAGAAATAGACTAGCTCCACTTTTAAGAATTATTTATGCAATTAAATACATGGGTGACCAAAAGAGCGGGCGGATACACGCGTCACCACAAGCAGAATAAAAGGTAAACCTGAAATTGTTTTAACATA AAATG

cDNA_p25A_pG5_standard_30:

/5Phos/CTTCCGATCACTTGCCTGTCGCTCTATCTTCCCCCCTTTTTTTTTTTTTTTTTTTTTTTTTGTAAATAGAAATAGACTAGCTCCACTTTTAAGAATTATTTATGCAATTAAATACATGGGTGACCAAAAGAGCGGGCGGATACACGCGTCACCACAAGCAGAATAAAAGGTAAACCTGAAATTGTTTTAACATA AAATG

cDNA_pA_internalG_standard_30:

/5Phos/CTTCCGATCACTTGCCTGTCGCTCTATCTTCTTTTCTTTTCTTTTCTTTTTTTTTTTTTTTGTAAATAGAAATAGACTAGCTCCACTTTTAAGAATTATTTATGCAATTAAATACATGGGTGACCAAAAGAGCGGGCGGATACACGCGTCACCACAAGCAGAATAAAAGGTAAACCTGAAATTGTTTTAACATA AAATG

### Sequencing and analysis of dRNA-seq datasets

Direct RNA sequencing (dRNA-seq) library preparations were prepared following manufacturer’s recommendations, and were sequenced in an R 9.4.1 MinION flowcell using a GridION sequencing device. For sequins, reads were base-called using stand-alone Guppy version 3.0.3 with default parameters and then the base-called reads were mapped to sequin sequences ^18^ with minimap2 with - ax splice -k14 -uf --MD parameters ^58^. For zebrafish dRNA-seq samples, reads were base-called with Guppy version 4.0. Base-called reads were first mapped to maternal and somatic zebrafish ribosomal RNA sequences taken from ^42^ and then to the genome (GRCz11) with minimap2 ^58^ with -ax splice -k14 -uf --MD parameters. Mapped reads were intersected with ENSEMBL version 103 annotation (Danio_rerio.GRCz11.103.2.gtf) using bedtools intersect option ^59^.

### Analysis of Nano3P-seq datasets

All the Nano3P-seq runs were basecalled and demultiplexed using stand-alone Guppy version 6.0.1 with default parameters. All runs were mapped using minimap2 ^58^ with the following parameters: -ax splice -k14 -uf --MD (when mapping to genome) and -ax map-ont -k14 –MD (when mapping to transcriptome). For the synthetic constructs (curlcakes), base-called reads were mapped to Curlcake 1 and 2 sequences ^56^, and mapped reads were then intersected with the annotations of Curlcake 1 and 2 sequences to filter out the incomplete reads using bedtools. For yeast total RNA, we first mapped the base-called reads to *S. cerevisiae* ribosomal RNAs (25s, 18s, 5s, 5.8s) and then mapped the rest of the reads to the *S. cerevisiae* genome (SacCer3). Mapped reads were then intersected with SacCer64 annotation exon ends, to filter out incomplete reads. For nuclear/mitochondrial enriched mouse brain RNA spiked in with sequins ^18^, we first mapped the base-called reads to *M. musculus* ribosomal RNAs and then mapped the rest of the rest of the reads to the *M. musculus* genome (GRCm38), supplemented with sequin chromosome (chrIS). Mapped reads were then intersected with ENSEMBL version 102 annotation (Mus_musculus.GRCm38.102.gtf) and sequin annotation (RNAsequins.v2.2.gtf) exon ends, in order to filter the incomplete reads. For zebrafish RNA, we first mapped the base-called reads to ribosomal RNAs and then mapped the rest of the reads to the genome (GRCz11). Mapped read starts were then intersected with ENSEMBL version 103 annotation (Danio_rerio.GRCz11.103.2.gtf) exon ends, to filter the incomplete reads. For assignment of reads to isoforms, IsoQuant package was used (https://github.com/ablab/IsoQuant) with Danio_rerio.GRCz11.103.2.gtf annotation using the following parameters : --genedb gtf_file --complete_genedb --bam bam_file --data_type nanopore -o output. A complete step-by-step command line of the bioinformatic analysis done on Nano3P-seq datasets can be found in the GitHub repository https://github.com/novoalab/Nano3P_Seq.

### Estimation of polyA tail lengths

For direct RNA sequencing reads, polyA tail length estimation was performed using *NanoTail*, a module from *Master of Pores* ^60^, a nextflow workflow for the analysis of direct RNA datasets, which uses internally Nanopolish v0.11.1 ^26^. In *NanoTail*, all reads stored in the fastq files are first indexed with *nanopolish index* using default parameters, and the function *nanopolish polya* is used to perform polyA tail length estimations.

For Nano3P-seq reads, polyA tail length estimation was performed using the Nano3P-seq version of *tailfindR* (https://github.com/adnaniazi/tailfindr/tree/nano3p-seq). All code used to estimate polyA tail lengths and post-process Nano3P-seq data can be found at https://github.com/novoalab/Nano3P_Seq.

### Analysis of tail composition

Base-called reads mapped to the zebrafish genome and cDNA standards for tail composition quantification were first trimmed using the Porechop tool (https://github.com/rrwick/Porechop) with the following parameters; –extra_end_trim 0 –end_threshold 50, in order to remove the adapter sequences. Because Porechop sometimes failed at removal of the adapter sequences, only reads containing more than 80% A bases in their tail composition were kept for downstream analyses, thus ensuring that untrimmed reads are not included in downstream analyses (see **Figure S8**). Finally, an in-house python script was used to extract the tail FASTA sequences of the tail regions from trimmed reads, which has been made available in GitHub (https://github.com/novoalab/Nano3P_Seq/blob/master/soft_clipped_content.py).

### Animal Ethics

Fish lines were maintained according to the International Association for Assessment and Accreditation of Laboratory Animal Care research guidelines, and protocols were approved by the Yale University Institutional Animal Care and Use Committee (IACUC). Mice maintenance was approved by the Garvan/St Vincent’s Hospital Animal Ethics Committee, in accordance with the guidelines of the Australian Code of Practice for the Care and Use of Animals for Scientific Purposes (Project No. 16/14 and 16/26). All animals were entered into the study in a randomized order and operators were blinded to genotype and treatments.

## Data availability

Base-called FAST5 reads from Nano3P-seq and dRNA-seq libraries have been made publicly available in ENA, under accession code PRJEB46978. PolyA tail lengths using PAL-Seq were obtained from GEO with the accession code GSE52809 ^10^. All sequencing runs included in this work are listed in **Table S11**.

## Code availability

All scripts and code used in this work have been made available in GitHub: https://github.com/novoalab/Nano3P_Seq. The code for analyzing Nano3P-seq polyA tail lengths using tailfindR is available on GitHub (https://github.com/adnaniazi/tailfindr/tree/nano3p-seq) and is included as **File S2.**

## Acknowledgements

We thank all the members of the Novoa lab for their valuable insights and discussion. We thank Dr. Tim Mercer for providing us with the sequins used as spike-ins in Nano3P-seq zebrafish runs. OB is supported by a UNSW International PhD fellowship and Australian Government Research Training Program Scholarship. GD is part of the ROPES ITN which received funding from the European Union’s Horizon 2020 research and innovation programme under the Marie Sklodowska-Curie grant agreement no. 956810. AD-T is supported by an FPI-SO fellowship (PRE2019-088498). This work was supported by the Australian Research Council (DP180103571 to EMN) and the Spanish Ministry of Economy, Industry and Competitiveness (MEIC) (PGC2018-098152-A-100 to EMN). We acknowledge the support of the MEIC to the EMBL partnership, Centro de Excelencia Severo Ochoa and CERCA Programme/Generalitat de Catalunya.

## Author contributions

OB performed all the wet lab experiments. OB analyzed the data, together with EMN. GD performed qPCR experiments and G/I-based polyA tail length experiments to orthogonally validate Nano3P-seq predictions. HL contributed code for the analysis of polyA tail lengths and nucleotide content. AD-T processed and analyzed the direct RNA sequencing zebrafish data. AMN and EV adapted tailfindR code to accommodate the adapters that are used in Nano3P-seq libraries. CK, AJG and JDB provided the zebrafish total and polyA-selected RNA samples used in this study. EMN conceived the project. EMN supervised the work, with the assistance of JSM. OB and EMN wrote the paper, with contributions from all authors.

