## Supplementary Figures for "Nano3P-seq: transcriptome-wide analysis of gene expression and tail dynamics using end-capture nanopore cDNA sequencing"

**Figure S1. Nano3P-seq captures non-polyA-tailed and polyA-tailed RNAs (a)** Tapestation profiles of synthetic RNAs (‘curlcakes’) after being *in-vitro* transcribed and polyA tailed (pA). **(b)** Tapestation profiles of the input RNA (curlcake mix) for reverse-transcription and cDNA produced after annealing based or template-switching based (Nano3P-seq) reverse-transcription. **(c)** IGV snapshots of synthetic RNAs (Curlcake1 and Curlcake2, see *Methods*) illustrating that Nano3P-seq captures both non-polyadenylated (left) and polyadenylated (right) RNAs. The PolyA tail region is shown in green. **(d)** Pie chart showing the abundance of different RNA types in Nano3P-seq of mouse nuclear/mitochondria enriched RNA. **(e)** IGV snapshot of reads mapping to *Aldoc* gene with polyA tail shown in green. **(f)** IGV snapshot of reads mapping to *Rps3* and *Snord15b* genes. PolyA tail can be seen in green on the reads mapping to *Rps3* mRNA, while it can’t be seen in *Snord15b* snoRNA. **(g)** IGV snapshot of reads mapping to *Rn7sk* miscRNA, which are not expected to contain polyA tails. **(h)** Scatter plot of the log transformed concentrations (Attomoles/uL) and read counts of sequin transcripts (Pearson R : 0.89, Slope: 0.92). Each dot represents a sequin transcript. **(i)** Scatter plot of the replicability of the log (read counts) of synthetic sequins using Nano3P-seq, both at per-gene level (left panel, Pearson’s R: 0.99) as well as per-transcript level (right panel, Pearson’s R: 0.98).

**Figure S1**

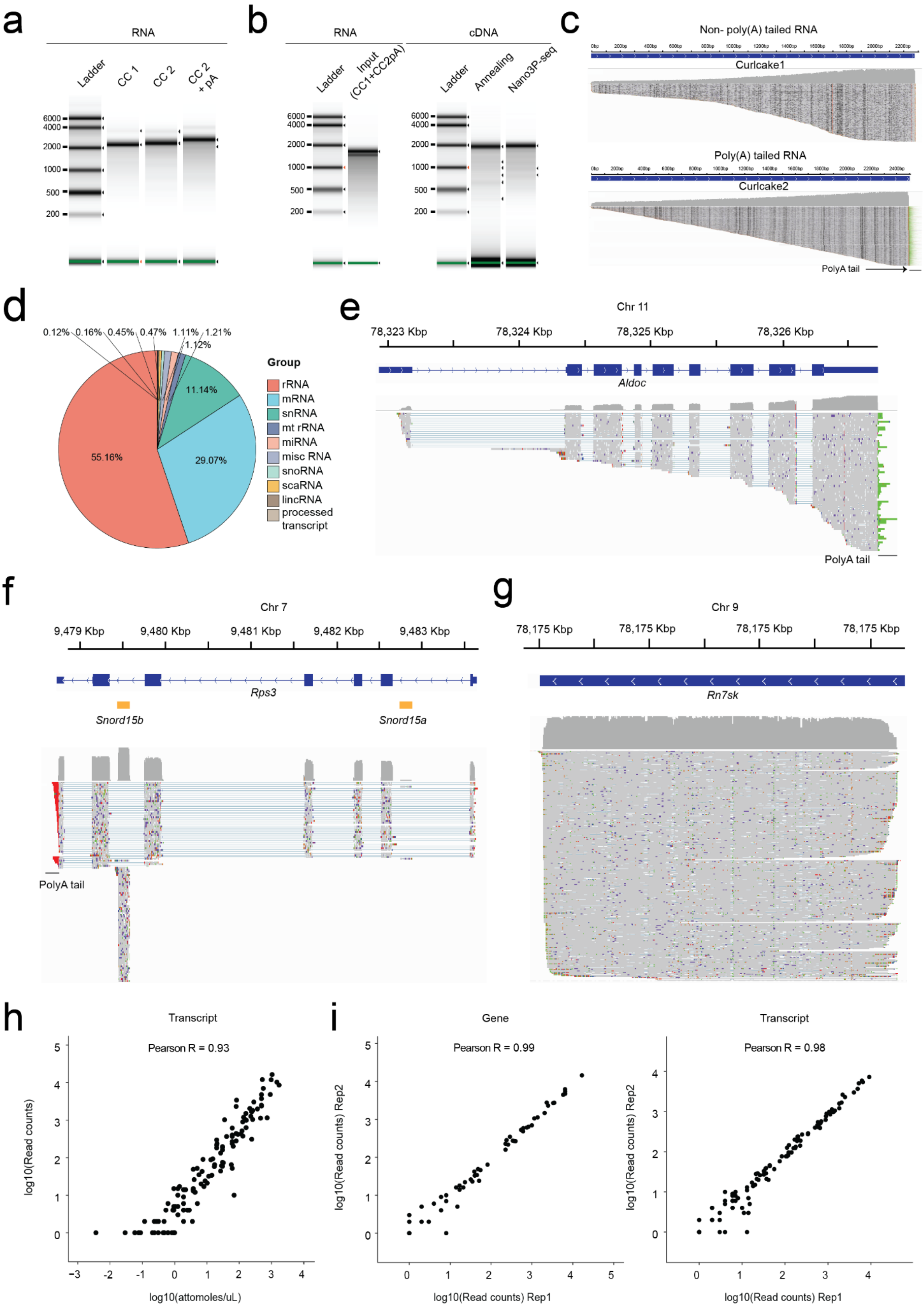
a
RNA
Ladder
CC 1
CC 2
CC 2 + pA
6000
4000
2000
1000
500
200
b
RNA
cDNA
Ladder
Input (CC1+CC2pA)
Annealing
Nano3P-seq
c
Non- poly(A) tailed RNA
Curlcake1
Poly(A) tailed RNA
Curlcake2
PolyA tail
d
0.12%
0.16%
0.45%
0.47%
1.11%
1.21%
1.12%
11.14%
29.07%
55.16%
Group
rRNA
mRNA
snRNA
mt rRNA
miRNA
misc RNA
snoRNA
scaRNA
lincRNA
processed transcript
e
Chr 11
78,323 Kbp
78,324 Kbp
78,325 Kbp
78,326 Kbp
Aldoc
PolyA tail
f
Chr 7
9,479 Kbp
9,480 Kbp
9,481 Kbp
9,482 Kbp
9,483 Kbp
Rps3
Snord15b
Snord15a
PolyA tail
g
Chr 9
78,175 Kbp
78,175 Kbp
78,175 Kbp
78,175 Kbp
Rn7sk
h
Transcript
Pearson R = 0.93
log10(Read counts)
log10(attomoles/uL)
i
Gene
Pearson R = 0.99
log10(Read counts) Rep2
log10(Read counts) Rep1
Transcript
Pearson R = 0.98
log10(Read counts) Rep2
log10(Read counts) Rep1

**Figure S2. Analysis of abundances and polyA tails in mitochondrial rRNAs. (a)** Tapestation profiles of RNAs from zebrafish embryos collected in three different time points during the MZT (2,4,6 hours post-fertilization). Profiles include total RNA, ribodepleted and polyA selected RNA. **(b)** Tapestation profiles of the reverse-transcription products of ribodepleted (left) and polyA selected (right) samples from zebrafish embryos collected in three different time points during the MZT. **(c)** Scatterplots depicting the correlation of mRNA RPM (Read per million) levels biological replicates in three different time points during the MZT. **(d)** Pearson correlation matrix illustrating the similarity between biological replicates and different time points during the MZT. **(e)** Heatmap of log10(RPM) values of micro-RNAs in three different time points during the MZT in three biological replicates**. (f)** Percentage of reads mapping to 12s and 16s mitochondrial rRNAs in two different methods: Nano3P-seq of ribodepleted and polyA selected samples and dRNA-seq of polyA selected samples from zebrafish embryos at 4 hours post-fertilization. **(g)** IGV snapshot of reads mapping to zebrafish 16s mitochondrial rRNA, where reads have been grouped as non-polyA tailed and polyA tailed based on their predicted polyA tail length. PolyA tail region is shown with an arrow and colored green. **(h)** IGV snapshot of reads mapping to mouse 16s mitochondrial rRNA, where reads have been grouped as non-polyA tailed and polyA tailed based on their predicted polyA tail length. PolyA tail region is shown with an arrow and colored green. **(i)** Fold change of the polyA tailed 16s mitochondrial rRNA amount to the total 16s mitochondrial rRNA amount measured by qPCR and Nano3P-seq. **(j)** Outline of PolyA Tail-Length Assay Kit (Thermo, #764551KT) (left panel). Agarose gel electrophoresis image of the PCR products of the PolyA Tail-Length Assay Kit illustrating bands in the tail-specific PCR of *ACTB* control and both tail-specific and gene-specific PCR of mouse 16s mitochondrial rRNA (middle panel). Sanger sequencing result of the PCR product extracted from the agarose gel showing the presence of polyA tail after the reference end indicated by dashed line (right panel).

**Figure S2**

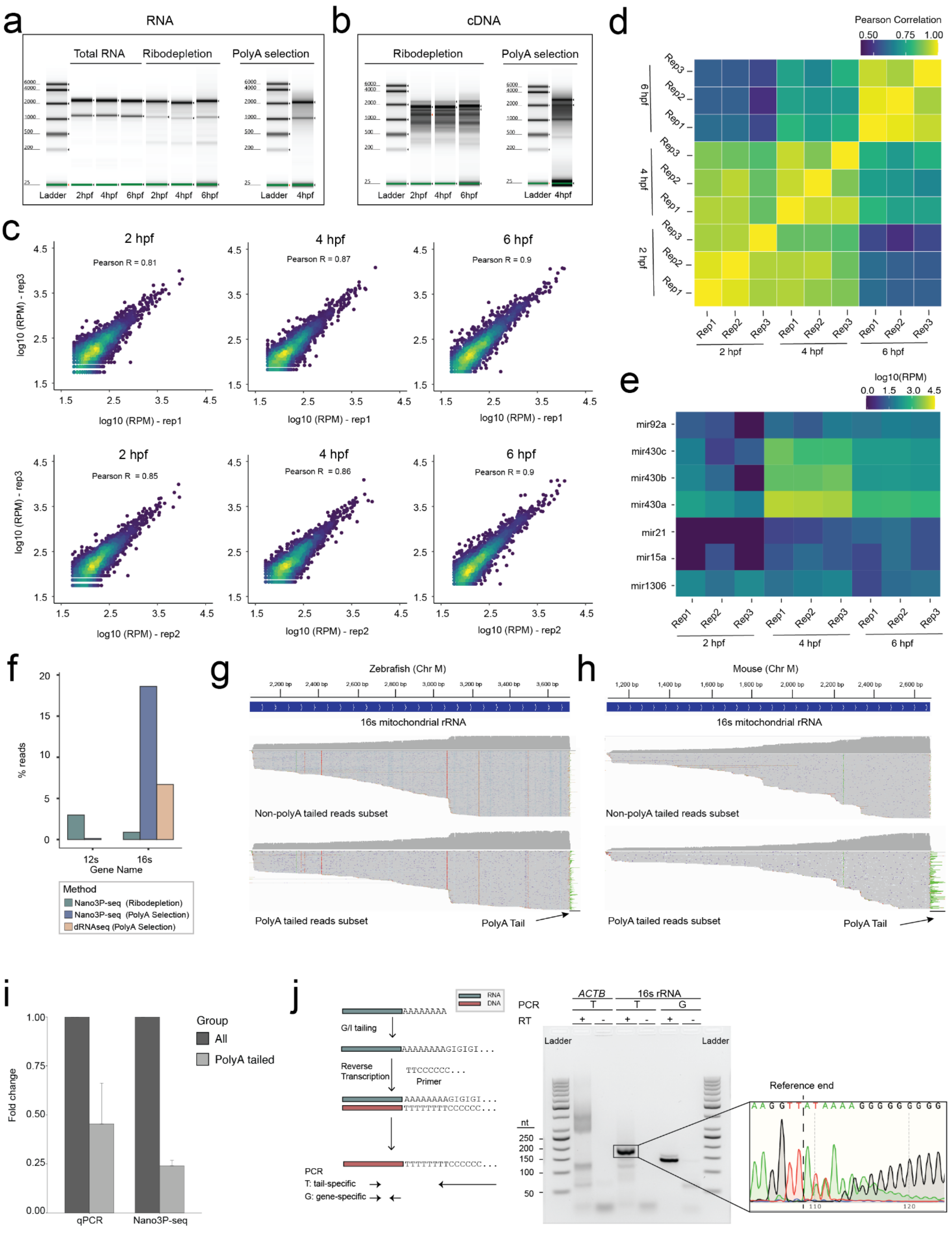
a
RNA
Total RNA
Ribodepletion
PolyA selection
Ladder 2hpf 4hpf 6hpf 2hpf 4hpf 6hpf Ladder 4hpf
b
cDNA
Ribodepletion
PolyA selection
Ladder 2hpf 4hpf 6hpf Ladder 4hpf
c
2 hpf
Pearson R = 0.81
4 hpf
Pearson R = 0.87
6 hpf
Pearson R = 0.9
log10 (RPM) - rep3
log10 (RPM) - rep1
Pearson R = 0.85
Pearson R = 0.86
Pearson R = 0.9
log10 (RPM) - rep2
d
Pearson Correlation
0.50 0.75 1.00
Rep1 Rep2 Rep3
2 hpf
4 hpf
6 hpf
e
log10(RPM)
0.0 1.5 3.0 4.5
mir92a
mir430c
mir430b
mir430a
mir21
mir15a
mir1306
f
% reads
12s
16s
Gene Name
Method
Nano3P-seq (Ribodepletion)
Nano3P-seq (PolyA Selection)
dRNAseq (PolyA Selection)
g
Zebrafish (Chr M)
16s mitochondrial rRNA
Non-polyA tailed reads subset
PolyA tailed reads subset
PolyA Tail
h
Mouse (Chr M)
16s mitochondrial rRNA
Non-polyA tailed reads subset
PolyA tailed reads subset
PolyA Tail
i
Fold change
Group
All
PolyA tailed
qPCR
Nano3P-seq
j
RNA
DNA
AAAAAAAA
G/I tailing
AAAAAAAAGIGIGI...
Reverse Transcription
TTCCCCCC...
Primer
AAAAAAAAGIGIGI...
TTTTTTTTCCCCCC...
TTTTTTTTCCCCCC...
PCR
T: tail-specific
G: gene-specific
PCR
RT
ACTB
16s rRNA
Ladder
nt
Reference end

**Figure S3. Analysis of polyA tail lengths using Nano3P-seq. (a)** Current intensity (pA) plot of a synthetic polyA+ read, obtained using Nano3P-seq. The homopolymeric poly(T) region is highlighted in orange. **(b)** Replicability of median per-gene polyA tail length estimation in sequins captured with Nano3P-seq. The polyA tail length of synthetic sequins is 30 nt (R1 sequins) or 60 nt (R2 sequins). **(c)** Overall comparison of polyA tail length estimation of R1 and R2 sequins which contain 30 nt and 60 nt polyA tail lengths, respectively, obtained using dRNA-seq (orange) and Nano3P-seq (green). Genes with coverage greater than 30 reads are included. **(d)** Per-gene variance of polyA tail length estimations of sequins obtained using dRNA-seq (orange) and Nano3P-seq (green). The number of reads included in the analysis is shown below each boxplot. **(e)** Scatter plot showing the comparison between per-gene variance of polyA tail length estimations obtained using dRNA-seq and Nano3P-seq. **(f)** Correlation between expected tail length (nt) and estimated tail length (nt) of cDNA standards. **(g)** Distribution of polyA tail lengths in mRNAs across zebrafish developmental stages (2, 4 and 6 hpf, shown in blue, green and red respectively) in three biological replicates (shown as full lines, dashed lines, and dotted/dashed lines respectively). **(h)** IGV snapshot of reads mapping to *hist1h2a6* mRNA, which lacks polyA tails. **(i)** Scatterplots of median per-gene polyA tail length estimations using Nano3P-seq and PAL-seq from zebrafish mRNAs at 2 hpf (left), 4 hpf (middle) and 6 hpf (right). Each dot represents the median polyA tail length of a given gene. **(j)** Violin plots depicting the distribution of median per-gene polyA tail length estimations during the zebrafish MZT, estimated using PAL-Seq (left) or Nano3P-seq (right). Statistical comparison of means was made using Kruskal Wallis test. **(k)** Comparative analysis of the abundance (shown as $\log_2$ RPKM) of zebrafish mRNAs that have been binned according to their previously annotated decay mode (maternal decay, zygotic activation-dependent decay, miR-430-dependent decay and no decay) during embryogenesis (t= 2, 4 and 6 hpf) using PAL-seq data. The number of genes included in the analysis is shown below each violinplot. Statistical comparison of means was performed using Kruskal-Wallis test. **(l)** Median per-gene polyA tail length estimations of the 4 groups of zebrafish mRNAs (maternal, miR-430, zygotic and no decay) at 2, 4 and 6 hpf using PAL-seq data. The number of genes included in the analysis is shown below each violinplot. Boxplot limits are defined by lower (bottom) and upper (top) quartile, while the bar indicates the median and whiskers indicate +/- 1.5X inter-quartile range. Statistical analyses were performed using Kruskal-Wallis test. ($p>0.05$:ns, $p\leq0.05$:*, $p\leq0.01$:**, $p\leq0.001$:***, $p\leq0.0001$:****).

**Figure S3**

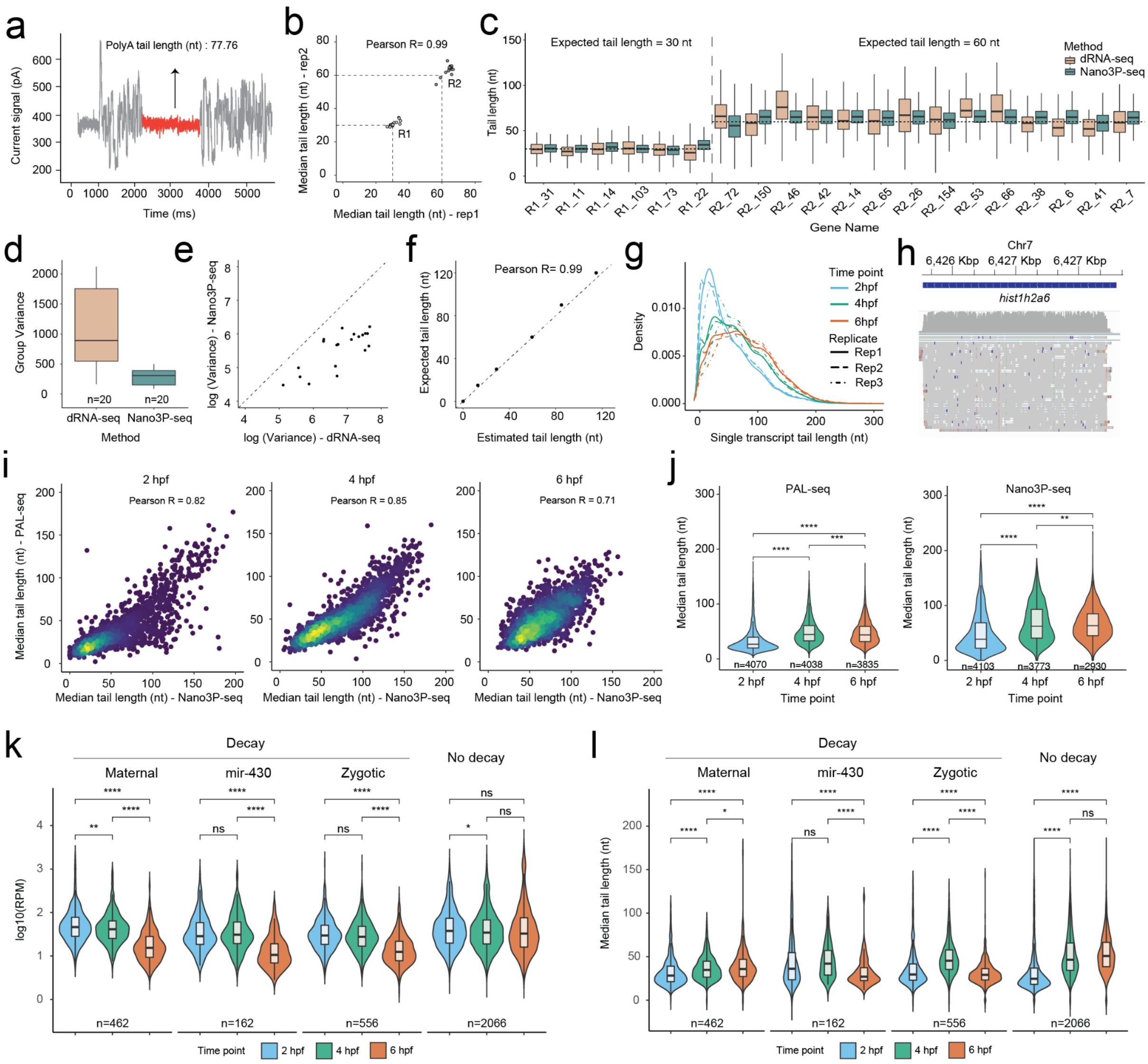
a
PolyA tail length (nt) : 77.76
Current signal (pA)
Time (ms)
b
Pearson R= 0.99
Median tail length (nt) - rep2
Median tail length (nt) - rep1
R1
R2
c
Expected tail length = 30 nt
Expected tail length = 60 nt
Method
dRNA-seq
Nano3P-seq
Tail length (nt)
Gene Name
d
Group Variance
n=20
n=20
dRNA-seq
Nano3P-seq
Method
e
log (Variance) - Nano3P-seq
log (Variance) - dRNA-seq
f
Pearson R= 0.99
Expected tail length (nt)
Estimated tail length (nt)
g
Density
Time point
2hpf
4hpf
6hpf
Replicate
Rep1
Rep2
Rep3
Single transcript tail length (nt)
h
Chr7
6,426 Kbp
6,427 Kbp
6,427 Kbp
hist1h2a6
i
2 hpf
Pearson R = 0.82
4 hpf
Pearson R = 0.85
6 hpf
Pearson R = 0.71
Median tail length (nt) - PAL-seq
Median tail length (nt) - Nano3P-seq
j
PAL-seq
Nano3P-seq
Median tail length (nt)
n=4070
n=4038
n=3835
n=4103
n=3773
n=2930
2 hpf
4 hpf
6 hpf
Time point
k
Decay
Maternal
mir-430
Zygotic
No decay
log10(RPM)
n=462
n=162
n=556
n=2066
Time point
2 hpf
4 hpf
6 hpf
l
Decay
Maternal
mir-430
Zygotic
No decay
Median tail length (nt)
n=462
n=162
n=556
n=2066
Time point
2 hpf
4 hpf
6 hpf

**Figure S4. Analysis of isoform-specific polyA tail and modification dynamics using Nano3P-seq. (a)** IGV coverage tracks of reads mapping to *khdrbs1a* gene from zebrafish embryos at 2 hpf obtained by Nano3P-seq. Annotations of the gene and two main isoforms are shown at the top of the panels. Individual reads mapping to each isoform is illustrated below each coverage track. PolyA tails of individual reads are shown in red. **(b)** Comparison of polyA tail length distributions of reads mapping to *syncrip*, illustrated at the per-gene (left panel) and per-isoform (middle and right panel) level measured at two time points during the zebrafish MZT. Annotations of the gene and two main isoforms are shown at the top of the panels. **(c)** Comparison of polyA tail length distributions of reads mapping to three distinct isoforms (full, dashed and dotted outline) of *ddx5* measured at the three time points during the zebrafish MZT. Annotations of the gene and three main isoforms are shown at the top of the panels. Only isoforms with more than 10 reads are shown. The number of reads included in the analysis is shown below each violinplot. P-values have been computed using the Kruskal–Wallis test and corrected for multiple testing using the Benjamini–Hochberg procedure ($p > 0.05$:ns, $p \leq 0.05$:*, $p \leq 0.01$:**, $p \leq 0.001$:***). **(d)** Comparison of the per-site mismatch frequencies observed in reads mapping to yeast precursor SSU rRNA and to yeast processed SSU rRNA sequenced by dRNA-seq, showing that the unique identified outlier is $m^1acp^3\Psi$. **(e)** IGV coverage tracks of reads mapping to yeast processed small subunit (SSU) rRNA (upper track) and precursor SSU rRNA (lower track), including a magnified image at the position known to be modified with $m^1acp^3\Psi$ (left panel). Positions with a mismatch frequency lower than 0.1 are shown in gray. Mismatch frequency values in yeast precursor and processed SSU rRNA at the position known to be modified with $m^1acp^3\Psi$ (middle panel) (*n* = 3 biological replicates, error bars indicate s.d.). Comparison of the per-site mismatch frequencies observed in reads mapping to yeast precursor SSU rRNA and to yeast processed SSU rRNA, showing that the unique identified outlier is $m^1acp^3\Psi$ (right panel). Boxplot limits are defined by the lower (bottom) and upper (top) quartile. The bar indicates the median, and whiskers indicate +/- 1.5X interquartile range.

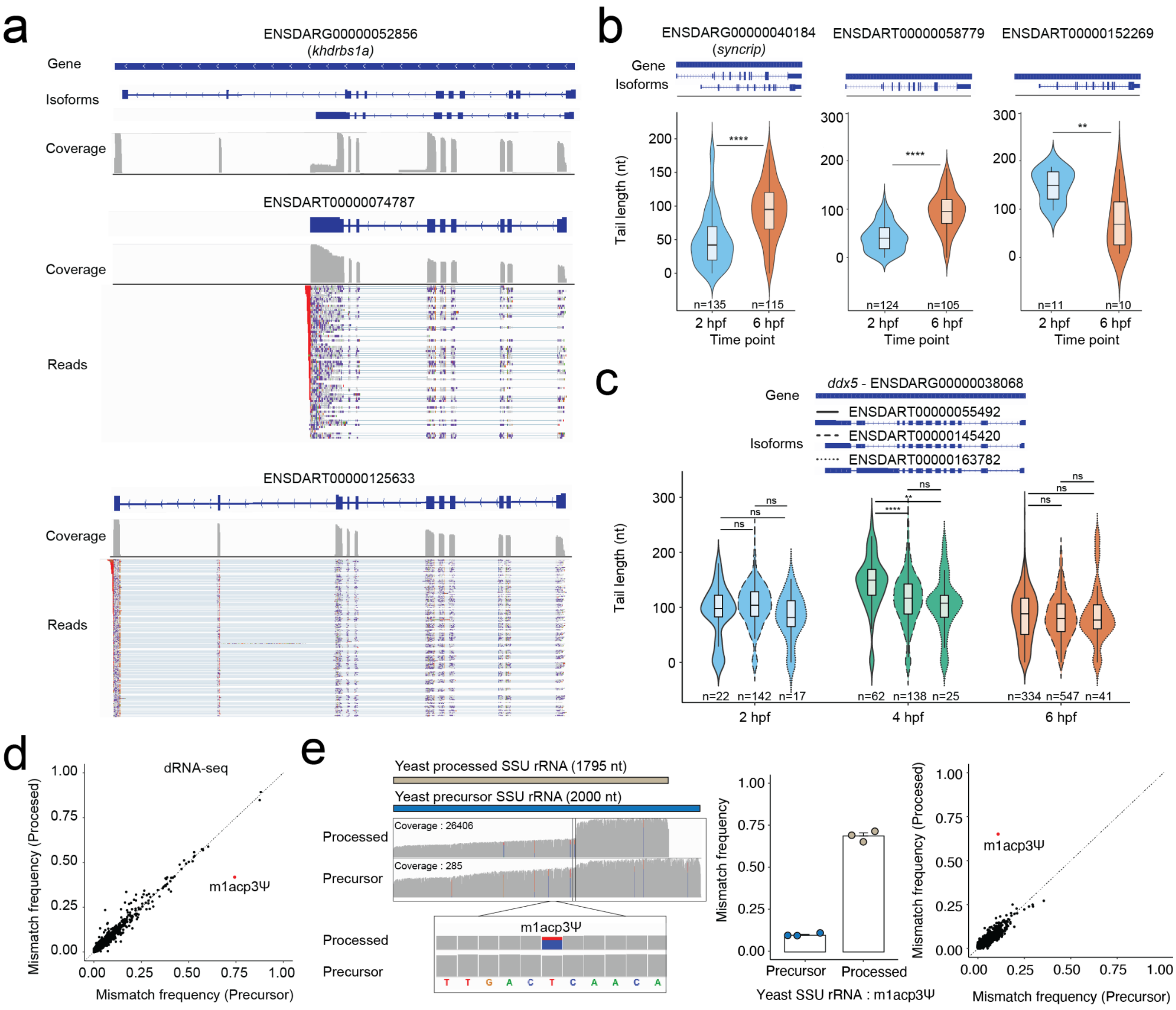

**Figure S5. Analysis of tail composition using Nano3P-seq in *in vitro* samples. (a)** IGV snapshots of nucleotide composition in cDNA standard tails sequenced using Nano3P-seq. Grey regions indicate the mapped part of the reads, whereas colored letters indicate soft-clipped bases (unmapped) which are the base-called tails, after trimming the adapter. **(b)** PolyA tail base frequency distribution (A: green, G: orange, C: blue, U: red) of cDNA standards sequenced with Nano3P-seq.

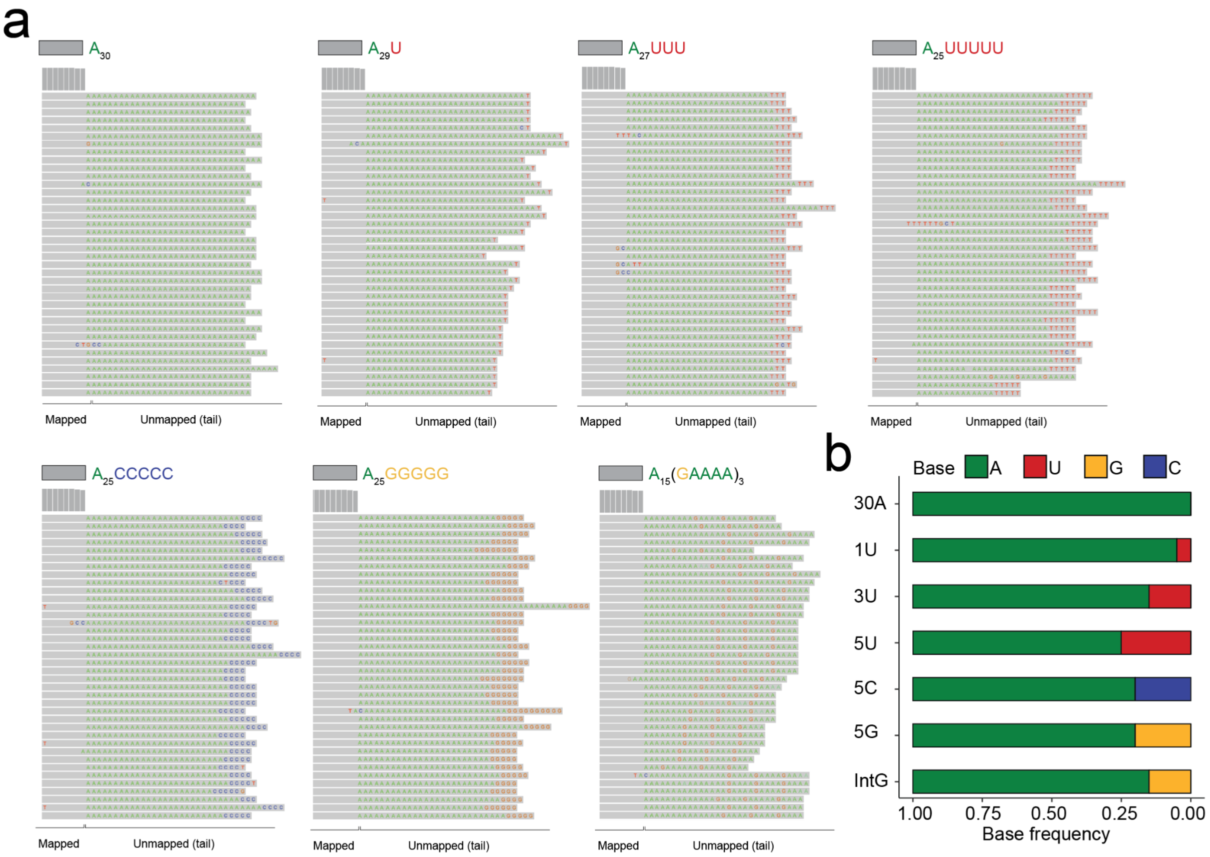

**Figure S5. Analysis of tail composition using Nano3P-seq in *in vivo* samples. (a)** Overall nucleotide composition in mRNA tails at 3 time points during the zebrafish MZT (2, 4 and 6 hpf, shown in blue, green and red respectively) and control that includes sequin R1 and R2 groups of RNAs (gray). P-values have been computed using Kruskal-Wallis test. (*n* = 3 biological replicates, error bars indicate s.d.). **(b)** Probability of base composition (A: green, G: orange, C: blue and U: red) per-position in the last 20 nucleotide of the mRNA tails at 3 time points during the zebrafish MZT (2, 4 and 6 hpf). **(c)** IGV snapshots of reads mapping to zebrafish *ppt2a.4* mRNA (left panel). In the right panel, zoomed images of 3' ends of individual reads with different terminal bases (all reads: top, Term-G reads: bottom) are shown. **(d)** PolyA tail length estimation distributions of mRNA reads belonging to groups classified based on their polyA tail base composition in mouse. **(e)** PolyA tail length estimation distributions of mRNA reads belonging to groups classified based on their polyA tail base composition in yeast. (p>0.05:ns, p≤0.05:*, p≤0.01:**, p≤0.001:***, p≤0.0001:****).

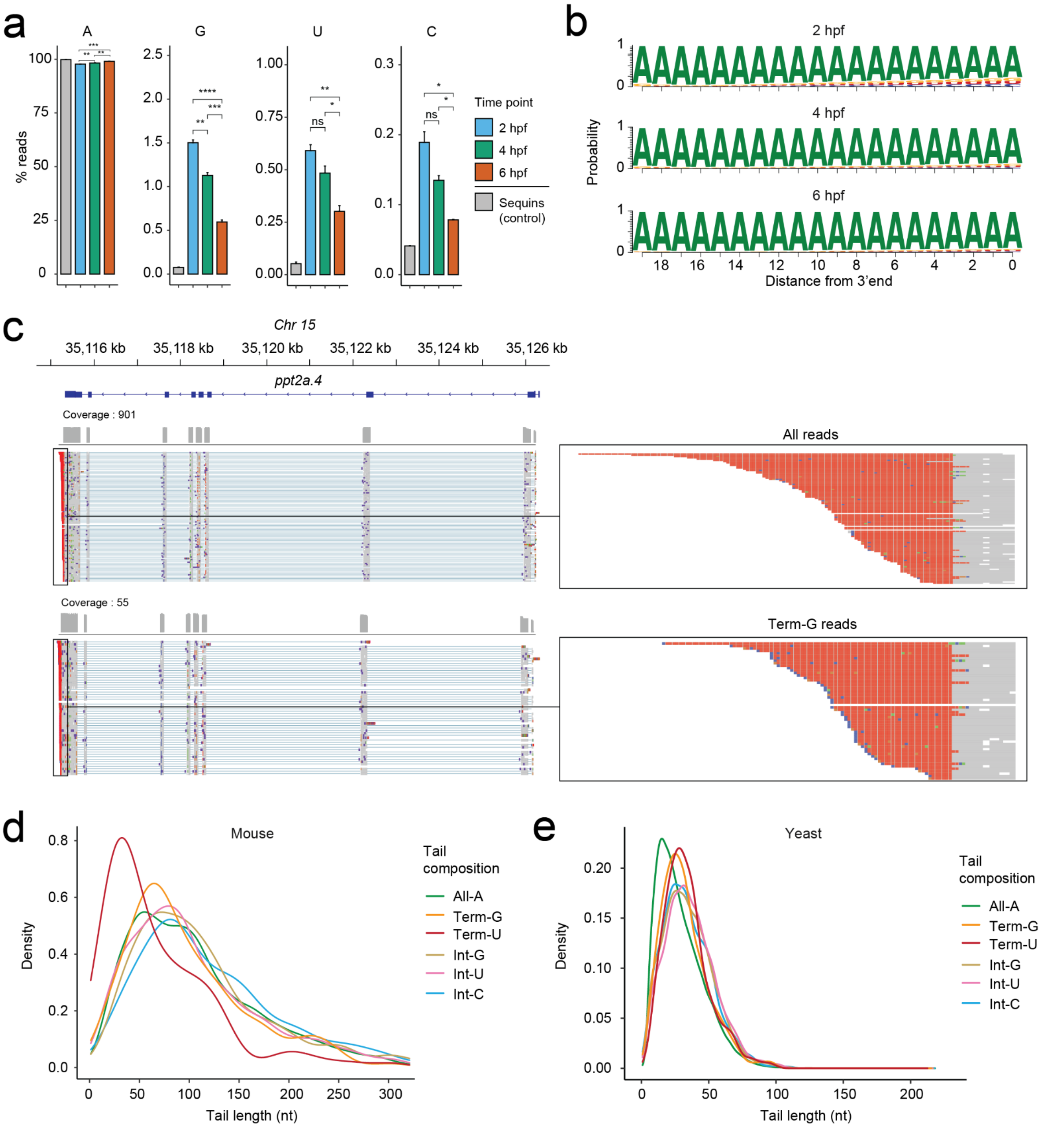

**Figure S7. Comparison of polyA tail length estimations and run stats between dRNA-seq and Nano3P-seq.** **(a)** Distribution of median per-gene polyA tail length estimations from 4 hpf zebrafish embryos, isolated using either polyA selection (green) or ribodepletion (blue). **(b)** Comparison of median per-gene polyA tail length estimations between polyA selected and ribodepleted zebrafish mRNAs isolated at 4 hpf. Each dot represents a gene. **(c)** Comparison of median per-gene polyA tail length estimations of mRNAs in zebrafish ribodepleted samples (replicate 1 and 2) isolated at 4 hpf. Each dot represents a gene. **(d)** Distribution of polyA tail length estimations in mRNAs from 4 hpf zebrafish embryos isolated using polyA selection and sequenced with dRNA-seq (orange) or Nano3P-seq (green). **(e)** Comparison of median per-gene polyA tail length estimations of polyA-selected mRNAs isolated at 4 hpf with dRNA-seq or Nano3P-seq. Each dot represents a gene. **(f)** Read length distribution of mapped reads from 4 hpf zebrafish embryos isolated using polyA selection and sequenced with Nano3P-seq (green) and dRNA-seq (orange), with median lengths of 1307 nt and 907 nt, respectively. **(g)** Sequence identity (%) of the reads from 4 hpf zebrafish embryos isolated using polyA selection and sequenced with either Nano3P-seq (green) or dRNA-seq (orange) with median values of 90.3 and 90.8, respectively. The reads were mapped in both cases to *D. rerio* GRCz11 reference.

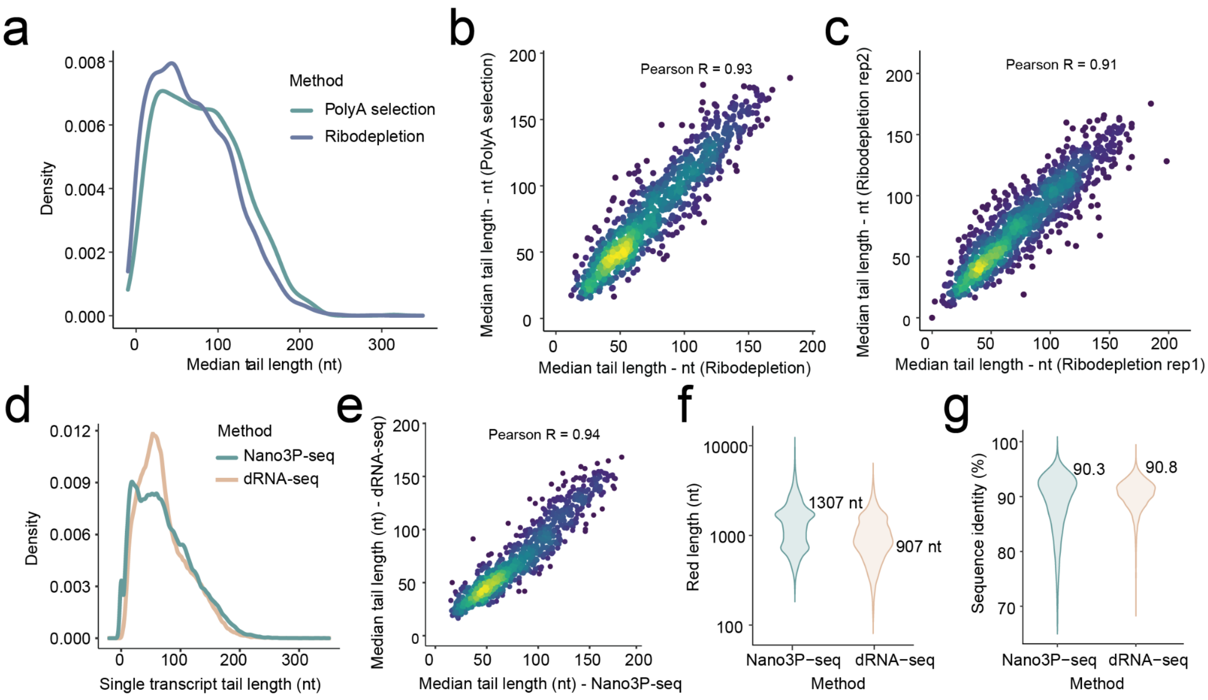

**Figure S8. Comparison of read ends mapping to *actb1* gene before and after filtering by adenine base (A) enrichment. (a)** Reads that are trimmed with porechop. **(b)** Same reads after removing incorrectly trimmed ones (filtered based on their A content). The labeled part indicates the polyA tail, which is dominantly colored in red.

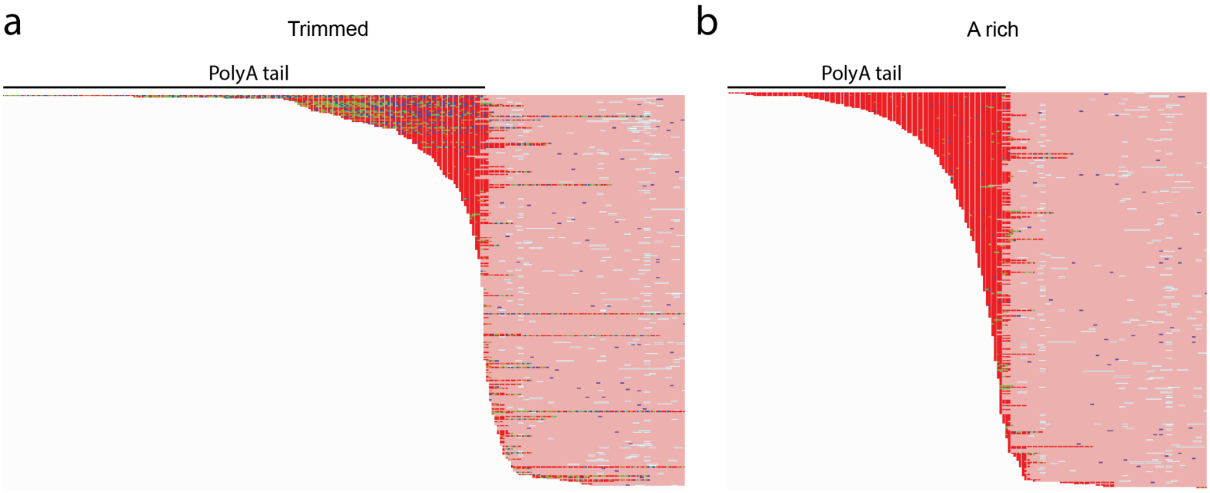
