## Supplementary material for "Nano3P-seq: transcriptome-wide analysis of gene expression and tail dynamics using end-capture nanopore cDNA sequencing": FileS1

Library Preparation protocol for Nano3P-Seq

**List of materials needed**

Materials and consumables required:

- Direct cDNA Sequencing Kit (SQK-DCS109)
- Flow Cell Priming Kit (EXP-FLP001)
- Agencourt AMPure XP beads
- NEB Blunt/TA Ligase Master Mix (M0367)
- 1.5 mL Eppendorf DNA LoBind Tubes
- 0.2 ml thin-walled PCR tubes
- Nuclease-free water
- Freshly prepared 70% ethanol in nuclease-free water
- 10 mM dNTP solution
- TGIRT™-III Enzyme (InGex)
- RNase Inhibitor, Murine (NEB)
- RNase Cocktail Enzyme Mix (ThermoFisher)

Oligos required :

| Oligo Name | Sequence |
| --- | --- |
| D_DNA | /5Phos/CTTCCGATCACTTGCCTGTCGCTCTATCTTCN |
| R_RNA | rGrArArGrArUrArGrArGrCrGrArCrArGrGrCrArArGrUrGrArUrCrGrGrArArG/3SpC3/ |
| CompA_DNA | GAAGATAGAGCGACAGGCAAGTGATCGGAAGA |

**1. Preannealing of the oligos**

We need pre-anneal R_RNA and D_DNA oligos in order to be able to initiate template switching.

| Reagent | Initial Concentration | Volume | Final Conc |
| --- | --- | --- | --- |
| R_RNA | 100 uM | 1 uL | 10 uM |
| D_DNA | 100 uM | 1 uL | 10 uM |
| Tris-Cl pH 7.5 | 0.1 M | 1 uL | 0.01 M |
| NaCl | 0.5 M | 1 uL | 0.05 M |
| RNAse Inhibitor |  | 0.5 uL |  |
| dH2O |  | 5 uL |  |
| Total |  | 10 uL |  |

- Heat the mixture for 94°C for 1 mins and ramp down to RT at 0.1°C/s (in PCR machine).

**2. Reverse Transcription**

| Component | Initial Conc | Volume | Final Conc/Amount |
| --- | --- | --- | --- |
| 5X Reaction Buffer | 2.25 M NaCl, 25 mM MgCl2, 100 mM Tris-HCl, pH 7.5 | 4 uL | 450 mM NaCl, 5 mM MgCl2, 20 mM Tris-HCl, pH 7.5 |
| DTT | 0.1 M | 1 uL | 5 mM |
| Pre-annealed oligos | 10 uM | 2 uL | 1 uM |
| RNA |  | Up to 10 uL | 50-100 ng |
| TGIRT | 10uM- 200 Unit/ul | 1-2 uL | 500 nM- 1000 nM |
| RnaseIN Promega |  | 1 uL |  |
| Total |  | 19 uL |  |

- *Pre-incubate at room temperature for 30 minutes, then add* ***1 ul of 10 mM dNTPs***
- Incubate at 60°C for 1 hour
- Inactivate the enzyme by incubating at 75°C for 15 mins
- Move reaction to ice

**3. RNase treatment**

- Add 1.5 ul RNAse Mix each tube
- 37°C 10 minutes incubation
- Move reaction to ice

**4. Cleanup using Ampure XP Beads**

- Mix the samples with the appropriate volume of beads (17 ul , 0.8 X keeps everything above 150bp, good for getting rid of adapters)
- Mix the beads by flicking
- Incubate 10 minutes at room temperature
- Spin down the tube and place it on the magnet
- Remove the supernatant
- Add 70% freshly prepared 200 ul ethanol to the tube
- Incubate for 30 seconds at room temperature
- Remove the ethanol completely by spinning down and placing back it on magnet
- Air-dry the pellet for maximum 1 minute, do not let it dry out completely!
- Resuspend the beads in 16 ul water
- Incubate 5-10 minutes in RT
- Place the beads on magnet
- Transfer the supernatant into a new tube.
- Quantify 1 μl of eluted sample using a Qubit fluorometer
- Move to the next step

**5. Annealing of Complementary DNA to VNP Oligo**

This step is essential to have a double-stranded DNA oligo with an A overhang, which will initiate the ligation to the adapter

| Components | Initial Concentration | Final Conc. | Volume |
| --- | --- | --- | --- |
| cDNA |  |  | 15 uL |
| Tris-Cl pH 7.5 | 0.1 M | 0.01 M | 2.25 uL |
| NaCl | 0.5 M | 0.05 M | 2.25 uL |
| CompA_DNA | 100 uM | 4.4 uM | 1 uL |
| Water |  |  | 2 uL |
| Total |  |  | 22.5 uL |

- Mix by flicking
- Heat the mixture for 90°C for 1 mins and ramp down to RT at 0.1°C/s (in PCR machine).
- Mix the following

**6. AMX Adapter Ligation**

22.5 uL cDNA-complement mix

2.5 uL Adapter Mix (AMX)

25 uL Blunt/TA Ligase Mixl

- Mix by flicking
- Spin down
- Incubate at RT for 10 minutes

**7. Ampure XP Beads Cleanup**

- Add 25 ul resuspended AMPure XP beads (0.5X) to the reaction and mix by flicking
- Incubate 10 minutes at room temperature
- Thaw ABB Buffer (ABB) and Elution Buffer (EB) at RT, mix by vortexing, spin down and place on ice. Check if the contents of each tube are clear of any precipitate.
- Spin down the tube and place it on the magnet
- Remove the supernatant
- Add 200 ul ABB to the beads. Close the tube lid, and resuspend the beads by pipetting. Return the tube to the magnetic rack, allow beads to pellet and pipette off the supernatant
- Repeat the previous step.
- Remove the ABB completely by spinning down and placing back it on magnet
- Air-dry the pellet for maximum 1 minute, do not let it dry out completely!
- Resuspend the beads in 13 ul Elution Buffer (EB)
- Incubate 5-10 minutes in RT
- Place the beads on magnet
- Place the elute in a 1.5 ml tube
- Quantify 1 μl of eluted sample using a Qubit fluorometer
- The prepared library is used for loading into the MinION flow cell. Store the library on ice until ready to load.
