## Supplementary material for "Nano3P-seq: transcriptome-wide analysis of gene expression and tail dynamics using end-capture nanopore cDNA sequencing": FileS2: annotate-tails.R

```
#' Assign transcript IDs to poly(A) tails
#'
#' This function assigns transcript IDs to the poly(A) tails estimated by
#' tailfindr's find_tails() function. It merges information alignment SAM
#' file and tailfindr's CSV output. The output is written to a file, and also
#' returned as dataframe.
#'
#' @param sam_file The path of alignment SAM file
#' @param tails_csv_file The path of the CSV file produced by find_tails
#' @param output_file The path of the output file
#'
#' @return A dataframe
#' @export
#' @importFrom magrittr "%>%"
annotate_tails <- function(sam_file,
                           tails_csv_file,
                           output_file) {

    #Read SAM file
    sam_list <- read_sam(sam_file)
    df_sam <- sam_list[["x"]]
    QNAME <- RNAME <- MAPQ <- FLAG <- NULL
    read_id <- transcript_id <- mapping_quality <- sam_flag <- NULL
    df_sam <- df_sam %>%
        dplyr::filter(FLAG != 256 & FLAG != 272 & FLAG != 4 & FLAG < 2048) %>%
        dplyr::rename(read_id = QNAME,
                      transcript_id = RNAME,
                      mapping_quality = MAPQ,
                      sam_flag = FLAG) %>%
        dplyr::select(read_id,
                      transcript_id,
                      mapping_quality,
                      sam_flag)


    #Read tails CSV file
    message("Reading poly(A) tail data...")

    df_tails <- read.csv(file = tails_csv_file,
                         header = TRUE,
                         stringsAsFactors = FALSE)

    message("Merging poly(A) tail data and transcript IDs...")

    df <- dplyr::inner_join(df_tails, df_sam, by = 'read_id')

    message("Writing CSV file...")

    data.table::fwrite(df, file = output_file)
    message("Done!")

    return(df)
}
```
