## Supplementary material for "Nano3P-seq: transcriptome-wide analysis of gene expression and tail dynamics using end-capture nanopore cDNA sequencing": FileS2: explore-basecaller-and-fast5type.R

```
#' Explore a fast5 file to find parameters of the experiment
#'
#' This function finds if a read is:
#' \itemize{
#'   \item multifast5 or single fast5
#'   \item 1D read or not-1D
#'   \item basecalled with Albacore or Guppy
#'   \item basecalled using standard model or flipflop model
#'   \item DNA or RNA
#' }
#'
#' @param fast5file_path a character string. Path of a fast5file for determining
#' parameter of the experiment.
#' @param basecall_group a character string. Name of the level
#' in the Fast5 file hierarchy from which to read the data e.g. "Basecall_1D_000"
#'
#' @return A list containing the all the relevant data:
#' \itemize{
#'   \item \code{basecalled_with: 'albacore', 'guppy'}
#'   \item \code{read_is_1d: TRUE, FALSE}
#'   \item \code{model: 'standard', 'flipflop'}
#'   \item \code{fast5type: 'multifast5', 'single'}
#'   \item \code{experiment_type: 'dna', 'rna'}
#' }
#'
#' @examples
#' \dontrun{
#'
#' lst <- explore_basecaller_and_fast5type('/path/to/fast5/file',
#'                                         basecall_group='Basecall_1D_000')
#' }
#'
explore_basecaller_and_fast5type <- function(fast5file_path, basecall_group) {

    f5_obj <- hdf5r::H5File$new(fast5file_path, mode='r')
    f5_obj_names <- f5_obj$names

    # find if the fast5 file is single or multifast5 file
    if (sum(grepl('read_', f5_obj_names)) <= 1) {
        fast5type <- 'single'
        f5_tree <- f5_obj$ls(recursive=TRUE)
    } else {
        fast5type <- 'multi'
        first_read <- f5_obj$open_by_idx(1)
        first_read_name <- f5_obj$names[1]
        f5_tree <- first_read$ls(recursive=TRUE)
    }
    f5_tree <- f5_tree$name

    # find if the reads are 1D
    path_to_check <- paste0('Analyses/', 'Basecall_1D_')
    basecall_1d_found <- sum(grepl(path_to_check, f5_tree))
    read_is_1d <- ifelse(basecall_1d_found > 0, TRUE, FALSE)

    # find if the basecaller is legacy Albacore or the newer Guppy
    if (fast5type == 'multi' & read_is_1d) {
        basecaller_path <- paste0('/', first_read_name, '/Analyses/', basecall_group)
    } else if (fast5type == 'single' & read_is_1d) {
        # handle both guppy and albacore single Fast5 files using regex
        basecaller_path <- paste0('.*Analyses/', basecall_group, '$')
        basecaller_path <- grep(basecaller_path,f5_tree, perl=TRUE, value=TRUE)
    }

    if (read_is_1d) {
        basecaller <- f5_obj[[basecaller_path]]$attr_open('name')$read()
        basecalled_with_albacore <- grepl('Albacore', basecaller)
        basecalled_with_guppy <- grepl('Guppy', basecaller)
        basecalled_with_minknow <- grepl('MinKNOW', basecaller)
        if (basecalled_with_albacore &
            !basecalled_with_guppy &
            !basecalled_with_minknow) {
            basecaller <- 'albacore'
        } else if (!basecalled_with_albacore &
                   basecalled_with_guppy &
                   !basecalled_with_minknow) {
            basecaller <- 'guppy'
        } else if (!basecalled_with_albacore &
                   !basecalled_with_guppy &
                   basecalled_with_minknow) {
            basecaller <- 'minknow'
        }
    } else {
        basecaller <- 'unknown'
    }

    # find if the model used was standard model or flipflop model
    if (basecaller == 'guppy' & read_is_1d) {
        trace <- sum(which(grepl('.*Trace$', f5_tree)))
        model <- ifelse(trace > 0, 'flipflop', 'standard')
    } else if (basecaller == 'albacore') {
        model <- 'standard'
    } else if (basecaller == 'minknow') {
        model <- 'unknown'
    }

    # find if data is dna or rna
    if (fast5type == 'single') {
        context_tags_path <- f5_tree[grepl('.*context_tags$', f5_tree)]
    } else {
        context_tags_path <- paste(first_read_name,
                            '/context_tags',
                            sep = '')
    }
    sequencing_kit <- f5_obj[[context_tags_path]]$attr_open('sequencing_kit')$read()
    if (grepl('rna', sequencing_kit)) {
        experiment_type <- 'rna'
    } else {
        experiment_type <- 'dna'
    }
    f5_obj$close_all()

    # basecalled with : 'albacore' or 'guppy
    # model : 'standard' or 'flipflop'
    # fast5type: 'multi' or 'single
    # read_is_1d: TRUE or FALSE
    # experiment_type: 'dna' or 'rna'
    list(basecalled_with = basecaller,
         read_is_1d = read_is_1d,
         model = model,
         fast5type = fast5type,
         experiment_type = experiment_type)
}
```
