## Supplementary material for "Nano3P-seq: transcriptome-wide analysis of gene expression and tail dynamics using end-capture nanopore cDNA sequencing": FileS2: extract-read-data.R

```
#' Extract the read data from a fast5 file
#'
#' This function can deal with both multifast5 files and single fast5 files.
#' It can handle files basecalled with standard or the flip flop model.
#'
#' @param file_path a character string [NA]. Path of the single fast5 file.
#' Use it if the file to be read is a single fast file. If the file to be
#' read is multifast5 file, then keep this parameter as NA. Also set
#' multifast5 flag to FALSE
#'
#' @param read_id_fast5_file a list [NA]. A list of 'read_id' and 'fast5_file'
#' path. Use this option when a read from a multifast5 file is to be read. In
#' such a case, you should set file_path to NA, and set multifast5 flag to TRUE.
#'
#' @param plot_debug a logical [FALSE]. Should data for plotting debug info in
#' the plots be computed
#'
#' @param basecalled_with a character string. Specify if the data is from
#''albacore' or 'guppy'
#'
#' @param basecall_group a character string. Name of the level
#' in the Fast5 file hierarchy from which to read the data e.g. "Basecall_1D_000"
#'
#' @param multifast5 a logical. Set it to TRUE if the file to be processed
#' is multifast5. Set it to FALSE if the file to be processed is a single fast5
#' file
#'
#' @param model a string. Set to 'flipflop' if the basecalling model is flipflop.
#' Set to 'standard' if the basecalling model is standard model.
#'
#' @param plotting_library a string
#'
#' @param experiment_type a string. Set to 'rna' for RNA data and 'dna' for DNA
#' data
#'
#' @return a list
#'
#' @examples
#' \dontrun{
#'
#' extract_read_data(file_path = '/path/to/the/file',
#'                   read_id_fast5_file = NA,
#'                   plot_debug = F,
#'                   basecalled_with = 'albacore',
#'                   basecall_group = 'Basecall_1D_000',
#'                   multifast5 = TRUE,
#'                   model = 'standard',
#'                   plotting_library = 'rbokeh',
#'                   experiment_type = 'rna')
#' }
#'
extract_read_data <- function(file_path = NA,
                              read_id_fast5_file = NA,
                              plot_debug = FALSE,
                              basecalled_with,
                              basecall_group,
                              multifast5,
                              model,
                              plotting_library,
                              experiment_type) {

    if (!multifast5) {
        f5_obj <- hdf5r::H5File$new(file_path, mode='r')
        f5_tree <- f5_obj$ls(recursive=TRUE)
        f5_tree <- f5_tree$name
        # define all the paths
        raw_signal_path <- grep('.*Signal$', f5_tree, perl = TRUE, value = TRUE)
        if (sum(grepl('Raw/Reads/Read_[0-9]+$', f5_tree)) == 0) {
            read_id_path <-  grep('.*Raw$', f5_tree, perl = TRUE, value = TRUE)
        } else {
            read_id_path <- f5_tree[grepl('Raw/Reads/Read_[0-9]+$', f5_tree)]
        }
        # make fastq, Event/Move path
        event_data_fastq_path <- grep(paste0('.*', basecall_group, '/BaseCalled_template$'),
                                      f5_tree, perl = TRUE, value = TRUE)
        # make segmentation path based on the basecall group
        sp <- strsplit(basecall_group, split = '_')
        seg_group <- sp[[1]][3]
        segmentation_path <- grep(paste0('.*', seg_group, '/Summary/segmentation$'),
                                  f5_tree, perl = TRUE, value = TRUE)
        # make basecalled_template path
        basecall_1d_template_path <- grep(paste0('.*', basecall_group, '/Summary/basecall_1d_template$'),
                                          f5_tree, perl = TRUE, value = TRUE)
    }
    else {
        f5_obj <- hdf5r::H5File$new(read_id_fast5_file$fast5_file, mode='r')
        full_read_id <- read_id_fast5_file$read_id
        # define all the paths
        raw_signal_path <- paste('/', full_read_id, '/Raw/Signal', sep='')
        event_data_fastq_path <- paste0('/', full_read_id, '/Analyses/', basecall_group, '/BaseCalled_template/')
        read_id_path <- paste('/', full_read_id, '/Raw', sep='')

        # make segmentation path based on the basecall group
        sp <- strsplit(basecall_group, split='_')
        seg_group <- sp[[1]][3]
        segmentation_path <- paste0('/', full_read_id, '/Analyses/Segmentation_', seg_group, '/Summary/segmentation')
        basecall_1d_template_path <- paste0('/', full_read_id, '/Analyses/', basecall_group, '/Summary/basecall_1d_template')
    }

    # get the data
    raw_data <- f5_obj[[raw_signal_path]]$read()
    read_id <- f5_obj[[read_id_path]]$attr_open('read_id')$read()
    fastq <- f5_obj[[event_data_fastq_path]]$open('Fastq')$read()

    # ONT's newest data format has no first sample template information
    # as it probably stores the raw data starting from the sample 1
    #start <- f5_obj[[segmentation_path]]$attr_open('first_sample_template')$read()
    start <- tryCatch(
        {
            f5_obj[[segmentation_path]]$attr_open('first_sample_template')$read()
        },
        error = function(e){
            1
        }
    )

    called_events <- f5_obj[[basecall_1d_template_path]]$attr_open('called_events')$read()
    # compute called_event if it has been set to W by ONT in the FAST5 file (Wierd! I know)
    compute_called_events <- ifelse(called_events == 'W', TRUE, FALSE)
    # get the event data, if present -- or make it, if not present
    make_event_data = FALSE
    if ('Events' %in% names(f5_obj[[event_data_fastq_path]])){
        event_data <- f5_obj[[event_data_fastq_path]]$open('Events')$read()
        if (!('length' %in% colnames(event_data))) {
            stride <- f5_obj[[basecall_1d_template_path]]$attr_open('block_stride')$read()
        } else {
            stride <- event_data$length[1]
        }

        # Albacore latest version does not output start column;
        # this is a dirty fix for it
        if (!("start" %in% colnames(event_data))) {
            make_event_data = TRUE
            move <- event_data$move
        }

    } else {
        move <- f5_obj[[event_data_fastq_path]]$open('Move')$read()
        stride <- f5_obj[[basecall_1d_template_path]]$attr_open('block_stride')$read()
        make_event_data = TRUE
    }

    # extract just the fastq removing quality scores
    fastq <- strsplit(fastq, split = "\n")
    fastq <- fastq[[1]][2]


    # if event_data wasn't present, make it now
    if (make_event_data) {
        # The line below is there to remove R CMD CHECK
        # "no visible binding for global variable" error
        move_cumsum <- fastq_bases <- NULL

        start_col <- seq(from = start,
                         to = start + stride * (length(move) - 1),
                         by = stride)
        event_data <- data.frame(move = move,
                                 start = start_col,
                                 move_cumsum = cumsum(move),
                                 fastq_bases = fastq,
                                 stringsAsFactors = FALSE)

        # The line below is there to remove R CMD CHECK
        # "no visible binding for global variable" error
        model_state <- NULL
        event_data <- dplyr::mutate(event_data,
                                    model_state = substr(fastq_bases,
                                                         start=move_cumsum,
                                                         stop=move_cumsum))
        event_data <- dplyr::select(event_data, model_state, start, move)
    }

    # compute the number of event if not present already
    if (compute_called_events) {
        called_events <- nrow(event_data)
    }

    # make a vector of moves interpolated for every sample i.e., make a sample-wise or per-sample vector of moves
    if (plot_debug & plotting_library == 'ggplot2') {
        if (start != 0) {
            moves_sample_wise_vector <- c(rep(NA, start-1),
                                          rep(event_data$move*0.25+1.5, each=stride),
                                          rep(NA, length(raw_data) - start - stride*called_events + 1))
        } else {
            moves_sample_wise_vector <- c(rep(event_data$move*0.25+1.5, each=stride),
                                          rep(NA, length(raw_data) - start - stride*called_events))
        }
    } else {
        moves_sample_wise_vector <- rep(NA, length(raw_data))
    }

    # compute event length vector
    if (model == 'flipflop') {
        # create event length data for tail normalization
        event_length_vector <- rep(NA, called_events)
        count <- 0
        for (i in seq(from=called_events, to=1, by=-1)) {
            if (event_data$move[i] == 1) {
                event_length_vector[i] <- count + 1
                count <- 0
            } else {
                count <- count + 1
            }
        }
        # multiply moves by length of the event (15 for RNA, 5 for DNA)
        event_length_vector <- event_length_vector * stride
        event_data <- cbind(event_data, event_length_vector)
        # remove NAs
        event_length_vector <- event_length_vector[!is.na(event_length_vector)]
        # Normalizer for flip-flop based data
        if (experiment_type == 'rna') {
            # overestimates tail length
            samples_per_nt_1 <- psych::geometric.mean(event_length_vector)
            # Underestimates tail length
            samples_per_nt_2 <- mean(event_length_vector[event_length_vector <= stats::quantile(event_length_vector, 0.90)])
            # Just about right
            samples_per_nt <- (samples_per_nt_1 + samples_per_nt_2)/2
        } else if (experiment_type == 'dna') {
            samples_per_nt <- mean(event_length_vector[event_length_vector <= stats::quantile(event_length_vector, 0.95)])
        }
        # add the start column to the event table for legacy purposes
        start_col <-seq(from=start, to=(start + (nrow(event_data)-1)*stride), by=stride)
        event_data <- dplyr::mutate(event_data, start=start_col)
    } else if (model == 'standard') {
        # create event length data for tail normalization
        l <- stride
        event_length_vector_1 <-  rep(NA, called_events)
        event_length_vector_2 <-  rep(NA, called_events)
        index <- called_events
        length_count <- 1
        divide_by <- 1

        # handle the first row of the event data
        if (event_data$move[1]==1) {
            event_length_vector_1[1] <- 1
        } else {
            event_length_vector_1[index] <- 0.5
            event_length_vector_2[index] <- 0.5
        }
        for(i in (called_events):2) {
            if (event_data$move[i]==2 & event_data$move[i-1]==0) {
                # record the index
                index <- i
                length_count <- 1
                divide_by <- 2
            } else if (event_data$move[i]==1 & event_data$move[i-1]==0) {
                index <- i
                length_count <- 1
                divide_by <- 1
            } else if (event_data$move[i]==0 & (event_data$move[i-1]==1 | event_data$move[i-1]==2)) {
                length_count <- length_count + 1
                # put the previous record
                if (divide_by == 1) {
                    event_length_vector_1[index] = length_count
                } else {
                    event_length_vector_1[index] = length_count/2
                    event_length_vector_2[index] = length_count/2
                }
            } else if ((event_data$move[i]==2 & event_data$move[i-1]==1) | (event_data$move[i]==2 & event_data$move[i-1]==2)) {
                event_length_vector_1[i] <- 0.5
                event_length_vector_2[i] <- 0.5
            } else if ((event_data$move[i]==1 & event_data$move[i-1]==1) | (event_data$move[i]==1 & event_data$move[i-1]==2))  {
                event_length_vector_1[i] <- 1
            } else if  (event_data$move[i]==0 & event_data$move[i-1]==0) {
                length_count <- length_count + 1
            }
        }
        # multiply moves by length of the event (15 for RNA, 5 for DNA)
        event_length_vector_1 <- event_length_vector_1 * l
        event_length_vector_2 <- event_length_vector_2 * l
        #event_data <- dplyr::select(event_data, start, length, move, model_state)
        event_data <- cbind(event_data, event_length_vector_1, event_length_vector_2)
        # combine the two vectors
        event_length_vector <- c(event_length_vector_1, event_length_vector_2)
        # reomve NAs
        event_length_vector <- event_length_vector[!is.na(event_length_vector)]
        # compute geometric mean of modfied Albacore events table to get the normalizer
        samples_per_nt <- psych::geometric.mean(event_length_vector)
    }
    f5_obj$close_all()


    # strip away read_ from the read_id if the reads have it
    read_id = gsub("^read_", "", read_id)

    list(read_id = read_id,
         raw_data = raw_data,
         event_data = event_data,
         moves_sample_wise_vector = moves_sample_wise_vector,
         fastq = fastq,
         start = start,
         stride = stride,
         samples_per_nt = samples_per_nt)
}
```
