## Supplementary material for "Nano3P-seq: transcriptome-wide analysis of gene expression and tail dynamics using end-capture nanopore cDNA sequencing": FileS2: extract-sequence-between-boundaries.R

```
#' Extract sequence between two samples boundaries
#'
#' The functions takes two sample numbers -- \code{start} and \code{end} -- as
#' inputs and finds their corresponding closest matching events in the event
#' table. It then finds the FASTA sequence between these bounding events.
#'
#' @param event_data a data frame or tibble. The event table of a read. Must
#' contain \code{start} and \code{model_state} columns
#' @param start a numeric. Starting sample number
#' @param end a numeric. Ending sample number
#'
#' @return
#' A character string is returned containing FastQ sequencing
#'
#' @examples
#' \dontrun{
#' event_data <- data.frame(start=seq(0, 150, 5),
#'                          model_state=sample(c("A", "T", "C", "G"), 1))
#' sequence <- extract_sequence_between_boundaries(event_data, 98, 131)
#' }
#'
extract_sequence_between_boundaries <- function(event_data, start, end) {
    # get the row index of the start and end raw samples in the event_data
    row_index_start <- which.min(abs(event_data$start - start))
    row_index_end <- which.min(abs(event_data$start - end))

    i <- row_index_start
    fastq_bases <- ''

    # make a fastq sequence
    while (i <= row_index_end){
        if (i == row_index_start){
            fastq_bases <- event_data$model_state[i]
        } else if (event_data$move[i] == 1) {
            fastq_bases <- paste(fastq_bases, substr(event_data$model_state[i], 5, 5), sep='')
        } else if (event_data$move[i] == 2) {
            fastq_bases <- paste(fastq_bases, substr(event_data$model_state[i], 4, 5), sep='')
        }
        i <- i + 1
    }
    return(fastq_bases)
}
```
