## Supplementary material for "Nano3P-seq: transcriptome-wide analysis of gene expression and tail dynamics using end-capture nanopore cDNA sequencing": FileS2: find-dna-tail-per-read.R

```
#' Find poly(A)/poly(T) tail in a single DNA read
#'
#' This function finds poly(A) or poly(T) tail length in DNA reads.
#'
#' @param file_path a character string. Full path of the file that needs to be
#' processed.
#' @param read_id_fast5_file a list. If the read to be processed is in a multi-
#' fast5 file, then a list containing \code{fast5_file} path and \code{read_id}
#' should be provided, otherwise, it should be set to \code{NA}
#' @param basecall_group a character string. Name of the level
#' in the Fast5 file hierarchy from which to read the data e.g. "Basecall_1D_000"
#' @param dna_datatype a character string. Either set to \code{"cdna"} or
#' \code{"pcr-dna"}
#' @param save_plots a logical. Whether to save plots
#' @param show_plots a logical. Whether to show plots in plots window in R-Studio
#' @param plot_debug a logical. Whether to include debug traces in the plots
#' @param save_dir a logical. Directory where plots should be saved
#' @param plotting_library a character string. Whether to use \code{"rbokeh"} or
#' \code{"ggplot2"} for plotting the plots
#' @param multifast5 a logical. Whether the Fast5 file to be processed is a
#' multifast5 file
#' @param basecalled_with a character. Specify whether the data has been basecalled
#' with the \code{"guppy"} or \code{"albacore"} software.
#' @param model a character. Specify whether the data has been basecalled using
#' the \code{"standard"} or the \code{"flipflop"} model
#' @param ... Any optional parameter. Reserved for future.
#'
#' @return a list
#'
find_dna_tail_per_read <- function(file_path = NA,
                                   read_id_fast5_file = NA,
                                   basecall_group = 'Basecall_1D_000',
                                   dna_datatype = 'cdna',
                                   save_plots = FALSE,
                                   show_plots = FALSE,
                                   plot_debug = FALSE,
                                   save_dir = NA,
                                   plotting_library = 'rbokeh',
                                   multifast5 = FALSE,
                                   basecalled_with = 'albacore',
                                   model = 'standard',
                                   ...) {

    do_plots <- ifelse(save_plots | show_plots, TRUE, FALSE)

    data_list <- find_dna_tailtype(file_path = file_path,
                                   basecall_group = basecall_group,
                                   dna_datatype = dna_datatype,
                                   plot_debug = plot_debug,
                                   basecalled_with = basecalled_with,
                                   multifast5 = multifast5,
                                   model= model,
                                   plotting_library = plotting_library,
                                   read_id_fast5_file = read_id_fast5_file,
                                   ...)

    # first read the data and find the tailtype
    if (multifast5) {
        file_path <- read_id_fast5_file$fast5_file
    }

    read_data <- data_list$read_data
    read_id <- read_data$read_id
    event_data <- read_data$event_data
    read_type <- data_list$read_type
    tail_is_valid <- data_list$tail_is_valid
    polya_end <- data_list$polya_end
    polyt_start <- data_list$polyt_start
    samples_per_nt <- read_data$samples_per_nt
    has_precise_boundary <- data_list$has_precise_boundary
    stride <- read_data$stride

    if (!tail_is_valid) {
        return(list(read_id = read_data$read_id,
                    read_type = 'no_adaptor_found',
                    tail_start = NA,
                    tail_end = NA,
                    samples_per_nt = samples_per_nt,
                    tail_length = NA,
                    file_path = file_path,
                    has_precise_boundary = has_precise_boundary))
    }

    # Empirical parameters
    #POLY_T_CNDA_THRESHOLD <- 0.20
    #POLY_A_CNDA_THRESHOLD <- 0.31
    SPIKE_THRESHOLD <- 2.0
    MOVING_WINDOW_SIZE <- 30
    MAX_GAP_BETWEEN_TAILS <- 120 * samples_per_nt
    #SEC_TAIL_MIN_SIZE <- read_data$samples_per_nt * 15
    SLOPE_THRESHOLD <- 0.20

    # Z-normalize the data
    raw_data <- read_data$raw_data
    norm_data <- z_normalize(read_data$raw_data)

    read_length <- length(norm_data)

    # Reverse polyA reads
    if (read_type=='polyA') {
        norm_data <- rev(norm_data)
        polya_start <- read_length - polya_end
        tail_start <- polya_start
    } else {
        tail_start <- polyt_start
    }


    # Recitfy the data (flip anything that is below zero)
    rectified_data <- rectify(norm_data)

    # Windsorize the data (clip anything above a threshold)
    truncated_data <- truncate_spikes(rectified_data,
                                      spike_threshold=SPIKE_THRESHOLD)

    # Smoothen the data
    smoothed_data_lr <- left_to_right_sliding_window_dna('mean',
                                                        truncated_data,
                                                        MOVING_WINDOW_SIZE, 1)
    smoothed_data_rl <- right_to_left_sliding_window_dna('mean',
                                                        truncated_data,
                                                        MOVING_WINDOW_SIZE, 1)
    smoothed_data <- pmin(smoothed_data_lr, smoothed_data_rl)

    ### 1. find the slope between consecutive means with a small smoothing window
    window_size <- 10
    step_size <-  10

    start <- tail_start
    end <- min(start+3000, length(norm_data))
    data <- truncated_data[start:end]
    total <- length(data)
    spots <- seq(from = 1, to = (total - window_size), by = step_size)
    mean_data <- vector(mode="numeric", length = length(spots))
    slope <- vector(mode="numeric", length = length(spots)-1)

    for (i in seq_len(length(spots)-1)) {
        if (i < length(spots)-2) {
            mean_data[i] <- mean(data[spots[i]:(spots[i+1] - 1)])
            mean_data[i+1] <- mean(data[spots[i+1]:(spots[i+2] - 1)])
            slope[i] = (mean_data[i+1] - mean_data[i]) #/ (spots[i+1] - spots[i])
        } else {
            mean_data[i] <- mean(data[spots[i]:(spots[i+1] - 1)])
            mean_data[i+1] <- mean(data[spots[i+1]:length(data)])
            slope[i] = (mean_data[i+1] - mean_data[i]) #/ (length(data) - spots[i])
        }
    }

    # Refine the tail start if the boundary we found based on the adaptor is not precise
    k <- 1
    precise_tail_start <- NA
    if (!has_precise_boundary) {
        while (k < 60) {
            if ((slope[k]   < SLOPE_THRESHOLD) & (slope[k]   > -SLOPE_THRESHOLD) &
                (slope[k+1] < SLOPE_THRESHOLD) & (slope[k+1] > -SLOPE_THRESHOLD) &
                (slope[k+2] < SLOPE_THRESHOLD) & (slope[k+2] > -SLOPE_THRESHOLD) &
                (slope[k+3] < SLOPE_THRESHOLD) & (slope[k+3] > -SLOPE_THRESHOLD) &
                (mean_data[k] < SLOPE_THRESHOLD+0.15) & (mean_data[k] > 0) &
                (mean_data[k+1] < SLOPE_THRESHOLD+0.15) & (mean_data[k+1] > 0) &
                (mean_data[k+2] < SLOPE_THRESHOLD+0.15) & (mean_data[k+2] > 0) &
                (mean_data[k+3] < SLOPE_THRESHOLD+0.15) & (mean_data[k+3] > 0)
                )
            { # fixes the overestimation problem
                precise_tail_start <- (k-1)*window_size + tail_start
                break
            }
            k <- k + 1
        }
    }

    #if it has precise boundary, but the signal
    # is sloping, then find the point where the signal looks like
    # a homopolymer stretch
    m <- 1
    if (has_precise_boundary) {
        while (m < 20) {
            if ((mean_data[m] > SLOPE_THRESHOLD) & (mean_data[m] > 0)) {
                m <- m + 1
            }
            else {
                break
            }
        }
    }


    # Find the tail end
    if (!has_precise_boundary) {
        i <- k
    } else {
        i <- m + 1
        # this is necesarry otherwise the loop below
        # will never find the tail end
    }

    small_glitch_count <- 0
    quit_searching <- FALSE
    tail_end <- NA

    # Main Algorithm:
    # DO while we haven't reached the end of the search window:
    #   - Extend the tail if slope and smoothened data is within the threshold
    #   - if the above criteria fails then:
    #     - check for gap
    #     - there must be at least 60-sample (or more) long tail adjacent to the gap
    #     - the gap should be smaller than 120 nucleotide
    #

    # define smoothened data threshold first
    if (read_type == 'polyA') {
        sm_data_threshold <- 0.6
    } else if  (read_type == 'polyT') {
        sm_data_threshold <- 0.3
    }

    last_good_end <- 1
    while (i < length(slope)){
        if ((slope[i] < SLOPE_THRESHOLD) & (slope[i] > -SLOPE_THRESHOLD) &
            (smoothed_data[tail_start+i*window_size] < sm_data_threshold)) {
            tail_end <- i
            i <- i + 1
            last_good_end <- i
        } else {
            j <- i
            while (j < length(slope)) {
                if (j > floor(MAX_GAP_BETWEEN_TAILS/window_size)) {
                    quit_searching <- TRUE
                    break
                }
                #if ((slope[j] > SLOPE_THRESHOLD) | (slope[j] < -SLOPE_THRESHOLD)) {
                if ((mean_data[j] > SLOPE_THRESHOLD+0.1) | (mean_data[j] < -SLOPE_THRESHOLD-0.1)) {
                    small_glitch_count <- 0
                    j <- j + 1
                } else {
                    small_glitch_count <- small_glitch_count + 1
                    j <- j + 1
                    if (small_glitch_count > 6){ # previously set to 6
                        i <- j
                        break
                    }
                }
            }
            if (quit_searching) {
                i <- last_good_end
                break
            }
            i <- j
        }
    }


    # # tail ends prematurely especially the poly(A) because of the main tail
    # # finding logic above. This code extends that tail so that it is correct
    # while (i < length(slope)) {
    #     if (smoothed_data[tail_start+i * window_size] < sm_data_threshold) {
    #         tail_end <- i
    #         i <- i + 1
    #     } else {
    #         break
    #     }
    # }


    # # tail ends prematurely especially the poly(A) because of the main tail
    # # finding logic above. This code extends that tail so that it is correct
    # while (i < length(slope)) {
    #     if (mean_data[i] < sm_data_threshold/2) {
    #         i <- i + 1
    #     } else {
    #         break
    #     }
    # }
    # tail_end <- i

    if (has_precise_boundary){
        tail_end <- start + (tail_end)*window_size
    } else {
        #tail_end <- start + (tail_end+4)*window_size
        tail_end <- start + (tail_end)*window_size

    }

    mean_data <- c(rep(NA, times=tail_start),
                   rep(mean_data, each=window_size),
                   rep(NA, times=length(truncated_data)-tail_start-length(mean_data)*window_size))

    slope <- c(rep(NA, times=tail_start),
               rep(slope, each=window_size),
               rep(NA, times=length(truncated_data)-tail_start-length(rep(slope, times=window_size))))


    # reverse the data back to its original orientation
    # in the case of polyA reads
    if (do_plots & read_type=='polyA'){
        norm_data <- rev(norm_data)
        truncated_data <- rev(truncated_data)
        smoothed_data <- rev(smoothed_data)
        mean_data <- rev(mean_data)
        slope <- rev(slope)
    }

    ##############################################################
    # set the tail_start to precise_tail_start if it was computed
    # N.B.: the location of this code should not be disturbed
    if (!is.na(precise_tail_start)) {
        tail_start <- precise_tail_start
    }
    #############################################################

    # correct tail start and ends for polyA reads (that have been reversed)
    if (read_type=='polyA'){
        polya_start <- read_length-tail_end
        polya_end <-  read_length-tail_start
        tail_start <- polya_start
        tail_end <- polya_end
    }

    # calculate the tail length
    tail_length = -999
    tail_length = (tail_end - tail_start)/read_data$samples_per_nt

    # validate the tail to remove false positive tails
    is_a_low_confidence_tail = FALSE
    if (!is.na(tail_end)) {
        tail_prox_transcript_variance = sd(raw_data[(tail_end+100):(tail_end+200)])
        tail_variance = sd(raw_data[tail_start:tail_end])
        if (tail_variance > tail_prox_transcript_variance/2.5) { # it is a fake tail
            tail_start <- NA
            tail_end <- NA
            tail_length <- NA
            is_a_low_confidence_tail <- TRUE
        }

    }

    if (save_plots | show_plots) {
        df = data.frame(x=c(0:(length(read_data$raw_data)-1)),
                        raw_data=raw_data,
                        truncated_data=truncated_data,
                        smoothed_data=smoothed_data,
                        moves=read_data$moves_sample_wise_vector-3.0,
                        mean_data=mean_data,
                        slope=slope)

        filename <- paste(read_id, '__', basename(file_path), sep = '')

        if (read_type == 'polyA') {
            if (!is.na(tail_start) & (!is.na(tail_end))) {
                polya_tail <- NULL  # CRAN R CMD
                df['polya_tail'] <- c(rep(NA, times=tail_start-1),
                                      raw_data[tail_start:tail_end],
                                      rep(NA, times=(read_length-tail_end)))
            }
            plot_title <- paste('Poly(A) tail  |  ',
                                'Tail length [nt]: ', round(tail_length, 2), '  |  ',
                                'Tail start: ', tail_start, '  |  ',
                                'Tail end: ', tail_end, '  |  ',
                                'Tail duration [Sa]: ', tail_end-tail_start, '  |  ',
                                'Samples per nt: ', round(samples_per_nt, 2),
                                sep='')
        } else {
            if (!is.na(tail_start) & (!is.na(tail_end))) {
                polyt_tail <- NULL  # CRAN R CMD
                df['polyt_tail'] <- c(rep(NA, times=tail_start-1),
                                      raw_data[tail_start:tail_end],
                                      rep(NA, times=(read_length-tail_end)))
            }
            plot_title <- paste('Poly(T) tail  |  ',
                                'Tail length [nt]: ', round(tail_length, 2), '  |  ',
                                'Tail start: ', tail_start, '  |  ',
                                'Tail end: ', tail_end, '  |  ',
                                'Tail duration [Sa]: ', tail_end-tail_start, '  |  ',
                                'Samples per nt: ', round(samples_per_nt, 2),
                                sep='')
        }

        if (plotting_library == 'rbokeh') {
            if (read_type=='polyA'){
                # p1 polyA/T
                # p2 debug traces
                # p3 moves
                p1 <- rbokeh::figure(data=df,
                                     width=1000, height=200,
                                     legend_location="top_left")
                y <- NULL  # R CMD CHECK
                p3 <- rbokeh::figure(data=data.frame(x=event_data$start,
                                                     y=event_data$move/2),
                                     width=1000, height=100,
                                     legend_location="top_left", ygrid = F)
                if (plot_debug)
                    p2 <- rbokeh::figure(data=df, width=1000, height=400,
                                         legend_location="top_left")

            } else {
                p1 <- rbokeh::figure(data=df,
                                     width=1000, height=200,
                                     legend_location="top_right")
                p3 <- rbokeh::figure(data=data.frame(x=event_data$start,
                                                     y=event_data$move/2),
                                     width=1000, height=100,
                                     legend_location="top_right", ygrid = F)
                if (plot_debug)
                    p2 <- rbokeh::figure(data=df, width=1000, height=400,
                                         legend_location="top_right")
            }

            # poly(A)/(T) plot
            p1 <- rbokeh::ly_lines(p1, x=x, y=raw_data, width=1.5, color='#b2b2b2', legend = "Raw data")
            if (!is.na(tail_start) & (!is.na(tail_end))) {
                if (read_type=='polyT') {
                    p1 <- rbokeh::ly_lines(p1, x=x, y=polyt_tail, width=1.5, color = '#ea3e13', legend = "Poly(T) tail")
                } else {
                    p1 <- rbokeh::ly_lines(p1, x=x, y=polya_tail, width=1.5, color = '#ea3e13', legend = "Poly(A) tail")
                }
            }

            p1 <- rbokeh::y_axis(p1, label='pA', num_minor_ticks=2)
            p1 <- rbokeh::x_axis(p1, label='Sample index')
            p1 <- rbokeh::tool_pan(p1, dimensions = "width")
            p1 <- rbokeh::tool_wheel_zoom(p1, dimensions = "width")

            # plot containing all the debug traces
            if (plot_debug) {
                p2 <- rbokeh::ly_lines(p2, x=x, y=truncated_data, width=1.0, color='#b2b2b2', legend = "Normalized windsorized signal")
                p2 <- rbokeh::ly_lines(p2, x=x, y=smoothed_data, color='#8c8a8a', width=1.5, legend = "Smoothed signal")
                p2 <- rbokeh::ly_lines(p2, x=x, y=slope, color='#f3c963', width=2, legend = "Slope of the signal in the search window")
                p2 <- rbokeh::ly_lines(p2, x=x, y=mean_data, color='#f58585', width=2, legend = "Mean of the signal in the search window")
                #p2 <- rbokeh::ly_lines(p2, moves, color='#b2b2b2', width=1, legend = "Moves")
                p2 <- rbokeh::ly_abline(p2, h=SLOPE_THRESHOLD, color = '#c581f5', type = 3, width=2, legend = "Slope upper bound")
                p2 <- rbokeh::ly_abline(p2, h=-SLOPE_THRESHOLD, color = '#ff6e24', width=2, type = 3, legend = "Slope lower bound")
                if (!is.na(tail_start) & (!is.na(tail_end))) {
                    p2 <- rbokeh::ly_abline(p2, v=tail_start, color = '#4497b9', width=2, legend = "Tail start")
                    p2 <- rbokeh::ly_abline(p2, v=tail_end, color = '#879833', width=2, legend = "Tail end")
                }
                p2 <- rbokeh::y_axis(p2, label='z-normalized data values', num_minor_ticks=4, desired_num_ticks = 5)
                p2 <- rbokeh::x_axis(p2, label='Sample index')
                p2 <- rbokeh::tool_pan(p2, dimensions = "width")
                p2 <- rbokeh::tool_wheel_zoom(p2, dimensions = "width")

                # does not work
                # if (read_type == 'polyA') {
                #     p2 <- rbokeh::y_axis(p2, position = "right")
                # }
            }

            # plot containing the moves
            p3 <- rbokeh::ly_crect(p3,
                                   x=x,
                                   y=y,
                                   width=rep(0.01, nrow(event_data)),
                                   height=event_data$move,
                                   color='#529c82')
            p3 <- rbokeh::y_axis(p3, label='', num_minor_ticks=0, desired_num_ticks = 3)
            p3 <- rbokeh::x_axis(p3, label='Sample index')
            p3 <- rbokeh::tool_pan(p3, dimensions = "width")
            p3 <- rbokeh::tool_wheel_zoom(p3, dimensions = "width")
            p3 <- rbokeh::tool_wheel_zoom(p3, dimensions = "height")

            if (plot_debug){
                lst <- list(p1, p2, p3)
                names(lst) <- c(plot_title, 'Debugging Traces', 'Moves')
                nrow <- 3
            } else {
                lst <- list(p1, p3)
                names(lst) <- c(plot_title, 'Moves')
                nrow <- 2
            }
            suppressWarnings({
            p <- rbokeh::grid_plot(lst,
                                   nrow = nrow,
                                   link_data = TRUE,
                                   same_axes=c(TRUE, FALSE))
            })

        } else { # plotting_library == 'ggplot2'
            if (plot_debug) {
                p <- ggplot2::ggplot(data=df, ggplot2::aes(x = x)) +
                    ggplot2::geom_line(ggplot2::aes(y = truncated_data), color = '#4040a1')+
                    ggplot2::geom_line(ggplot2::aes(y = moves), color = '#b2b2b2') +
                    ggplot2::geom_hline(yintercept = SLOPE_THRESHOLD, color = '#00B0DF', linetype = 'dotted') +
                    ggplot2::geom_line(ggplot2::aes(y = slope, color = '#BF1268')) +
                    ggplot2::geom_hline(yintercept = -SLOPE_THRESHOLD, color = '#FF94CD', linetype = 'dotted') +
                    ggplot2::geom_line(ggplot2::aes(y = smoothed_data), color='#060B54') +
                    ggplot2::geom_line(ggplot2::aes(y = mean_data, color = '#F79A14')) +
                    ggplot2::geom_line(ggplot2::aes(y = c(rep(NA, times=tail_start-1),
                                                          truncated_data[tail_start:tail_end],
                                                          rep(NA, times=(read_length-tail_end)))), color='#BF1268') +
                    ggplot2::ylab('z-normalized data values')
            } else {
                p <- ggplot2::ggplot(data=df, ggplot2::aes(x = x)) +
                    ggplot2::geom_line(ggplot2::aes(y = raw_data), color = '#8E8E8E') +
                    ggplot2::ylab('pA')
            }
            if (!is.na(tail_end) & !is.na(tail_start) & plot_debug==FALSE) {
                p <- p + ggplot2::geom_line(ggplot2::aes(y = c(rep(NA, times=tail_start-1),
                                                               raw_data[tail_start:tail_end],
                                                               rep(NA, times=(read_length-tail_end)))), color='#BF1268')
            }
            p <- p + ggplot2::ggtitle(plot_title) +
                ggplot2::xlab('Sample index')
        }

        if (show_plots) print(p)
        if (save_plots) {
            filename_png <- paste(filename,'.png', sep='')
            filename_html <- paste(filename,'.html', sep='')
            save_path_png <- file.path(save_dir, 'plots', filename_png, fsep = .Platform$file.sep)
            save_path_html <- file.path(save_dir, 'plots', filename_html, fsep = .Platform$file.sep)
            if (plotting_library == 'ggplot2') {
                ggplot2::ggsave(save_path_png, plot = p, width = 300, height = 70, units = 'mm')
            } else {
                #rbokeh::widget2png(p, file = save_path_png)
                rbokeh::rbokeh2html(p, file = save_path_html)

                # Hack to make the plots working
                tx  <- readLines(save_path_html)
                tx2  <- gsub(pattern="bokeh-0.12.5", replace="bokeh-0.12.3", x=tx)
                writeLines(tx2, con=save_path_html)
            }
        }
    }

    # if tail end is not found for whatever reason
    # perhaps due to wrong alignment location of end primer
    # then return
    if (is.na(tail_end)) {
        if (is_a_low_confidence_tail) {
            return(list(read_id = read_data$read_id,
                        read_type = 'qc_failed',
                        tail_start = NA,
                        tail_end = NA,
                        samples_per_nt = samples_per_nt,
                        tail_length = NA,
                        file_path = file_path,
                        has_precise_boundary = has_precise_boundary))
        } else {
            return(list(read_id = read_data$read_id,
                        read_type = 'contains_no_polyT_tail',
                        tail_start = NA,
                        tail_end = NA,
                        samples_per_nt = samples_per_nt,
                        tail_length = 0,
                        file_path = file_path,
                        has_precise_boundary = has_precise_boundary))
        }
    } else {
        return(list(read_id = read_data$read_id,
                    read_type = 'contains_a_polyT_tail',
                    tail_start = tail_start,
                    tail_end = tail_end,
                    samples_per_nt = samples_per_nt,
                    tail_length = tail_length,
                    file_path = file_path,
                    has_precise_boundary = has_precise_boundary))
    }


}
```
