## Supplementary material for "Nano3P-seq: transcriptome-wide analysis of gene expression and tail dynamics using end-capture nanopore cDNA sequencing": FileS2: find-dna-tailtype.R

```
#' Finds if a DNA read is poly(A) read or poly(T) read
#'
#' This function reads the data from a fast5 file, and then alings primers to
#' the read to discover if it is a poly(A) or poly(T) read. For poly(A) reads,
#' the function further tests if the read is a complete read -- and not truncated
#' prematurely. The function also find the rough end site of the poly(A) tail,
#'  and the rough start site of the poly(T) tail.
#'
#' @param file_path a character string[NA]. Full path of the read whose type is
#' to be determined. Use it if the read is basecalled with Albacore and is of
#' one-read-per-fast5 type.
#' @param dna_datatype a character string ['cdna']. Specify if the read is 'cdna'
#' or pcr-dna'.
#' @param plot_debug a logical [FALSE]. Specifies whether to compute data needed
#'  for plotting debug.
#' @param basecalled_with a character string. Specify if the data is from
#''albacore' or 'guppy'
#' @param multifast5 a logical. Set it to TRUE if the file to be processed
#' is multifast5. Set it to FALSE if the file to be processed is a single fast5
#' file
#' @param model a string. Set to 'flipflop' if the basecalling model is flipflop.
#' Set to 'standard' if the basecalling model is standard model.l
#'
#' @param plotting_library a string.
#'
#' @param read_id_fast5_file a list [NA]. A list of 'read_id' and 'fast5_file'
#' path. Use this option when a read from a multifast5 file is to be read. In
#' such a case, you should set file_path to NA, and set multifast5 flag to TRUE.
#' @param ... An other parameter. For future expansion.
#' @examples
#' \dontrun{
#'
#' # 1. If the data is multifast5 cDNA (direct cDNA or amplified cDNA)
#' data basecalled with flip-flop algorithm
#' read_id_fast5_file = list(read_id=read_id, fast5_file=full_path_of_fast5_file)
#' find_dna_tailtype(dna_datatype = 'cdna',
#'                   multifast5 = TRUE,
#'                   basecalled_with = 'guppy',
#'                   model = 'flipflop',
#'                   read_id_fast5_file = read_id_fast5_file)
#'
#' # 2. If the data is multifast5 pcr-DNA data basecalled with flip-flop
#' algorithm
#' read_id_fast5_file = list(read_id=read_id, fast5_file=full_path_of_fast5_file)
#' find_dna_tailtype(dna_datatype = 'pcr-dna',
#'                   multifast5=TRUE,
#'                   basecalled_with = 'guppy',
#'                   model = 'flipflop',
#'                   read_id_fast5_file = read_id_fast5_file)
#'
#' # 3. If the data is cDNA (direct cDNA or amplified cDNA) data basecalled with
#' albacore with single fast5 files as output
#' find_dna_tailtype(file_path = full_file_path_of_the_read,
#'                   dna_datatype = 'cdna',
#'                   multifast5 = FALSE,
#'                   basecalled_with = 'albacore',
#'                   model = 'standard')
#' }
#'
#' @return A list containing all the relevant information
#'

find_dna_tailtype <- function(file_path = NA,
                              basecall_group = 'Basecall_1D_000',
                              dna_datatype = 'cdna',
                              plot_debug = FALSE,
                              basecalled_with,
                              multifast5,
                              model,
                              read_id_fast5_file = NA,
                              plotting_library,
                              ...) {

    # get the substitution matrix -- if passed from outside
    if (length(list(...)) > 0) {
        lst <- list(...)
        if ("dna_opts" %in% names(lst)) {
            dna_opts <- lst$dna_opts
            match <- dna_opts$match
            mismatch <- dna_opts$mismatch
            type <- dna_opts$type
            gapOpening <- dna_opts$gapOpening
            gapExtension <- dna_opts$gapExtension
            submat <- dna_opts$submat
            if (dna_datatype == 'custom-cdna') {
                fp <- dna_opts$fp
                ep <- dna_opts$ep
            }
        }
    } else {
        # otherwise make one
        match <- 1
        mismatch <- -1
        type <- 'local'
        gapOpening <- 0
        gapExtension <- 1
        submat <- Biostrings::nucleotideSubstitutionMatrix(match = match,
                                                           mismatch = mismatch,
                                                           baseOnly = TRUE)
    }

    # extract read data
    read_data <- extract_read_data(file_path,
                                   read_id_fast5_file,
                                   plot_debug,
                                   basecalled_with,
                                   basecall_group,
                                   multifast5,
                                   model,
                                   plotting_library,
                                   experiment_type = 'dna')

    # get event data table and the fastQ
    #event_data <- read_data$event_data
    fastq <- read_data$fastq


    # define the primer sequences
    if (dna_datatype == 'cdna') {
        # Nano3P seq front and end primers
        ep = Biostrings::DNAString('CTTCTATCTCGCTGTCCGTTCACTAGCCTTC')
        #ep = Biostrings::DNAString('TCTTCTATCTCGCTGTCCGTTCA')
        fp = NA
        threshold <- 0.30 #raised the threshold from 0.35 to 0.40
    } else if (dna_datatype == 'pcr-dna') {
        fp <- Biostrings::DNAString('ATTTAGGTGACACTATAGCGCTCCATGCAAACCTGTC')
        ep <- Biostrings::DNAString('GAGTCCGGGCGGCGC')
        threshold <- 0.68
    } else if (dna_datatype == 'custom-cdna') {
        fp <- Biostrings::DNAString(fp)
        ep <- Biostrings::DNAString(ep)
        threshold <- 0.6
    }

    # RECIPE:
    # 1. Tail is poly(A) if nas_fp > nas_ep > 0.6 --> check if its not prematurely terminated read
    #                                                 by checking for rc_ep at the end of the of the read
    # 2. Tail is poly(T) if nas_ep > nas_fp > 0.6
    # 3. Invalid otherwise

    # define a search window width within which to find the ep and fp
    search_window <- 150

    as_ep <- Biostrings::pairwiseAlignment(pattern=ep,
                                           subject=Biostrings::DNAString(substr(fastq, start=1, stop=min(search_window, nchar(fastq)))),
                                           substitutionMatrix=submat,
                                           type=type,
                                           scoreOnly=FALSE,
                                           gapOpening=gapOpening,
                                           gapExtension=gapExtension)


    nas_ep <- as_ep@score/ep@length
    has_precise_boundary <- FALSE
    bases_to_match <- 3

    if (nas_ep > threshold) {
        read_type <- 'polyT'
        tail_is_valid <- T
        polyt_start_fastq <- as_ep@subject@range@start + as_ep@subject@range@width
        polyt_rough_start <- find_sample_index_for_fastq_base(read_data$event_data, polyt_start_fastq, read_type)
        polya_end_fastq <- NA
        polya_rough_end <- NA

        # check if we have captured the end of the tail perfectly
        # by finding out if we captured the bases adjacent to the tails
        if (substr(as_ep@subject,
                   start=nchar(as.character(as_ep@subject))-bases_to_match+1,
                   stop=nchar(as.character(as_ep@subject))) == substr(ep,
                                                                      start=ep@length-bases_to_match+1,
                                                                      stop=ep@length)) {
            has_precise_boundary <- F#TRUE
        }


        # if the above two checks fail then it is an invalid read
    } else {
        read_type <- 'invalid'
        tail_is_valid <- F
        polyt_start_fastq <- NA
        polyt_rough_start <- NA
        polya_end_fastq <- NA
        polya_rough_end <- NA
        has_precise_boundary <- NA
    }

    # df <- data.frame(read_data$raw_data)
    # p <- ggplot2::ggplot(data=df) +
    #     ggplot2::geom_line(ggplot2::aes(x=c(1:length(read_data$raw_data)), y=read_data.raw_data))
    #
    # if (!is.na(polyt_rough_start)){
    #     p <- p + ggplot2::geom_vline(xintercept = polyt_rough_start, color='red')
    # }
    # if (!is.na(polya_rough_end)){
    #     p <- p + ggplot2::geom_vline(xintercept = polya_rough_end, color='green')
    # }
    #
    # print(p)

    # nchar(fastq)
    list(read_data = read_data, # for max comment it out
         read_type = read_type,
         tail_is_valid = tail_is_valid,
         polya_end_fastq = polya_end_fastq,
         polyt_start_fastq = polyt_start_fastq,
         polya_end = polya_rough_end,
         polyt_start = polyt_rough_start,
         has_precise_boundary = has_precise_boundary
    )
    #nas_fp=nas_fp,
    #nas_rc_ep=nas_rc_ep,
    #nas_ep=nas_ep,
    #nas_rc_fp=nas_rc_fp))
}
```
