## Supplementary material for "Nano3P-seq: transcriptome-wide analysis of gene expression and tail dynamics using end-capture nanopore cDNA sequencing": FileS2: find-rna-polya-crude-start-end.R

```
#' Find the crude start and end of polyA tail in RNA read
#'
#' @param rle_lengths a numeric vector. A vector of RLE lengths in samples
#' @param rle_values a logical vector. A vector of logical RLE values
#' @param rle_indices a numeric vector. A vector if cumsum of RLE lengths
#' @param trough_threshold a numeric. The threshold below which to merge sections
#'
#' @return a list
#'
find_rna_polya_crude_start_end <- function(rle_lengths,
                                           rle_values,
                                           rle_indices,
                                           trough_threshold) {

    polya_crude_start <- NA
    polya_crude_end <- NA

    for (i in seq_len(length(rle_lengths)-1)) {
        # just check for a big enough trough in the begining of the read
        # followed by a peak in rle instersections
        if (rle_lengths[i] > trough_threshold &&
            !rle_values[i] && rle_values[i+1])  {
            polya_crude_start <- rle_indices[i]
            polya_crude_end <- rle_indices[i+1]
            break
        }
    }
    return (list(start=polya_crude_start, end=polya_crude_end))
}
```
