## Supplementary material for "Nano3P-seq: transcriptome-wide analysis of gene expression and tail dynamics using end-capture nanopore cDNA sequencing": FileS2: find-rna-polya-precise-start-end.R

```
#' Finds the precise RNA tail and start
#'
#' @param crude_polya_boundries list. A list of crude \code{start} and \code{end}
#' of the poly(A) tail
#' @param truncated_data numeric vector
#' @param POLY_A_RNA_THRESHOLD numeric
#' @param save_plots logical
#' @param show_plots logical
#'
#' @return a list
#'
find_rna_polya_precise_start_end <- function(truncated_data,
                                             POLY_A_RNA_THRESHOLD,
                                             crude_polya_boundries,
                                             save_plots,
                                             show_plots){

    crude_start <- crude_polya_boundries$start
    crude_end <- crude_polya_boundries$end

    precise_start <- NA
    precise_end <- NA

    if (!is.na(crude_start)) {

        ### 1. find the slope between consecutive means with a small smoothing window
        window_size <- 25
        step_size <-  25

        data <- truncated_data[crude_start:crude_end]
        total <- length(data)

        spots <- seq(from = 1, to = (total - window_size), by = step_size)

        mean_data <- vector(mode="numeric", length = length(spots))
        slope <- vector(mode="numeric", length = length(spots)-1)

        for (i in seq_len(length(spots)-1)) {
            if (i < length(spots)-2) {
                mean_data[i] <- mean(data[spots[i]:(spots[i+1] - 1)])
                mean_data[i+1] <- mean(data[spots[i+1]:(spots[i+2] - 1)])
                slope[i] = (mean_data[i+1] - mean_data[i]) #/ (spots[i+1] - spots[i])
            } else {
                mean_data[i] <- mean(data[spots[i]:(spots[i+1] - 1)])
                mean_data[i+1] <- mean(data[spots[i+1]:length(data)])
                slope[i] = (mean_data[i+1] - mean_data[i]) #/ (length(data) - spots[i])
            }
        }

        ### 2. Find the precise start
        # find the precise start of the poly(A), which is when the slope goes up in the begining and
        # then just starts to go down && the mean signal should be above - POLY_A_RNA_THRESHOLD
        for (i in seq_len(length(slope)-1)) {
            if ((slope[(i+1)] < slope[i]) &&
                #(slope[i] > POLY_A_RNA_THRESHOLD) &&
                (mean_data[i+1] > -POLY_A_RNA_THRESHOLD)) {
                precise_start <- i * window_size + crude_start
                break
            }
        }

        ### 3. Find the precise end
        # now that we have found the precise start, we now find the precise end
        # we move from the end and look at the four previous slopes
        # if the four previous slopes are between 0.3 and -0.3,
        # and the mean data is greater than the poly(A) threshold then we declare the
        # current slope as the potential precise end. We repeat this until we reach the very end.
        for (k in length(slope):(i+4) ) {
            if ((slope[k] < POLY_A_RNA_THRESHOLD && slope[k] > -POLY_A_RNA_THRESHOLD) &&
                (slope[k-1] < POLY_A_RNA_THRESHOLD && slope[k-1] > -POLY_A_RNA_THRESHOLD) &&
                (slope[k-2] < POLY_A_RNA_THRESHOLD && slope[k-2] > -POLY_A_RNA_THRESHOLD) &&
                (slope[k-3] < POLY_A_RNA_THRESHOLD && slope[k-3] > -POLY_A_RNA_THRESHOLD) &&
                (mean_data[k-1] > 0) &&
                (mean_data[k-2] > 0) &&
                (mean_data[k-3] > 0) &&
                (mean_data[k-4] > 0)) {
                precise_end <- (k+1) * window_size + crude_start
                break
            }
        }

        # Do a second round of finding the precise end, but this time make the
        # criteria even more stringent
        tryCatch({
            for (m in k:(i+5) )  {
                if ((slope[m] < POLY_A_RNA_THRESHOLD && slope[m] > -POLY_A_RNA_THRESHOLD) &&
                    (slope[m-1] < POLY_A_RNA_THRESHOLD && slope[m-1] > -POLY_A_RNA_THRESHOLD) &&
                    (slope[m-2] < POLY_A_RNA_THRESHOLD && slope[m-2] > -POLY_A_RNA_THRESHOLD) &&
                    (slope[m-3] < POLY_A_RNA_THRESHOLD && slope[m-3] > -POLY_A_RNA_THRESHOLD) &&
                    (slope[m-4] < POLY_A_RNA_THRESHOLD && slope[m-4] > -POLY_A_RNA_THRESHOLD) &&
                    (slope[m-5] < POLY_A_RNA_THRESHOLD && slope[m-5] > -POLY_A_RNA_THRESHOLD) &&
                    (mean_data[m-1] > 0) &&
                    (mean_data[m-2] > 0) &&
                    (mean_data[m-3] > 0) &&
                    (mean_data[m-4] > 0) &&
                    (mean_data[m-5] > 0) &&
                    (mean_data[m-6] > 0)) {
                    precise_end <- (m+1) * window_size + crude_start
                    break
                }
            }

        },
        error=function(e) e)

        # Do a third round of finding the precise end, but this time make the
        # criteria even more stringent
        tryCatch({
            for (j in m:(i+12) )  {
                if ((slope[j] < POLY_A_RNA_THRESHOLD && slope[j] > -POLY_A_RNA_THRESHOLD) &&
                    (slope[j-1] < POLY_A_RNA_THRESHOLD && slope[j-1] > -POLY_A_RNA_THRESHOLD) &&
                    (slope[j-2] < POLY_A_RNA_THRESHOLD && slope[j-2] > -POLY_A_RNA_THRESHOLD) &&
                    (slope[j-3] < POLY_A_RNA_THRESHOLD && slope[j-3] > -POLY_A_RNA_THRESHOLD) &&
                    (slope[j-4] < POLY_A_RNA_THRESHOLD && slope[j-4] > -POLY_A_RNA_THRESHOLD) &&
                    (slope[j-5] < POLY_A_RNA_THRESHOLD && slope[j-5] > -POLY_A_RNA_THRESHOLD) &&
                    (slope[j-5] < POLY_A_RNA_THRESHOLD && slope[j-6] > -POLY_A_RNA_THRESHOLD) &&
                    (slope[j-5] < POLY_A_RNA_THRESHOLD && slope[j-7] > -POLY_A_RNA_THRESHOLD) &&
                    (slope[j-5] < POLY_A_RNA_THRESHOLD && slope[j-8] > -POLY_A_RNA_THRESHOLD) &&
                    (slope[j-5] < POLY_A_RNA_THRESHOLD && slope[j-9] > -POLY_A_RNA_THRESHOLD) &&
                    (slope[j-5] < POLY_A_RNA_THRESHOLD && slope[j-10] > -POLY_A_RNA_THRESHOLD) &&
                    (slope[j-5] < POLY_A_RNA_THRESHOLD && slope[j-11] > -POLY_A_RNA_THRESHOLD) &&
                    (slope[j-5] < POLY_A_RNA_THRESHOLD && slope[j-12] > -POLY_A_RNA_THRESHOLD) &&
                    (mean_data[j-1] > 0) &&
                    (mean_data[j-2] > 0) &&
                    (mean_data[j-3] > 0) &&
                    (mean_data[j-4] > 0) &&
                    (mean_data[j-5] > 0) &&
                    (mean_data[j-6] > 0) &&
                    (mean_data[j-7] > 0) &&
                    (mean_data[j-8] > 0) &&
                    (mean_data[j-9] > 0) &&
                    (mean_data[j-10] > 0) &&
                    (mean_data[j-11] > 0) &&
                    (mean_data[j-12] > 0) &&
                    (mean_data[j-13] > 0)) {
                    precise_end <- (j+1) * window_size + crude_start
                    break
                }
            }
        },
        error=function(e) e)

        if (save_plots || show_plots) {
            mean_data <- c(rep(NA, times=crude_start),
                           rep(mean_data, each=window_size),
                           rep(NA, times=length(truncated_data)-crude_start-length(mean_data)*window_size))
            slope <- c(rep(NA, times=crude_start),
                           rep(slope, each=window_size),
                           rep(NA, times=length(truncated_data)-crude_start-length(rep(slope, times=window_size))))
        } else {
            mean_data = NA
            slope = NA
        }

    } else {# if no poly(A) tail is found
        if (save_plots || show_plots) {
            mean_data = rep(NA, times=length(truncated_data))
            slope = rep(NA, times=length(truncated_data))
        }
        else {
            mean_data = NA
            slope = NA
        }
    }

    return (list(mean_data=mean_data,
                 slope=slope,
                 start=precise_start,
                 end=precise_end))
}
```
