## Supplementary material for "Nano3P-seq: transcriptome-wide analysis of gene expression and tail dynamics using end-capture nanopore cDNA sequencing": FileS2: find-rna-polya-tail-per-read.R

```
#' Find Poly(A) tail in a single RNA read.
#'
#' @param file_path character string. Path of the FAST5 file
#' @param read_id_fast5_file list
#' @param multifast5 logical
#' @param basecalled_with character string.
#' @param basecall_group character string.
#' @param save_plots logical.
#' @param show_plots logical
#' @param save_dir character string
#' @param plotting_library character string
#' @param plot_debug logical
#' @param model character string
#'
#' @return A list of Fast5 file data
#'
#' @examples
#' \dontrun{
#' find_rna_polya_tail_per_read('path/to/fast5/file')
#' }
#'
find_rna_polya_tail_per_read <- function(file_path = NA,
                                         read_id_fast5_file = NA,
                                         multifast5,
                                         basecalled_with,
                                         basecall_group = 'Basecall_1D_000',
                                         model,
                                         save_plots = FALSE,
                                         show_plots = FALSE,
                                         save_dir = '~',
                                         plotting_library = 'rbokeh',
                                         plot_debug = FALSE) {
    # Empirical parameters
    POLY_A_RNA_THRESHOLD <- 0.3
    POLY_A_RNA_SPIKE_THRESHOLD <- 3.0
    POLY_A_RNA_MOVING_WINDOW_SIZE <- 400
    POLY_A_RNA_ADAPTOR_TROUGH_SIZE_THRESHOLD <- 2200

    # Read the FAST5 data
    if (!save_plots) {
        plot_debug <- FALSE
    }
    read_data <- extract_read_data(file_path,
                                   read_id_fast5_file,
                                   plot_debug,
                                   basecalled_with,
                                   basecall_group,
                                   multifast5,
                                   model,
                                   plotting_library,
                                   experiment_type = 'rna')

    event_data <- read_data$event_data
    raw_data <- read_data$raw_data
    stride <- read_data$stride

    # first read the data and find the tailtype
    if (multifast5) {
        file_path <- read_id_fast5_file$fast5_file
    }
    read_id <- read_data$read_id

    # Z-normalize the data
    norm_data <- z_normalize(raw_data)

    # Recitfy the data (flip anything that is below zero)
    rectified_data <- norm_data

    # Windsorize the data (clip anything above a threshold)
    truncated_data <- truncate_spikes(rectified_data,
                                      spike_threshold=POLY_A_RNA_SPIKE_THRESHOLD)

    # Smoothen the data
    smoothed_data_lr <- left_to_right_sliding_window_rna('mean',
                                                        truncated_data,
                                                        POLY_A_RNA_MOVING_WINDOW_SIZE, 1)
    smoothed_data_rl <- right_to_left_sliding_window_rna('mean',
                                                        truncated_data,
                                                        POLY_A_RNA_MOVING_WINDOW_SIZE, 1)
    smoothed_data <- pmax(smoothed_data_lr, smoothed_data_rl)

    # Find intersections with the threshold
    intersections <- smoothed_data > POLY_A_RNA_THRESHOLD
    rle_intersections <- rle(intersections)

    # Smoothen RLE intersecitons to remove the annoying W pattern formed by the
    # merging the two smoothened signals that have a relative shift with each
    # other.
    rle_intersections <- smoothen_rle_intersections_rna(rle_intersections)
    rle_lengths <- rle_intersections$lengths
    rle_values <- rle_intersections$values
    rle_indices <- cumsum(rle_lengths)

    # Find crude poly(A) boundries
    crude_polya_boundries <- find_rna_polya_crude_start_end(rle_lengths,
                                                            rle_values,
                                                            rle_indices,
                                                            POLY_A_RNA_ADAPTOR_TROUGH_SIZE_THRESHOLD)

    # Refine the curde poly(A) boundries
    precise_polya_boundries <- find_rna_polya_precise_start_end(truncated_data,
                                                                POLY_A_RNA_THRESHOLD,
                                                                crude_polya_boundries,
                                                                save_plots,
                                                                show_plots)
    len_rle <- length(rle_values)
    read_length <- length(read_data$raw_data)

    # Find moves in the poly(A) region
    poly_a_fastq <- NA
    tail_length <- NA
    tail_start <- NA
    tail_end <- NA
    if (!is.na(precise_polya_boundries$start) & !is.na(precise_polya_boundries$end)) {
        # poly_a_fastq <- extract_sequence_between_boundaries(read_data$event_data,
        #                                                     precise_polya_boundries$start,
        #                                                     precise_polya_boundries$end)
        poly_a_fastq <- NA
        tail_start <- precise_polya_boundries$start
        tail_end <- precise_polya_boundries$end

        tail_length <- (tail_end - tail_start) / read_data$samples_per_nt

        if (tail_length > 6) {
            tail_length <-  tail_length - 5 # with the new normalizer
        }
    }

    if (show_plots | save_plots) {
        filename <- paste(read_id, '__', basename(file_path), sep = '')
        plot_title <- paste('Poly(A) tail  |  ',
                            'Tail length [nt]: ', round(tail_length, 2), '  |  ',
                            'Tail start: ', tail_start, '  |  ',
                            'Tail end: ', tail_end, '  |  ',
                            'Tail duration [Sa]: ', tail_end-tail_start, '  |  ',
                            'Samples per nt: ', round(read_data$samples_per_nt, 2),
                            sep='')

        slope <- mean_data <- NULL  # R CMD CHECK
        df = data.frame(x=c(0:(length(read_data$raw_data)-1)),
                        truncated_data=truncated_data,
                        smoothed_data=smoothed_data,
                        moves=read_data$moves_sample_wise_vector + 1,
                        mean_data=precise_polya_boundries$mean_data,
                        slope=precise_polya_boundries$slope)

        # add a poly(A) tail to the dataframe if it exists
        if (!is.na(tail_start) & !is.na(tail_end)) {
            polya_tail <- NULL  # R CMD CHECK
            df['polya_tail'] <- c(rep(NA, times=tail_start-1),
                                  raw_data[tail_start:tail_end],
                                  rep(NA, times=(read_length-tail_end)))
        }

        if (plotting_library == 'ggplot2') {
            p <- ggplot2::ggplot(data=df, ggplot2::aes(x = x))
            if (plot_debug) {
                p <- p + ggplot2::geom_line(ggplot2::aes(y = truncated_data), color='#4040a1') +
                ggplot2::geom_line(ggplot2::aes(y = moves), color = '#b2b2b2') +
                ggplot2::geom_hline(yintercept = POLY_A_RNA_THRESHOLD, color = "#00B0DF") +
                ggplot2::geom_line(ggplot2::aes(y = slope, color = "#BF1268")) +
                ggplot2::geom_hline(yintercept = -POLY_A_RNA_THRESHOLD, color = "#FF94CD")
            } else {
                p <- p + ggplot2::geom_line(ggplot2::aes(y = truncated_data), color='#8E8E8E')
            }

            if (!is.na(tail_start) & !is.na(tail_end)) {
                p <- p + ggplot2::geom_line(ggplot2::aes(y = c(rep(NA, times = tail_start-1),
                                                               truncated_data[tail_start:tail_end],
                                                               rep(NA, times = (read_length-tail_end)))), color='#BF1268')
            }
            if (plot_debug){
                p <- p + ggplot2::geom_line(ggplot2::aes(y = smoothed_data), color='#060B54') +
                ggplot2::geom_line(ggplot2::aes(y = mean_data, color = "#F79A14"))
            }
            p <- p + ggplot2::ggtitle(plot_title) +
                ggplot2::xlab('Sample index') +
                ggplot2::ylab('z-normalized data')
        }

        if (plotting_library == 'rbokeh') {
            p1 <- rbokeh::figure(data=df,
                                 width=1000, height=200,
                                 legend_location="top_right")
            y <- NULL  # R CMD CHECK
            p3 <- rbokeh::figure(data=data.frame(x=event_data$start,
                                                 y=event_data$move/2),
                                 width=1000, height=100,
                                 legend_location="top_right", ygrid = FALSE)
            if (plot_debug)
                p2 <- rbokeh::figure(data=df, width=1000, height=400,
                                     legend_location="top_right")

            p1 <- rbokeh::ly_lines(p1, x=x, y=raw_data, width=1.5, color='#b2b2b2', legend = "Raw data")
            if (!is.na(tail_start) & (!is.na(tail_end))) {
                p1 <- rbokeh::ly_lines(p1, x=x, y=polya_tail, width=1.5, color = '#ea3e13', legend = "Poly(A) tail")
            }
            p1 <- rbokeh::y_axis(p1, label='pA', num_minor_ticks=2)
            p1 <- rbokeh::x_axis(p1, label='Sample index')
            p1 <- rbokeh::tool_pan(p1, dimensions = "width")
            p1 <- rbokeh::tool_wheel_zoom(p1, dimensions = "width")

            # plot containing all the debug traces
            if (plot_debug) {
                p2 <- rbokeh::ly_lines(p2, x=x, y=truncated_data, width=1.0, color='#b2b2b2', legend = "Normalized windsorized signal")
                p2 <- rbokeh::ly_lines(p2, x=x, y=smoothed_data, color='#8c8a8a', width=1.5, legend = "Smoothed signal")
                p2 <- rbokeh::ly_lines(p2, x=x, y=slope, color='#f3c963', width=2, legend = "Slope of the signal in the search window")
                p2 <- rbokeh::ly_lines(p2, x=x, y=mean_data, color='#f58585', width=2, legend = "Mean of the signal in the search window")
                #p2 <- rbokeh::ly_lines(p2, moves, color='#b2b2b2', width=1, legend = "Moves")
                p2 <- rbokeh::ly_abline(p2, h=POLY_A_RNA_THRESHOLD, color = '#c581f5', type = 3, width=2, legend = "Slope upper bound")
                p2 <- rbokeh::ly_abline(p2, h=-POLY_A_RNA_THRESHOLD, color = '#ff6e24', width=2, type = 3, legend = "Slope lower bound")
                if (!is.na(tail_start) & (!is.na(tail_end))) {
                    p2 <- rbokeh::ly_abline(p2, v=tail_start, color = '#4497b9', width=2, legend = "Tail start")
                    p2 <- rbokeh::ly_abline(p2, v=tail_end, color = '#879833', width=2, legend = "Tail end")
                }
                p2 <- rbokeh::y_axis(p2, label='z-normalized data values', num_minor_ticks=4, desired_num_ticks = 5)
                p2 <- rbokeh::x_axis(p2, label='Sample index')
                p2 <- rbokeh::tool_pan(p2, dimensions = "width")
                p2 <- rbokeh::tool_wheel_zoom(p2, dimensions = "width")
            }

            # plot containing the moves
            p3 <- rbokeh::ly_crect(p3,
                                   x=x,
                                   y=y,
                                   width=rep(0.01, nrow(event_data)),
                                   height=event_data$move,
                                   color='#529c82')
            p3 <- rbokeh::y_axis(p3, label='', num_minor_ticks=0, desired_num_ticks = 3)
            p3 <- rbokeh::x_axis(p3, label='Sample index')
            p3 <- rbokeh::tool_pan(p3, dimensions = "width")
            p3 <- rbokeh::tool_wheel_zoom(p3, dimensions = "width")
            p3 <- rbokeh::tool_wheel_zoom(p3, dimensions = "height")

            if (plot_debug){
                lst <- list(p1, p2, p3)
                names(lst) <- c(plot_title, 'Debugging Traces', 'Moves')
                nrow <- 3
            } else {
                lst <- list(p1, p3)
                names(lst) <- c(plot_title, 'Moves')
                nrow <- 2
            }
            suppressWarnings({
            p <- rbokeh::grid_plot(lst,
                                   nrow = nrow,
                                   link_data = TRUE,
                                   same_axes=c(TRUE, FALSE))
            })
        }

        if (save_plots) {
            filename_png <- paste(filename,'.png', sep='')
            filename_html <- paste(filename,'.html', sep='')
            save_path_png <- file.path(save_dir, 'plots', filename_png, fsep = .Platform$file.sep)
            save_path_html <- file.path(save_dir, 'plots', filename_html, fsep = .Platform$file.sep)
            if (plotting_library == 'ggplot2') {
                ggplot2::ggsave(save_path_png, plot = p, width = 300, height = 70, units = 'mm')
            }
            else {
                #rbokeh::widget2png(p, file = save_path_png)
                rbokeh::rbokeh2html(p, file = save_path_html)
                # Hack to make the plots working
                tx  <- readLines(save_path_html)
                tx2  <- gsub(pattern="bokeh-0.12.5", replace="bokeh-0.12.3", x=tx)
                writeLines(tx2, con=save_path_html)
            }
        }

        if (show_plots) {
            print(p)
        }
    }

    data <- list(read_id = read_id,
                 tail_start = precise_polya_boundries$start,
                 tail_end = precise_polya_boundries$end,
                 samples_per_nt = read_data$samples_per_nt,
                 tail_length = tail_length,
                 polya_fastq = poly_a_fastq,
                 file_path = file_path)
    return(data)
}
```
