## Supplementary material for "Nano3P-seq: transcriptome-wide analysis of gene expression and tail dynamics using end-capture nanopore cDNA sequencing": FileS2: find-tails.R

```
#' Estimates poly(A)/(T) tail lengths in Oxford Nanopore RNA and DNA reads
#'
#' This function estimates poly(A) tail length in RNA reads, and both poly(A)
#' and poly(T) tail lengths in DNA reads. It can operate on reads base called
#' with any version of Albacore and Guppy using either the standard or the
#' recent 'flip-flop' model. The function outputs a CSV file containing poly(A)
#' tail information organised by the read ID; it also returns the same
#' information as a tibble for further processing by the end-user. Currently,
#' the algorithm works only on ONT 1D reads.
#'
#' @param fast5_dir character string. Full path of the directory to search the
#' basecalled fast5 files in. The Fast5 files can be single or multi-fast5 file.
#' The directory is searched recursively.
#'
#' @param save_dir character string. Full path of the directory where the CSV
#' file containing the tail-length information should be stored. If save_plots
#' is set to \code{TRUE}, then a \code{plots} directory is also created within
#' the \code{save_dir}.
#'
#' @param csv_filename character string ["tails.csv"]. Filename of the
#' CSV file in which to store the tail length data
#'
#' @param num_cores numeric [1]. Num of physical cores to use in processing
#' the data. Always use 1 less than the number of cores at your disposal.
#'
#' @param basecall_group a character string ["Basecall_1D_000"]. Name of the
#' level in the Fast5 file hierarchy from which tailfindr should read the data.
#'
#' @param save_plots logical [FALSE]. If set to \code{TRUE}, a plots
#' directory will be created within the save_dir, and plots showing poly(A) and
#' poly(T) tails in the raw squiggle will be saved in this \code{plots}
#' directory. Creating plots and saving them to the disk is a slow process. We
#' recommend that you keep this option set to \code{FALSE}. If you still want
#' to create plots, we recommend that you run tailfindr on a subset of reads.
#' Plots are automatically named by concatenating read ID with the name of the
#' Fast5 file containing this read; the read ID and fast5 file name are
#' separated by two underscores (__).
#'
#' @param plot_debug_traces logical [FALSE]. This option works only
#' if \code{save_plots} option is also set to \code{TRUE}.If set to \code{TRUE},
#' debugging information is plotted in the plots as well. This includes mean
#' signal, slope signal,thresholds, smoothened signal, etc. We use this option
#' internally to debug our algorithm.
#'
#' @param plotting_library character string ["rbokeh"]. \code{rbokeh}
#' is the default plotting library used if \code{save_plots} is set to
#' \code{TRUE}. The plots will be saved as HTML files in the
#' \code{/save_dir/plots} directory. You can open these HTLM files in any
#' web-browser and interactively view the plots showing the tail region in the
#' raw squiggle. If this option is set to \code{'ggplot2'}, then the polts will
#' be saved as static \code{.png} files.
#'
#' @param ... list. A list of optional parameters. This is currently, reserved
#' for internal use only.
#'
#' @return A data tibble containing tail information organzied by
#' the read ID is returned. Always save this returned tibble in a variable (see
#' examples below), otherwise the long tibble will be printed to the
#' console, which may hang up your R session.
#'
#' A CSV file containing the same information is also saved on disk in the
#' \code{save_dir}.
#'
#' @export
#'
#' @examples
#' \dontrun{
#'
#' library(tailfindr)
#'
#' # 1. Suppose you have 11 cores at your disposal, then you should run tailfindr
#' # on your data as following:
#' df <- find_tails(fast5_dir = system.file('extdata', 'rna', package = 'tailfindr'),
#'                  save_dir = '~/Downloads',
#'                  csv_filename = 'rna_tails.csv',
#'                  num_cores = 10)
#' # In the above example, we have  used tailfindr on example RNA reads
#' # present in the tailfindr package. You should substitute the path of
#' # your data for the fast5_dir parameter.
#'
#' # 2. If you want to save interactive HTML plots using rbokeh,
#' # then you should run tailfindr as following:
#' df <- find_tails(fast5_dir = system.file('extdata', 'cdna', package = 'tailfindr'),
#'                  save_dir = '~/Downloads',
#'                  csv_filename = 'cdna_tails.csv',
#'                  num_cores = 10,
#'                  save_plots = TRUE,
#'                  plotting_library = 'rbokeh')
#'
#' # 3. If you also want to plot debug traces, then you should run tailfindr as
#' # below:
#' df <- find_tails(fast5_dir = system.file('extdata', 'cdna', package = 'tailfindr'),
#'                  save_dir = '~/Downloads',
#'                  csv_filename = 'cdna_tails.csv',
#'                  num_cores = 10,
#'                  save_plots = TRUE,
#'                  plot_debug_traces = TRUE,
#'                  plotting_library = 'rbokeh')
#'
#' # N.B.: Making and saving plots is a computationally slow process.
#' # Only generate plots by running tailfindr on a small subset of your reads.
#'
#' # 4. By default, tailfindr uses Events/Move table in the Basecall_1D_000
#' # section of the FAST5 file. If you want tailfindr to pick Events/Move table
#' # from some other section of the FAST5 file -- lets say Basecall_1D_001--
#' # then you should use tailfindr like below:
#' df <- find_tails(fast5_dir = system.file('extdata', 'rna_basecall_1D_001', package = 'tailfindr'),
#'                  save_dir = '~/Downloads',
#'                  csv_filename = 'rna_tails.csv',
#'                  num_cores = 2,
#'                  basecall_group = 'Basecall_1D_001',
#'                  save_plots = TRUE,
#'                  plot_debug_traces = TRUE,
#'                  plotting_library = 'rbokeh')
#' # N.B.: tailfindr cannot work if it can't find Events or Move table in
#' # your FAST5 files. MinKNOW Live Basecalling currently does not save the
#' # Events/Move table in the FAST5 file. If your reads have been live
#' # basecalled, then you should rebasecall them using Albacore or Guppy, and
#' # subsequently use tailfindr and specify the basecall_group parameter. Most
#' # probably, in the second round of your basecalling, the Events/Move table
#' # is stored in the 'Basecall_1D_001' section, so set this as the value of the
#' # basecall_group parameter. You can also confirm this by viewing your
#' # re-basecalled reads in HDFView.
#' }
#'
find_tails <- function(fast5_dir,
                       save_dir,
                       csv_filename = 'tails.csv',
                       num_cores = 1,
                       basecall_group = 'Basecall_1D_000',
                       save_plots = FALSE,
                       plot_debug_traces = FALSE,
                       plotting_library = 'rbokeh',
                       ...) {

    plot_debug <- plot_debug_traces
    if (save_plots == FALSE) {
        plot_debug <- FALSE
    }
    # Taking out these parameter from the function parameters list
    # as they may be dangerous for normal users
    show_plots <- FALSE

    dna_datatype <- get_dna_datatype(...)  # R CMD NOTE
    if (dna_datatype == 'custom-cdna') {
        fp_ep_list <- get_custom_fp_ep(...)  # R CMD NOTE
    }

    # start a log file
    if (dir.exists(file.path(save_dir))) {
        logfile_name <- paste(format(Sys.time(), "%Y-%m-%d_%H-%M-%S"), "_tailfinder.log", sep = "")
        logfile_path <- file.path(save_dir, logfile_name, fsep = .Platform$file.sep)
        con <- file(logfile_path, open = "a")
        sink(con, append=TRUE, split = TRUE, type='output')
        on.exit(sink(file=NULL, type = 'output'))
    }

    # display console messages
    version <- utils::packageDescription("tailfindr")$Version
    cat(cli::rule(left=''), '\n', sep = "")
    cat(cli::rule(left=paste("Started tailfindr ", '(version ', version, ')', sep='')), '\n', sep = "")
    cat(cli::rule(left=''), '\n',  sep = "")

    # display the user-specified parameters
    cat(paste(cli::symbol$menu, ' You have configured tailfindr as following:', '\n', sep=''))
    cat(paste(cli::symbol$pointer, ' fast5_dir:         ', fast5_dir, '\n', sep=''))
    cat(paste(cli::symbol$pointer, ' save_dir:          ', save_dir, '\n', sep=''))
    cat(paste(cli::symbol$pointer, ' csv_filename:      ', csv_filename, '\n', sep=''))
    cat(paste(cli::symbol$pointer, ' num_cores:         ', num_cores, '\n', sep=''))
    cat(paste(cli::symbol$pointer, ' basecall_group:    ', basecall_group, '\n', sep=''))
    cat(paste(cli::symbol$pointer, ' save_plots:        ', save_plots, '\n', sep=''))
    cat(paste(cli::symbol$pointer, ' plot_debug_traces: ', plot_debug_traces, '\n', sep=''))
    cat(paste(cli::symbol$pointer, ' plotting_library:  ', plotting_library, '\n', sep=''))
    if (dna_datatype == 'pcr-dna' | dna_datatype == 'custom-cdna') {
        cat(paste(cli::symbol$pointer, ' dna_datatype:      ', dna_datatype, '\n', sep=''))
    }

    cat(cli::rule(left=paste('Processing started at ', Sys.time(), sep = '')), '\n', sep = "")

    # Try to create the save directory
    if (!dir.exists(file.path(save_dir))) {
        cat(paste(cli::symbol$bullet, ' Save dir does not exist. Trying to create it...\n', sep=''))
        tryCatch({
            dir.create(file.path(save_dir, fsep = .Platform$file.sep))
            cat('  Done!\n')
        },
        error=function(e){
            cat(paste(cli::symbol$bullet, ' Failed to create the save dir. Results will be stored in the "~/" directory instead.\n', sep=''))
            save_dir <- '~/'
        })
        logfile_name <- paste(format(Sys.time(), "%Y-%m-%d_%H-%M-%S"), "_tailfinder.log", sep = "")
        logfile_path <- file.path(save_dir, logfile_name, fsep = .Platform$file.sep)
        con <- file(logfile_path, open = "a")
        sink(con, append=TRUE, split = TRUE, type='output')
        on.exit(sink(file=NULL, type = 'output'))
    }

    # Create a sub-direcotry to save all the plots
    if (save_plots){
        plots_dir <- file.path(save_dir, 'plots', fsep = .Platform$file.sep)
        if (!dir.exists(file.path(plots_dir))) {
            cat(paste(cli::symbol$bullet, ' Creating a sub-directory to save the plots in.\n', sep=''))
            dir.create(plots_dir)
            cat('  Done! All plots will be saved in the following direcotry:\n')
            cat(paste('  ', file.path(save_dir, 'plots', fsep = .Platform$file.sep), '\n', sep = ''))
        }
    }

    # search for all the fast5 files in the user-specified directory
    cat(paste(cli::symbol$bullet,' Searching for all Fast5 files...\n', sep=''))

    # must handle fast5 dir, or a pre-made character vector of file paths
    if (is.na(fast5_dir[2])) {
        fast5_files_list <- list.files(path = fast5_dir,
                                       pattern = "\\.fast5$",
                                       recursive = TRUE,
                                       full.names = TRUE)
    } else {
        fast5_files_list <- fast5_dir
    }

    num_files <- length(fast5_files_list)
    cat(paste0('  Done! Found ', num_files, ' Fast5 files.\n'))

    # read the first read in the list of reads,
    # and determine all the properties of the data
    cat(paste(cli::symbol$bullet,' Analyzing a single Fast5 file to assess if your data \n', sep=''))
    cat('  is in an acceptable format...\n')

    type_info <- explore_basecaller_and_fast5type(fast5_files_list[1],
                                                  basecall_group = basecall_group)

    # currently MinKNOW does not store Events/Move table without which we
    # cannot compute the read-specific normalizer
    if (type_info$basecalled_with == 'minknow' & type_info$model == 'unknown') {
        cat(paste0('  ', cli::symbol$cross,
                  ' Fatal error! Your data has been basecalled with MinKNOW\n'))
        cat('    live basecalling which currently does not save the\n')
        cat('    Events/Move table in the Analyses/Basecall_1D_000 section of\n')
        cat('    the FAST5 file. You should rebasecall your FAST5 files using \n')
        cat('    standalone Guppy or Albacore, and then use tailfindr on the \n')
        cat('    rebasecalled files. Please adjust the value of basecall_group\n')
        cat('    parameter in such a case, so that tailfindr can find the \n')
        cat('    Events/Move table in the specified basecall_group. You can\n')
        cat('    check which basecall_group the Event/Move is residing by\n')
        cat('    opening your FAST5 file in HDFView.\n\n')
        cat('    If the Events/Move is present in the data and you have\n')
        cat('    specified the correct basecall_group, but you still\n')
        cat('    get this error then please open an issue on GitHub:\n')
        cat('    https://github.com/adnaniazi/tailfindr/issues\n')
        cat('    Remember to attach a few (around 5) of your FAST5 files\n')
        cat('    to help us understand the issue.\n')
        cat(cli::rule(left=paste('Processing ended at ',
                                 Sys.time(), sep = '')), '\n', sep = "")
        cat(paste(crayon::green(cli::symbol$cross),
                  ' tailfindr finished unsuccessfully!\n', sep=''))
        return(0)
    }

    basecalled_with <- type_info$basecalled_with
    multifast5 <- ifelse(type_info$fast5type == 'multi', TRUE, FALSE)
    experiment_type <- type_info$experiment_type
    read_is_1d <- type_info$read_is_1d
    model <- type_info$model
    if (basecalled_with == 'albacore'){
        cat(paste('  ', crayon::green(cli::symbol$tick),
                  ' The data has been basecalled using Albacore.\n', sep=''))
    } else {
        cat(paste('  ', crayon::green(cli::symbol$tick),
                  ' The data has been basecalled using Guppy.\n', sep=''))
    }
    if (model == 'flipflop'){
        cat(paste('  ', crayon::green(cli::symbol$tick),
                  ' Flipflop model was used during basecalling.\n', sep=''))
    } else {
        cat(paste('  ', crayon::green(cli::symbol$tick),
                  ' Standard model was used during basecalling.\n', sep=''))
    }
    if (multifast5){
        cat(paste('  ', crayon::green(cli::symbol$tick),
                  ' The reads are packed in multi-fast5 file(s).\n', sep=''))
    } else {
        cat(paste('  ', crayon::green(cli::symbol$tick),
                  ' Every read is in a single fast5 file of its own.\n', sep=''))
    }
    if (experiment_type == 'rna'){
        cat(paste('  ', crayon::green(cli::symbol$tick),
                  ' The experiment type is RNA, so we will search\n', sep=''))
        cat('    for poly(A) tails.\n')
    } else {
        cat(paste('  ', crayon::green(cli::symbol$tick),
                  ' The experiment type is DNA (Nano3P-seq), so we \n', sep=''))
        cat('    search for reads with and without poly(T) tails.\n')
    }
    if (read_is_1d == TRUE){
        cat(paste('  ', crayon::green(cli::symbol$tick),
                  ' The reads are 1D reads.\n', sep=''))
    } else {
        cat(paste('  ', crayon::red(cli::symbol$cross),
                  ' The reads are not 1D. Currently, we only support\n', sep=''))
        cat('    1D reads. If you believe your reads are 1D, and you are\n')
        cat('    getting this cat erroneously, please feel free\n')
        cat('    to contact us at. Do not forget to\n')
        cat('    send us one of the problematic reads so that we can\n')
        cat('    debug our software, and send you a patch.\n')
        cat(paste('  ', crayon::red(cli::symbol$cross),
                  'Finished because of the error!\n', sep=''))
        cat(cli::rule(left=paste('tailfindr finished with a fatal error at ',
                                 Sys.time(), sep = '')), '\n', sep = "")
        return(0)
    }

    # Make a compute cluster
    cat(paste(cli::symbol$bullet,' Starting a parallel compute cluster...\n', sep=''))
    #cl <- parallel::makeCluster(num_cores, outfile='')
    cl <- parallel::makeCluster(num_cores)
    on.exit(parallel::stopCluster(cl))

    doSNOW::registerDoSNOW(cl)
    `%dopar%` <- foreach::`%dopar%`
    `%do%` <- foreach::`%do%`
    cat('  Done!\n')
    mcoptions <- list(preschedule = TRUE, set.seed = FALSE, cleanup = FALSE)

    # if the data is DNA then make a substitution matrix
    if (experiment_type == 'dna') {
        match <- 1
        mismatch <- -1
        type <- 'local'
        gapOpening <- 0
        gapExtension <- 1
        submat <- Biostrings::nucleotideSubstitutionMatrix(match = match,
                                                           mismatch = mismatch,
                                                           baseOnly = TRUE)
        dna_opts <- list(match = match,
                         mismatch = mismatch,
                         type = type,
                         gapOpening = gapOpening,
                         gapExtension = gapExtension,
                         submat = submat)

        # append custom front and end primers to DNA options if it is custom
        # cDNA
        if (dna_datatype == 'custom-cdna') {
            dna_opts <- c(dna_opts, fp_ep_list)
        }
    }

    # If the fast5 are multifast5, then build an index of all the reads within these files
    remove_last_duplicate_read <- FALSE
    if (multifast5) {
        cat(paste(cli::symbol$bullet, ' Discovering reads in the ',
                  num_files, ' multifast5 files...\n', sep=''))
        read_id_fast5_file <- dplyr::tibble(read_id = character(),
                                            fast5_file = character())
        for (fast5_file in fast5_files_list) {
            f5_obj <- hdf5r::H5File$new(fast5_file, mode = 'r')
            f5_tree <- f5_obj$ls(recursive = FALSE)
            f5_tree <- f5_tree$name
            f5_tree <- dplyr::mutate(dplyr::tibble(f5_tree), fast5_file = fast5_file)
            value <- NULL  # R CMD CHECK
            f5_tree <- dplyr::rename(f5_tree, read_id = f5_tree)
            read_id_fast5_file <- rbind(read_id_fast5_file, f5_tree)
            f5_obj$close_all()
        }
        cat(paste0('  Done! Found ', nrow(read_id_fast5_file), ' reads\n'))
        # convert the data frame to list with rows as elements of the list
        read_id_fast5_file <- split(read_id_fast5_file, seq(nrow(read_id_fast5_file)))

        # Split the data into chunks
        files_per_chunk <- 4000
        total_files <- length(read_id_fast5_file)
        total_chunks <- ceiling(total_files/files_per_chunk)

        #loop
        if (experiment_type == 'dna') {
            cat(paste(cli::symbol$bullet,
                      ' Searching for reads with and without Poly(T) tails...\n', sep=''))
        } else {
            cat(paste(cli::symbol$bullet,
                      ' Searching for Poly(A) tails...\n', sep=''))
        }
        counter <- 0
        result <- list()
        for(chunk in seq_len(total_chunks)){
            # divide data in chunks
            if(chunk == total_chunks) {
                read_id_fast5_file_subset <-
                    read_id_fast5_file[((counter*files_per_chunk)+1):total_files]
                # if the last chunk has only one read then just duplicate this
                # one read so that the progressbar works, and does not throw an error
                if (length(read_id_fast5_file_subset) == 1) {
                    read_id_fast5_file_subset[[2]] <- read_id_fast5_file_subset
                    remove_last_duplicate_read <- TRUE
                }
            } else {
                read_id_fast5_file_subset <-
                    read_id_fast5_file[((counter*files_per_chunk)+1):((counter+1)*files_per_chunk)]
            }
            counter <- counter + 1
            cat(paste('  Processing chunk ', chunk, ' of ', total_chunks, '\n', sep = ''))

            # progress bar
            pb <- utils::txtProgressBar(min = 1,
                                        max = length(read_id_fast5_file_subset),
                                        style = 3)
            progress <- function(n) utils::setTxtProgressBar(pb, n)
            opts <- list(progress = progress)

            # foreach loop
            sink(file=NULL, type = 'output')
            close(con)
            if (experiment_type == 'dna') {
                riff <- NULL  # R CMD CHECK
                data_list <- foreach::foreach(riff = read_id_fast5_file_subset,
                                              .combine = 'rbind',
                                              .inorder = FALSE,
                                              .errorhandling = 'pass',
                                              .options.snow = opts,
                                              .options.multicore = mcoptions) %dopar% {
                                                  tryCatch({
                                                      find_dna_tail_per_read(read_id_fast5_file = riff,
                                                                             file_path = NA,
                                                                             basecall_group = basecall_group,
                                                                             dna_datatype = dna_datatype,
                                                                             save_plots = save_plots,
                                                                             show_plots = show_plots,
                                                                             plot_debug = plot_debug,
                                                                             save_dir = save_dir,
                                                                             plotting_library = plotting_library,
                                                                             multifast5 = multifast5,
                                                                             basecalled_with = basecalled_with,
                                                                             model = model,
                                                                             dna_opts = dna_opts)
                                                  },
                                                  error=function(e){
                                                      ls <- list(read_id = riff$read_id,
                                                                 read_type = NA,
                                                                 tail_start = NA,
                                                                 tail_end = NA,
                                                                 samples_per_nt = NA,
                                                                 tail_length = NA,
                                                                 file_path = riff$fast5_file,
                                                                 has_precise_boundary = NA)
                                                  })
                                              }
            } else {
                riff <- NULL  # R CMD CHECK
                data_list <- foreach::foreach(riff = read_id_fast5_file_subset,
                                              .combine = 'rbind',
                                              .inorder = FALSE,
                                              .errorhandling = 'pass',
                                              .options.snow = opts,
                                              .options.multicore = mcoptions) %dopar% {
                                                  tryCatch({
                                                      find_rna_polya_tail_per_read(file_path = NA,
                                                                                   read_id_fast5_file = riff,
                                                                                   basecall_group = basecall_group,
                                                                                   multifast5 = multifast5,
                                                                                   basecalled_with = basecalled_with,
                                                                                   model = model,
                                                                                   save_plots = save_plots,
                                                                                   show_plots = show_plots,
                                                                                   save_dir = save_dir,
                                                                                   plotting_library = plotting_library,
                                                                                   plot_debug = plot_debug)
                                                  },
                                                  error=function(e){
                                                      ls <- list(read_id = NA,
                                                                 tail_start = NA,
                                                                 tail_end = NA,
                                                                 samples_per_nt = NA,
                                                                 tail_length = NA,
                                                                 polya_fastq = NA,
                                                                 file_path = riff$fast5_file)
                                                  })
                                              }

            }
            cat('\n')
            if (remove_last_duplicate_read) data_list <- data_list[[1]]
            result[[chunk]] <- data_list
            con <- file(logfile_path, open = "a")
            sink(con, append=TRUE, split = TRUE, type='output')
        }
    } else if (!multifast5) {
        # Split the data into chunks
        files_per_chunk <- 4000
        total_files <- length(fast5_files_list)
        total_chunks <- ceiling(total_files/files_per_chunk)

        counter <- 0
        result <- list()

        if (experiment_type == 'dna') {
            cat(paste(cli::symbol$bullet,
                      ' Searching for Poly(A) and Poly(T) tails...\n', sep=''))
        } else {
            cat(paste(cli::symbol$bullet,
                      ' Searching for Poly(A) tails...\n', sep=''))
        }

        for (chunk in seq_len(total_chunks)) {
            if(chunk == total_chunks) {
                fast5_files_subset <-
                    fast5_files_list[((counter*files_per_chunk)+1):total_files]
                # if the last chunk has only one read then just duplicate this
                # one read so that the progressbar works and does not throw an error
                if (length(fast5_files_subset) == 1) {
                    fast5_files_subset[[2]] <- fast5_files_subset
                    remove_last_duplicate_read <- TRUE
                }
            } else {
                fast5_files_subset <-
                    fast5_files_list[((counter*files_per_chunk)+1):((counter+1)*files_per_chunk)]
            }
            counter <- counter + 1

            # progress bar
            cat(paste('  Processing chunk ', chunk, ' of ', total_chunks, '\n', sep=''))
            pb <- utils::txtProgressBar(min = 1,
                                        max = length(fast5_files_subset),
                                        style = 3)
            progress <- function(n) utils::setTxtProgressBar(pb, n)
            opts <- list(progress = progress)

            # foreach loop
            sink(file=NULL, type = 'output')
            close(con)
            if (experiment_type == 'dna') {
                file_path <- NULL  # R CMD CHECK
                data_list <- foreach::foreach(file_path = fast5_files_subset,
                                              .combine = 'rbind',
                                              .inorder = FALSE,
                                              .options.snow = opts,
                                              .options.multicore = mcoptions) %dopar% {
                                                 tryCatch({
                                                     find_dna_tail_per_read(file_path = file_path,
                                                                            basecall_group = basecall_group,
                                                                            dna_datatype = dna_datatype,
                                                                            save_plots = save_plots,
                                                                            show_plots = show_plots,
                                                                            plot_debug = plot_debug,
                                                                            save_dir = save_dir,
                                                                            plotting_library = plotting_library,
                                                                            multifast5 = multifast5,
                                                                            basecalled_with = basecalled_with,
                                                                            model = model,
                                                                            dna_opts = dna_opts)
                                                 },
                                                 error=function(e){
                                                     ls <- list(read_id = NA,
                                                                read_type = NA,
                                                                tail_start = NA,
                                                                tail_end = NA,
                                                                samples_per_nt = NA,
                                                                tail_length = NA,
                                                                file_path = file_path,
                                                                has_precise_boundary = NA)
                                                 })
                                             }
            } else {
                file_path <- NULL  # R CMD CHECK
                data_list <-foreach::foreach(file_path = fast5_files_subset,
                                             .combine = 'rbind',
                                             .inorder = FALSE,
                                             .options.snow = opts,
                                             .options.multicore = mcoptions) %dopar% {
                                                 tryCatch({
                                                     find_rna_polya_tail_per_read(file_path = file_path,
                                                                                  read_id_fast5_file = NA,
                                                                                  basecall_group = basecall_group,
                                                                                  multifast5 = multifast5,
                                                                                  basecalled_with = basecalled_with,
                                                                                  model = model,
                                                                                  save_plots = save_plots,
                                                                                  show_plots = show_plots,
                                                                                  save_dir = save_dir,
                                                                                  plotting_library = plotting_library,
                                                                                  plot_debug = plot_debug)
                                                 },
                                                 error=function(e){
                                                     ls <- list(read_id = NA,
                                                                tail_start = NA,
                                                                tail_end = NA,
                                                                samples_per_nt = NA,
                                                                tail_length = NA,
                                                                polya_fastq = NA,
                                                                file_path = file_path)
                                                 })
                                             }
            }
            cat('\n')
            if (remove_last_duplicate_read) data_list <- data_list[[1]]
            result[[chunk]] <- data_list
            con <- file(logfile_path, open = "a")
            sink(con, append=TRUE, split = TRUE, type='output')
        }
    }

    # format the results list into a tibble
    cat(paste0(cli::symbol$bullet,' Formatting the tail data...\n'))
    result <- purrr::map(result, function(.x) tibble::as_tibble(.x))
    result <- dplyr::bind_rows(result, .id = "chunk")
    result <- dplyr::select(result, -chunk)
    # cleanup the tibble
    result <- tidyr::unnest(result)
    if (experiment_type == 'dna') {
        has_precise_boundary <- NULL  # R CMD CHECK
        result <- within(result, rm(has_precise_boundary))
    } else {
        polya_fastq <- NULL  # R CMD CHECK
        result <- within(result, rm(polya_fastq))
    }
    result$tail_length <- round(result$tail_length, digits = 2)
    result$samples_per_nt <- round(result$samples_per_nt, digits = 2)
    cat('  Done!\n')

    # write the result to a csv file
    cat(paste(cli::symbol$bullet,
              ' Saving the data in the CSV file...\n', sep=''))
    data.table::fwrite(result, file.path(save_dir, csv_filename, fsep = .Platform$file.sep))
    cat('  Done! Below is the path of the CSV file:\n')
    cat(paste0('  ', file.path(save_dir, csv_filename, fsep = .Platform$file.sep), '\n'))


    cat(paste0(cli::symbol$bullet,
               ' A logfile containing all this information has been saved in this path: \n'))
    cat(paste0('  ', logfile_path, '\n'))

    cat(cli::rule(left=paste('Processing ended at ',
                             Sys.time(), sep = '')), '\n', sep = "")
    cat(paste(crayon::green(cli::symbol$tick),
              ' tailfindr finished successfully!\n', sep=''))
    return(result)

    # close logfile connection
    close(con)
}
```
