## Supplementary material for "Nano3P-seq: transcriptome-wide analysis of gene expression and tail dynamics using end-capture nanopore cDNA sequencing": FileS2: find_sample_index_for_fastq_base.R

```
#' Finds raw data sample index corresponding to a base.
#'
#' Finds raw data sample index in the event data table corresponding to a FastQ
#' base number. The closet possible matching sample is dependent on the type of
#' the tail.
#'
#' @param event_data dataframe
#' @param fastq_base_number numeric
#' @param read_type character string
#'
#' @return numeric
#'
#' @examples
#' \dontrun{
#' event_data <- data.frame(start = seq(0, 150, 5),
#'                          move = 5,
#'                          model_state = sample(c("A", "T", "C", "G"), 1))
#' idx <- find_sample_index_for_fastq_base(event_data, 10, 'polyA')
#' }
#'
find_sample_index_for_fastq_base <- function(event_data, fastq_base_number, read_type){
    model_state_length <- nchar(event_data$model_state[1])
    cumm_sum <- cumsum(event_data$move) + model_state_length - 1
    event_data <- cbind(event_data, cumm_sum)

    # make a cummulative sum vector
    if (read_type=='polyT') {
        # for polyT reads take the first move that is
        # equal to the fastq_base_number
        row_index <- min(which(cumm_sum==fastq_base_number))
        if (row_index==-Inf | row_index==Inf) {
            row_index <- min(which(cumm_sum==(fastq_base_number-1)))
        }
    } else {
        # for polyA reads take the last move that is
        # equal to the fastq_base_number
        fastq_base_number <- fastq_base_number + model_state_length # we want the tail to have fully passed through the pore
        row_index <- min(which(cumm_sum==fastq_base_number))
        if (row_index==-Inf | row_index==Inf) {
            row_index <- min(which(cumm_sum==fastq_base_number+1))
        }
    }
    sample_index <- event_data$start[row_index]
    return(sample_index)
}
```
