## Supplementary material for "Nano3P-seq: transcriptome-wide analysis of gene expression and tail dynamics using end-capture nanopore cDNA sequencing": FileS2: get-custom-fp-ep.R

```
#' Get custom front and end primers from optional parameters list
#'
#' @param ... Optional paramters list
#'
#' @return A list of front and end primers

get_custom_fp_ep <- function(...){
    if (length(list(...)) > 0) {
        opt_params <- list(...)
        fp <- opt_params$front_primer
        ep <- opt_params$end_primer
    } else {
        cat('Error! You must specify custom front primer (fp) and end primer (ep) sequences')
    }
    return(list(fp = fp, ep = ep))
}
```
