## Supplementary material for "Nano3P-seq: transcriptome-wide analysis of gene expression and tail dynamics using end-capture nanopore cDNA sequencing": FileS2: get-dna-datatype.R

```
#' Parse optional params to get the type of DNA
#'
#' @param ... Optional parameters
#'
#' @return a character sting
#'
get_dna_datatype <- function(...){
    if (length(list(...)) > 0) {
        opt_params <- list(...)
        dna_datatype <- opt_params$dna_datatype
    } else {
        dna_datatype <- 'cdna'
    }
    return(dna_datatype)
}
```
