## Supplementary material for "Nano3P-seq: transcriptome-wide analysis of gene expression and tail dynamics using end-capture nanopore cDNA sequencing": FileS2: read-sam.R

```
#' Read SAM file
#' Author credits: Grischa Toedt & VladaMilch
#'
#' @param sam_file_path The path of the SAM file
#'
#' @return a list
#'
read_sam <- function(sam_file_path){
    message("Reading SAM file...")
    x = readLines(sam_file_path)
    message("Fetching header...")
    headerpos = grep("^@", x)
    header = x[headerpos]
    message("Converting header...")

    header = list("HD" = lapply(gsub("^@HD\t", "", header[grep("^@HD", header)]),
                                function(x)strsplit(x, "\t")[[1]]),
                  "SQ" = lapply(gsub("^@SQ\t", "", header[grep("^@SQ", header)]),
                                function(x)strsplit(x,"\t")[[1]]),
                  "RG" = lapply(gsub("^@RG\t", "", header[grep("^@RG", header)]),
                                function(x)strsplit(x, "\t")[[1]]),
                  "PG" = lapply(gsub("^@PG\t", "", header[grep("^@PG", header)]),
                                function(x)strsplit(x, "\t")[[1]]),
                  "CO" = gsub("^@CO\t", "", header[grep("^@CO", header)])
    )

    message("Fetching data...")
    x = lapply(x[-headerpos], line2vals)
    message("Convert to data.frame...")
    x = matrix(unlist(x),
               nrow = length(x),
               ncol = length(x[[1]]),
               byrow = TRUE,
               dimnames = list(NULL,c("QNAME","FLAG","RNAME","POS","MAPQ",
                                      "CIGAR","RNEXT","PNEXT","TLEN","SEQ",
                                      "QUAL","ATTR")))
    x = as.data.frame(x,stringsAsFactors = FALSE)
    x$FLAG <- as.integer(x$FLAG)
    x$POS <- as.integer(x$POS)
    x$MAPQ <- as.integer(x$MAPQ)
    x$PNEXT <- as.integer(x$PNEXT)
    x$TLEN <- as.integer(x$TLEN)
    #message("done!\n")
    return(list("header" = header, "x" = x))
}

line2vals <- function(x, nfields = 11){
    x = strsplit(x, "\t", fixed = TRUE)[[1]]
    c(x[1:nfields], paste(x[-c(1:nfields)], collapse = "\t"))
}
```
