## Supplementary material for "Nano3P-seq: transcriptome-wide analysis of gene expression and tail dynamics using end-capture nanopore cDNA sequencing": FileS2: rectify.R

```
#' Convert negative values in the signal to corresponding positive values.
#'
#' @param data a numeric. A numeric vector containing the raw signal.
#'
#' @return A numeric vector containing the modified signal.
#'
rectify <- function(data){
    data <- abs(data)
    return(data)
}
```
