## Supplementary material for "Nano3P-seq: transcriptome-wide analysis of gene expression and tail dynamics using end-capture nanopore cDNA sequencing": FileS2: sliding-windows-dna.R

```
#' Sliding window from right to left.
#'
#' Applies a function to window that can slide across a timeseries from right to
#' left i.e., from a read's end to its begining.
#' Acknowledgment: evobiR tool
#'
#' @param FUN Function to apply to each window's data
#' @param data Timeseries data to which to apply the sliding window
#' @param window_size Size of the window
#' @param step_size Step-size of the sliding window
#'
#' @return A signal that has been smoothed using the sliding window.
#'
right_to_left_sliding_window_dna <-
    function(FUN, data, window_size, step_size) {
        #Reverse the data first
        data <- rev(data)
        total <- length(data)
        spots <- seq(from = 1, to = (total - window_size), by = step_size)
        result <- vector(length = length(spots))
        for (i in seq_along(spots)) {
            result[i] <- match.fun(FUN)(data[spots[i]:(spots[i] + window_size - 1)])
        }

        data_to_append <- mean(utils::tail(result, n=150))
        result <- c(result, rep(data_to_append, times=(length(data)-length(result))))

        result <- rev(result)
        return(result)
    }


#' Sliding window from left to right
#'
#' Applies a function to window that can slide across a timeseries from left to
#' right, i.e. from a read's beginning to it end.
#' Acknowledgment: evobiR tool
#'
#' @param FUN Function to apply to each window's data
#' @param data Timeseries data to which to apply the sliding window
#' @param window_size Size of the window
#' @param step_size Step-size of the sliding window
#'
#' @return A signal that has been smoothed using the sliding window.
#'
left_to_right_sliding_window_dna <-
    function(FUN, data, window_size, step_size) {
        total <- length(data)
        spots <- seq(from = 1, to = (total - window_size), by = step_size)
        result <- vector(length = length(spots))
        for (i in seq_along(spots)) {
            result[i] <- match.fun(FUN)(data[spots[i]:(spots[i] + window_size - 1)])
        }

        data_to_append <- mean(utils::tail(result, n=400))
        result <- c(result, rep(data_to_append, times=(length(data)-length(result))))
        return(result)
    }
```
