## Supplementary material for "Nano3P-seq: transcriptome-wide analysis of gene expression and tail dynamics using end-capture nanopore cDNA sequencing": FileS2: smoothen-rle-intersections-rna.R

```
#' Smoothen RLE intersections for RNA
#'
#' @param rle_intersections RLE object
#'
#' @return a list
#'

smoothen_rle_intersections_rna <- function(rle_intersections){

    #rle_lengths <-  c(13, 137, 12, 8243, 45, 104, 45, 380)
    #rle_values <- c(TRUE, FALSE, TRUE, FALSE, TRUE, FALSE, TRUE, FALSE)

    rle_lengths <- rle_intersections$lengths
    rle_values <- rle_intersections$values
    new_rle_lengths <- c()
    new_rle_values <- c()
    i = 0
    while (i < (length(rle_lengths)-2)){
        i = i+1
        # If you find a W looing pattern, then merge it to form one
        # contiguous section
        if ((rle_values[i] && !rle_values[i+1] && rle_values[i+2]) &&
            (rle_lengths[i] > 100 && rle_lengths[i+2] > 100 && rle_lengths[i+1] < 250)){
            new_rle_values <- c(new_rle_values, !rle_values[i+1])
            new_rle_lengths <- c(new_rle_lengths, rle_lengths[i]+rle_lengths[i+1]+rle_lengths[i+2])
            i = i + 2 # skip two iterations
        }
        else{
            new_rle_values <- c(new_rle_values, rle_values[i])
            new_rle_lengths <- c(new_rle_lengths, rle_lengths[i])
        }
    }

    # because we stop the loop 2-items early, we need to take them into account
    # and append them to the end
    remaining_items = length(rle_lengths) - i
    if (remaining_items > 0){
        for (k in seq_len(remaining_items)) {
            j = k + i
            new_rle_values <- c(new_rle_values, rle_values[j])
            new_rle_lengths <- c(new_rle_lengths, rle_lengths[j])
        }
    }
    return(list(lengths=new_rle_lengths, values=new_rle_values))
}
```
