## Supplementary material for "Nano3P-seq: transcriptome-wide analysis of gene expression and tail dynamics using end-capture nanopore cDNA sequencing": FileS2: truncate-spikes.R

```
#' Truncate spikes
#'
#' It has been observed that the signal contains a lot of high-voltage spikes
#' distributed throughout the lenght of the read. This function truncates/clips
#' these spikes to a specified threshold value
#'
#' @param data Data that needs its spikes to be truncated
#' @param spike_threshold Threshold beyond which spikes will be clipped
#'
#' @return Same data as input but with spikes clipped
#'
#' @examples
#' \dontrun{
#'
#' truncate_spikes(data, 2)
#' }
#'
truncate_spikes <- function (data, spike_threshold){
    data[data > spike_threshold] = spike_threshold
    data[data < -spike_threshold] = -spike_threshold
    return(data)
}
```
