## Supplementary material for "Nano3P-seq: transcriptome-wide analysis of gene expression and tail dynamics using end-capture nanopore cDNA sequencing": FileS2: z-normalize.R

```
#' z-normalize
#'
#' Apply z-normalization to the time series data
#'
#' @param timeseries Timeseries data that needs to be z-normalized
#'
#' @return z_normalized_timeseries A z-normalized timeseries
#'
z_normalize <- function(timeseries) {
    ts_mean <- mean(timeseries)
    ts_stdev <- stats::sd(timeseries)
    z_normalized_timeseries <- (timeseries - ts_mean)/ts_stdev
    return(z_normalized_timeseries)
}
```
