## Supplementary material for "Nano3P-seq: transcriptome-wide analysis of gene expression and tail dynamics using end-capture nanopore cDNA sequencing": FileS2: README.md

<!-- README.md is generated from README.Rmd. Please edit that file -->

# tailfindr <a href=''><img src='man/figures/tailfindr-logo.png' align="right" height="250" /></a>

<!-- badges: start -->
<!-- badges: end -->

## What is *tailfindr*?

tailfindr is a R package for estimating poly(A)-tail lengths in Oxford
Nanopore reads. This branch is specifically designed for the
[Nano3P-seq](https://www.biorxiv.org/content/10.1101/2021.09.22.461331)
protocol. For all other needs, please install from the
[main](https://github.com/adnaniazi/tailfindr) branch.

## Features of *tailfindr*

-   Works for both RNA and DNA reads.
-   Supports data that has been basecalled with Albacore or Guppy. It
    also support data that has been basecalled using the newer
    ‘flipflop’ model.
-   Can work on single or multi-fast5 file reads.

*tailfindr* has been developed at [Valen
Lab](https://www.cbu.uib.no/valen/) in [Computational Biology
Unit](https://www.cbu.uib.no/) at the [University of
Bergen](https://www.uib.no/), Norway.

## Installation

#### Step 1. Installing VBZ plugin

VBZ is a compression format used by Nanopore to compress the data inside
a FAST5 file. You must install VBZ plugin for your operating system so
that tailfindr can successfully read FAST5 files. Download and install
VBZ plugin from
[here](https://github.com/nanoporetech/vbz_compression/releases).

#### Step 2. Installing HDF5 library

tailfindr depends on the HDF5 library for reading Fast5 files. For OS X
and Linux, the HDF5 library needs to be installed via one of the (shell)
commands specified below:

| System                                      | Command                            |
|:--------------------------------------------|:-----------------------------------|
| **OS X (using Homebrew)**                   | `brew install hdf5`                |
| **Debian-based systems (including Ubuntu)** | `sudo apt-get install libhdf5-dev` |
| **Systems supporting yum and RPMs**         | `sudo yum install hdf5-devel`      |

HDF5 1.8.14 has been pre-compiled for Windows and is available
[here](https://github.com/mannau/h5-libwin) — thus no manual
installation is required.

#### Step 3. Installing devtools

Currently, *tailfindr* is not listed on CRAN/Bioconductor, so you need
to install it using `devtools`. To install `devtools` use the following
command in R/R-studio:

``` r
install.packages("devtools")
```

#### Step 4. Installing rbokeh

rbokeh has been delisted from CRAN and therefore tailfindr cannot
install itself. You need to manually install it yourself using the
command below:

``` r
devtools::install_url('https://cran.r-project.org/src/contrib/Archive/rbokeh/rbokeh_0.5.1.tar.gz', type = "source", dependencies = TRUE)
```

#### Step 5. Installing tailfindr

Now you can install tailfindr using the command below in R/R-studio:

``` r
devtools::install_github("adnaniazi/tailfindr", ref='nano3p-seq')
```

If you also want to build the vignette while installing tailfindr, then
run the command below:

``` r
remotes::install_github('adnaniazi/tailfindr', ref='nano3p-seq', build = TRUE, build_opts = c("--no-resave-data", "--no-manual"), force = TRUE)
```

Now you are ready to use *tailfindr*.

## Usage

#### 1. Minimal working example

`find_tails()` is the main function that you can use to find tail
lengths in both RNA and DNA reads. It saves a CSV file containing all
the tail-length data. Furthermore, it also returns the same data as a
tibble.

Give below is a minimal use case in which we will run tailfindr on
example RNA reads present in the tailfindr package.

``` r
library(tailfindr)
df <- find_tails(fast5_dir = system.file('extdata', 'rna', package = 'tailfindr'),
                 save_dir = '~/Downloads',
                 csv_filename = 'rna_tails.csv',
                 num_cores = 2)
```

In the above example, *tailfindr* returns a tibble containing the tail
data which is then stored in the variable `df`. tailfindr also saves
this dataframe as a csv file (`rna_tails.csv`) in the user-specified
`save_dir`, which in this case is set to `~/Downloads`. A logfile is
also saved in the `save_dir`. The parameter `num_cores` can be increased
depending on the number of *physical* cores at your disposal.

#### 2. Plotting the tail

Additionally, *tailfindr* allows you to generate plots that show the
tail location in the raw squiggle. You can save these plots as
interactive `.html` files by using `'rbokeh'` as the `plotting_library`.
You can zoom in on the tail region in the squiggle and see the exact
location of the tail.

Give below is a minimal use case in which we will run tailfindr on
example cDNA reads present in the tailfindr package, and also save the
plots:

``` r
df <- find_tails(fast5_dir = system.file('extdata', 'cdna', package = 'tailfindr'),
                 save_dir = '~/Downloads',
                 csv_filename = 'cdna_tails.csv',
                 num_cores = 2,
                 save_plots = TRUE,
                 plotting_library = 'rbokeh')
```

![Poly(T) read squiggle
plot](https://github.com/adnaniazi/tailfindr/raw/master/man/figures/poly_t_without_debug.gif)

However, note that generating plots can slow down the performance of
tailfindr. We recommend that you generate these plots only for a small
subset of your reads.

#### 3. Plotting the tail and debug traces

*tailfindr* can plot additional information that it used while deriving
the tail boundaries. Please read our preprint to learn how *tailfindr*
works. To plot this information, set the `plot_debug_traces` parameter
to `TRUE`.

``` r
df <- find_tails(fast5_dir = system.file('extdata', 'cdna', package = 'tailfindr'),
                 save_dir = '~/Downloads',
                 csv_filename = 'cdna_tails.csv',
                 num_cores = 2,
                 save_plots = TRUE,
                 plot_debug_traces = TRUE,
                 plotting_library = 'rbokeh')
```

![Poly(A) read squiggle
plot](https://github.com/adnaniazi/tailfindr/raw/master/man/figures/poly_a_with_debug.gif)

#### 4. Specifying custom basecall group

*tailfindr* needs `Fastq` and `Events/Move` table to work on. By
default, it searches for them in the `Basecall_1D_000` group in the
Analyses section of the FAST5 file. If for whatever reason, you need
*tailfindr* to read data from another basecall group – lets say
`Basecall_1D_001` – then you can run *tailfindr* as below:

``` r
df <- find_tails(fast5_dir = system.file('extdata', 'rna_basecall_1D_001', package = 'tailfindr'),
                 save_dir = '~/Downloads',
                 csv_filename = 'rna_tails.csv',
                 num_cores = 2,
                 basecall_group = 'Basecall_1D_001',
                 save_plots = TRUE,
                 plot_debug_traces = TRUE,
                 plotting_library = 'rbokeh')
```

In this case, the input FAST5 have two basecall groups:
`Basecall_1D_000` and `Basecall_1D_001` but we configured *tailfindr* to
use `Events/Move` table from the `Basecall_1D_001` group.

There are more options available in the find_tails() function. Please
see its
[documentation](https://rdrr.io/github/adnaniazi/tailfindr/man/find_tails.html).

#### 5. Specifying custom cDNA primers

If you have used custom front and end primers while designing you cDNA
sequences, you can now specify them in *tailfindr*. *tailfindr* will use
these sequences instead of the defaults ones to find the orientation of
the reads and one of the ends of the poly(A)/(T) tail. Here is how you
call *tailfindr* if you have used custom primers:

``` r
df <- find_tails(fast5_dir = system.file('extdata', 'cdna', package = 'tailfindr'),
                 save_dir = '~/Downloads/tailfindr_output',
                 csv_filename = 'cdna_tails.csv',
                 num_cores = 4,
                 dna_datatype = 'custom-cdna',
                 front_primer = "TTTCTGTTGGTGCTGATATTGCTGCCATTACGGCCGG",
                 end_primer = "ACTTGCCTGTCGCTCTATCTT")
```

Important thing to note here is the use of three additional parameters:
`dna_datatype`, `front_primer`, and `end_primer`.

`front_primer` and `end_primer` sequences should always be specified in
the 5’ to 3’ direction.

![cDNA](https://github.com/adnaniazi/tailfindr/raw/master/man/figures/cdna_construct.png)

### Description of the CSV/Dataframe columns

*tailfindr* returns tail data in a dataframe and also saves this
information in a user-specified CSV file. The columns generated depend
on the whether *tailfindr* was run on RNA or DNA data. Below is a
description of columns for both thses scenarios:

##### When input data is RNA

| Column Names   | Datatype  | Description                                                                                                      |
|:---------------|:----------|:-----------------------------------------------------------------------------------------------------------------|
| read_id        | character | Read ID as given in the Fast5 file                                                                               |
| tail_start     | numeric   | Sample index of start site of the tail in raw data                                                               |
| tail_end       | numeric   | Sample index of end site of the tail in raw data                                                                 |
| samples_per_nt | numeric   | Read rate in terms of samples per nucleotide                                                                     |
| tail_length    | numeric   | Tail length in nucleotides. It is the difference between `tail_end` and `tail_start` divided by `samples_per_nt` |
| file_path      | character | Absolute path of the Fast5 file                                                                                  |

##### When input data is DNA

Here are the columns that you will get from *tailfindr* if you have run
it on DNA data:

| Column Names   | Datatype         | Description                                                                                                                                                                                                                                                                                                                                                           |
|----------------|------------------|-----------------------------------------------------------------------------------------------------------------------------------------------------------------------------------------------------------------------------------------------------------------------------------------------------------------------------------------------------------------------|
| read_id        | character        | Read ID as given in the Fast5 file                                                                                                                                                                                                                                                                                                                                    |
| read_type      | character factor | `"no_adaptor_found"` if the Nano3P-seq adaptor is not found with high-confidence, `"contains_no_polyT_tail"` if the Nano3P-seq adaptor was found with high-confidence but no poly(T) was found next to it so the read has no poly(T) tail, and `"contains_a_polyT_tail"` if the Nano3P-seq adaptor was found with high-confidence and a poly(T) was found next to it. |
| tail_start     | numeric          | Sample index of start site of the tail in raw data                                                                                                                                                                                                                                                                                                                    |
| tail_end       | numeric          | Sample index of end site of the tail in raw data                                                                                                                                                                                                                                                                                                                      |
| samples_per_nt | numeric          | Read rate in terms of samples per nucleotide                                                                                                                                                                                                                                                                                                                          |
| tail_length    | numeric          | Tail length in nucleotides. It is the difference between `tail_end` and `tail_start` divided by `samples_per_nt`                                                                                                                                                                                                                                                      |
| file_path      | character        | Absolute path of the Fast5 file                                                                                                                                                                                                                                                                                                                                       |

## The devil👹 in the details

-   tailfindr needs the `Events/Move` table in the FAST5 file to
    calculate the read-specific normalizer – `samples_per_nt` – which is
    used to convert tail length in samples to tail length in
    nucleotides. If your data was basecalled with
    *MinKNOW-Live-Basecalling*, then the Events/Move table might not be
    saved in the FAST5 file. In such a case, you can rebasecall your
    reads and adjust the `basecall_group` parameter accordingly when
    calling `find_tails()` function as demonstrated in the use case # 4
    above. This is because now the Events/Move table will now be under
    `Basecall_1D_001` instead of *tailfindr’s* default search location
    `Basecall_1D_000`. See the figure below: The panel on left shows
    that the MinKNOW live basecalled read; it has no Event/Move table.
    The panel on the right shows the same read after it has been
    re-basecalled using standalone Guppy. Now there is Event/Move table
    under the freshly-added basaecall group (`Basecall_1D_001`).
    `find_tails()` should be called with `basecall_group` set to
    `"Basecall_1D_001"` as shown in the use case # 4 above.

![MinKNOW Live Basecalling
problem](https://github.com/adnaniazi/tailfindr/raw/master/man/figures/minkow_live_basecalling.png)

-   For DNA data, *tailfindr* decides whether a read is poly(A) or
    poly(T) based on finding Nanopore primers/adaptors. If you are using
    the flipflop model to basecall DNA data, please ensure that the
    nanopore adaptors are not trimmed off while basecalling. This can be
    done by turning off `enabling_trimming` option in the basecalling
    script. The script below shows you how we have basecalled our reads
    using the flipflop model

``` bash
#!/bin/sh
INPUT=/raw/fast5/files/path/
OUTPUT=/output/folder/path/
guppy_basecaller \
    --config dna_r9.4.1_450bps_flipflop.cfg \
    --input $INPUT \
    --save_path $OUTPUT \
    --recursive \
    --fast5_out \
    --hp_correct 1 \
    --disable_pings 1 \
    --enable_trimming 0 
```

-   *tailfindr* has been tested and validated using the following
    sequencing kits:

1.  SQK-RNA001 and SQK-RNA002 for RNA
2.  SQK-LSK108 and SQK-LSK109 for DNA
3.  SQK-PCS110 for PCR cDNA

-   *tailfindr* has been tested and validated using the basecallers:

1.  Albacore v2.3.1 and v2.3.3.
2.  Guppy v2.2.2, v2.3.1, v3.0.1. Guppy v2.2.2, v2.3.1 were tested with
    flipflop basecalling for DNA, and Guppy v3.0.1 was tested with
    flipflop basecalling for RNA.

## Want to know more?

We have written a book chapter on how you can use *tailfindr* for
transcript isofrom-specific poly(A) tail profiling. It has a detailed
protocol and analysis pipeline for processing the data through
*tailfindr* .The chapter can be downloaded from
[here](https://adnaniazi.com/website-assets/niazi_and_krause_2019_springer_book_chapter.pdf).

## Getting help

If you encounter a clear bug, please file a minimal reproducible example
on [github](https://github.com/adnaniazi/tailfindr/issues). Please do
provide us a few reads (around 10) so that we can reproduce the problem
at our end, and figure out a solution for you.

## Citation

Krause, M., Niazi, A. M., Labun, K., Cleuren, Y. N. T., Müller, F. S., &
Valen, E. (2019). *tailfindr*: alignment-free poly(A) length measurement
for Oxford Nanopore RNA and DNA sequencing. Rna, 25(10), 1229–1241. doi:
10.1261/rna.071332.119

## License

And of course:

GPL-3: <https://www.gnu.org/licenses/gpl-3.0.en.html>
