## Supplementary material for "Nano3P-seq: transcriptome-wide analysis of gene expression and tail dynamics using end-capture nanopore cDNA sequencing": FileS2: test-rectify.R

```
context("Rectify")

test_that("Rectification(flipping) is working correctly", {
    data <- c(1, 2, 3, 4, 5, 0, -1, -2, -3, -4, -5)
    expected <- c(1, 2, 3, 4, 5, 0, 1, 2, 3, 4, 5)
    expect_equal(rectify(data), expected)
})
```
