## Supplementary material for "Nano3P-seq: transcriptome-wide analysis of gene expression and tail dynamics using end-capture nanopore cDNA sequencing": FileS2: test-truncate-spikes.R

```
context("Truncate spike")

test_that("testing truncate spikes", {
    data <- c(1, 1, 2, 2, 3, 3, 4, 4, -1, -1, -2, -2, -3, -3, -4, -4)
    expected_data <- c(1, 1, 2, 2, 2, 2, 2, 2, -1, -1, -2, -2, -2, -2, -2, -2)
    expect_equal(truncate_spikes(data, 2), expected_data)

    data <- c(1, 1, 2, 2, 3, 3, 4, 4)
    expected_data <- c(1, 1, 2, 2, 2, 2, 2, 2)
    expect_equal(truncate_spikes(data, 2), expected_data)

    data <- c( -1, -1, -2, -2, -3, -3, -4, -4)
    expected_data <- c(-1, -1, -2, -2, -2, -2, -2, -2)
    expect_equal(truncate_spikes(data, 2), expected_data)

})
```
