## Supplementary material for "Nano3P-seq: transcriptome-wide analysis of gene expression and tail dynamics using end-capture nanopore cDNA sequencing": FileS2: test-z-normalize.R

```
context("Z Normalization")

test_that("Normalization is working correctly", {
    expect_equal(z_normalize(c(1,2,3,4,5)),
                 c(-1.2649111, -0.6324555,  0.0000000,  0.6324555 , 1.2649111),
                 tolerance=1e-3)
})
```
