## Supplementary material for "Nano3P-seq: transcriptome-wide analysis of gene expression and tail dynamics using end-capture nanopore cDNA sequencing": FileS2: tailfindr.Rmd

`````
---
title: "Introduction to tailfindr"
author: "Adnan M. Niazi"
date: "`r Sys.Date()`"
output: rmarkdown::html_vignette
vignette: >
  %\VignetteIndexEntry{Introduction to tailfindr}
  %\VignetteEngine{knitr::rmarkdown}
  %\VignetteEncoding{UTF-8}
fig_width: 6 
fig_height: 2 
---

```{r setup, include = FALSE}
knitr::opts_chunk$set(
  collapse = TRUE,
  comment = "#>"
  )
```
## What is tailfindr?

tailfindr is a R package for estimating poly(A)-tail lengths in Oxford Nanopore reads. 

## Features of tailfindr

* Works for both RNA and DNA reads. In the case of DNA reads, it estimates both poly(A)- and poly(T)-tail lengths.
* Supports data that has been basecalled with Albacore or Guppy. It also support data that has been basecalled using the newer 'flipflop' model.
* Can work on single or multi-fast5 file reads.

tailfindr has been developed at [Valen Lab](https://www.cbu.uib.no/valen/) in [Computational Biology Unit](https://www.cbu.uib.no/) at the [University of Bergen](https://www.uib.no/), Norway.

## Installation

#### Step 1. Installing HDF5 library

tailfindr depends on the HDF5 library for reading Fast5 files. For OS X and Linux, the HDF5 library needs to be installed via one of the (shell) commands specified below:

| System                                    | Command
|:------------------------------------------|:---------------------------------|
|**OS X (using Homebrew)**                  | `brew install hdf5`
|**Debian-based systems (including Ubuntu)**| `sudo apt-get install libhdf5-dev` 
|**Systems supporting yum and RPMs**        | `sudo yum install hdf5-devel`

HDF5 1.8.14 has been pre-compiled for Windows and is available [here](https://github.com/mannau/h5-libwin) &mdash; thus no manual installation is required.

#### Step 2. Installing devtools
Currently, tailfindr is not listed on CRAN/Bioconductor, so you need to install it using `devtools`. To install `devtools` use the following command:

```{r, message = FALSE, eval = FALSE}
install.packages("devtools")
```

#### Step 3. Installing tailfindr

Now you can install tailfindr using the command below:
````{r, message = FALSE, eval = FALSE}
devtools::install_github("adnaniazi/tailfindr")
```

Now you are ready to use tailfindr.

## Usage

#### 1. Minimal working example

`find_tails()` is the main function that you can use to find tail lengths in both RNA and DNA reads. It saves a CSV file containing all the tail-length data. Furthermore, it also returns the same data as a tibble. 

Give below is a minimal use case in which we will run tailfindr on example RNA reads present in the tailfindr package. 

```{r, message = FALSE, eval = FALSE}
library(tailfindr)
df <- find_tails(fast5_dir = system.file('extdata', 'rna', package = 'tailfindr'),
                 save_dir = '~/Downloads',
                 csv_filename = 'rna_tails.csv',
                 num_cores = 2)
```
In the above example, tailfindr returns a tibble containing the tail data which is then stored in the variable `df`. tailfindr also saves this dataframe as a csv file (`rna_tails.csv`) in the user-specified `save_dir`, which in this case is set to `~/Downloads`. A logfile is also saved in the `save_dir`. The parameter `num_cores` can be increased depending on the number of *physical* cores at your disposal.

#### 2. Plotting the tail

Additionally, tailfindr allows you to generate plots that show the tail location in the raw squiggle. You can save these plots as interactive `.html` files by using `'rbokeh'` as the `plotting_library`. You can zoom in on the tail region in the squiggle and see the exact location of the tail.

Give below is a minimal use case in which we will run tailfindr on example cDNA reads present in the tailfindr package, and also save the plots:

```{r, message = FALSE, eval = FALSE}
df <- find_tails(fast5_dir = system.file('extdata', 'cdna', package = 'tailfindr'),
                 save_dir = '~/Downloads',
                 csv_filename = 'cdna_tails.csv',
                 num_cores = 2,
                 save_plots = TRUE,
                 plotting_library = 'rbokeh')
```

However, note that generating plots can slow down the performace of tailfindr. We recommend that you generate these plots only for a small subset of your reads.

#### 3. Plotting the tail and debug traces

tailfindr can plot additional information that it used while deriving the tail boundaries. Please read our preprint to learn how tailfindr works. To plot this information, set the `plot_debug_traces` parameter to `TRUE`.
```{r, message = FALSE, eval = FALSE}
df <- find_tails(fast5_dir = system.file('extdata', 'cdna', package = 'tailfindr'),
                 save_dir = '~/Downloads',
                 csv_filename = 'cdna_tails.csv',
                 num_cores = 2,
                 save_plots = TRUE,
                 plot_debug_traces = TRUE,
                 plotting_library = 'rbokeh')
```


#### 4. Specifying custom basecall group
tailfindr needs `Fastq` and `Events/Move` table to work on. By default, it searches for them in the `Basecall_1D_000` group in the Analyses section of the FAST5 file. If for whatever reason, you need tailfindr to read data from another basecall group -- lets say `Basecall_1D_001` -- then you can run tailfindr as below:
```{r, message = FALSE, eval = FALSE}
df <- find_tails(fast5_dir = system.file('extdata', 'rna_basecall_1D_001', package = 'tailfindr'),
                 save_dir = '~/Downloads',
                 csv_filename = 'rna_tails.csv',
                 num_cores = 2,
                 basecall_group = 'Basecall_1D_001',
                 save_plots = TRUE,
                 plot_debug_traces = TRUE,
                 plotting_library = 'rbokeh')
```
In this case, the input FAST5 have two basecall groups: `Basecall_1D_000` and `Basecall_1D_001` but we configured tailfindr to use `Events` table from the `Basecall_1D_001` group.

There are more options available in the find_tails() function. Please see its [documentation](https://rdrr.io/github/adnaniazi/tailfindr/man/find_tails.html). 

### Description of the CSV/Dataframe columns
tailfindr returns tail data in a dataframe and also saves this information in a user-specified CSV file. The columns generated depend on the whether tailfindr was run on RNA or DNA data. Below is a description of columns for both thses scenarios:

##### When input data is RNA

| Column Names   | Datatype  | Description                                                                                                |
|:---------------|:----------|:-----------------------------------------------------------------------------------------------------------|
| read_id        | character | Read ID as given in the Fast5 file                                                                         |
| tail_start     | numeric   | Sample index of start site of the tail in raw data                                                         |
| tail_end       | numeric   | Sample index of end site of the tail in raw data                                                           |
| samples_per_nt | numeric   | Read rate in terms of samples per nucleotide                                                               |
| tail_length    | numeric   | Tail length in nucleotides. It is the difference between `tail_end` and `tail_start` divided by `samples_per_nt` |
| file_path      | character | Absolute path of the Fast5 file                                                                            |

##### When input data is DNA
Here are the columns that you will get from tailfindr if you have run it on DNA data:

| Column Names   | Datatype         | Description                                                                                                                                                                       |
|----------------|------------------|-----------------------------------------------------------------------------------------------------------------------------------------------------------------------------------|
| read_id        | character        | Read ID as given in the Fast5 file                                                                                                                                                |
| read_type      | character factor | Whether a read is `"polyA"`, `"polyT"`, or `"invalid"`. Invalid reads are those in which tailfindr wasn't able to find Nanopore primers with high confidence.                           |
| tail_is_valid  | logical          | Whether a poly(A) tail is a full-length read or not. This is important because a poly(A) tail is at the end of the read, and premature termination of reads is prevelant in cDNA. |
| tail_start     | numeric          | Sample index of start site of the tail in raw data                                                                                                                                |
| tail_end       | numeric          | Sample index of end site of the tail in raw data                                                                                                                                  |
| samples_per_nt | numeric          | Read rate in terms of samples per nucleotide                                                                                                                                      |
| tail_length    | numeric          | Tail length in nucleotides. It is the difference between `tail_end` and `tail_start` divided by `samples_per_nt`                                                                        |
| file_path      | character        | Absolute path of the Fast5 file                                                                                                                                                   |
## The devil👹 in the details

* tailfindr needs the `Events/Move` table in the FAST5 file to calculate the read-specific normalizer -- `samples_per_nt` -- which is used to convert tail length in samples to tail length in nucleotides. If your data was basecalled with *MinKNOW-Live-Basecalling*, then the Events/Move table might not be saved in the FAST5 file. In such a case, you can rebasecall your reads and adjust the `basecall_group` parameter accordingly in the `find_tails()` function as demonstrated in the use case # 4 above.

* For DNA data, tailfindr decides whether a read is poly(A) or poly(T) based on finding Nanopore primers/adaptors. If you are using the flipflop model to basecall DNA data, please ensure that the nanopore adaptors are not trimmed off while basecalling. This can be done by turning off `enabling_trimming` option in the basecalling script. The script below shows you how we have basecalled our reads using the flipflop model

```{r, engine='bash', message = FALSE, eval = FALSE}
#!/bin/sh
INPUT=/raw/fast5/files/path/
OUTPUT=/output/folder/path/
guppy_basecaller \
    --config dna_r9.4.1_450bps_flipflop.cfg \
    --input $INPUT \
    --save_path $OUTPUT \
    --recursive \
    --fast5_out \
    --hp_correct 1 \
    --disable_pings 1 \
    --enable_trimming 0 
```

## Getting help

If you encounter a clear bug, please file a minimal reproducible example on [github](https://github.com/adnaniazi/tailfindr/issues). FPlease do provide us a few reads (around 10) so that we can reproduce the problem at our end, and figure out a solution for you.

# Citation
Maximilian Krause, Adnan M. Niazi, Kornel Labun, Florian Sebastian Müller, Yamila Nicole Torres Cleuren, Eivind Valen (2019): ***tailfindr*: Alignment-free poly(A) length measurement for Oxford Nanopore RNA and DNA sequencing**. bioRxiv 588343; doi: https://doi.org/10.1101/588343

## License

And of course:

GPL-3: https://www.gnu.org/licenses/gpl-3.0.en.html
`````
