## Supplementary figures and images for "Nano3P-seq: transcriptome-wide analysis of gene expression and tail dynamics using end-capture nanopore cDNA sequencing"

### cdna_construct.png

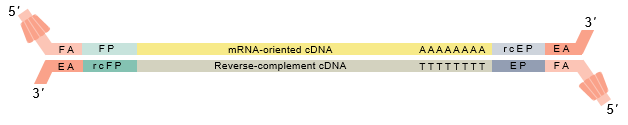

### minkow_live_basecalling.png

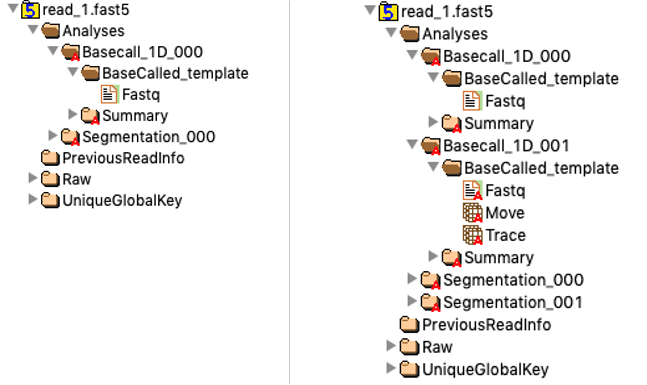

### poly_a_with_debug.gif

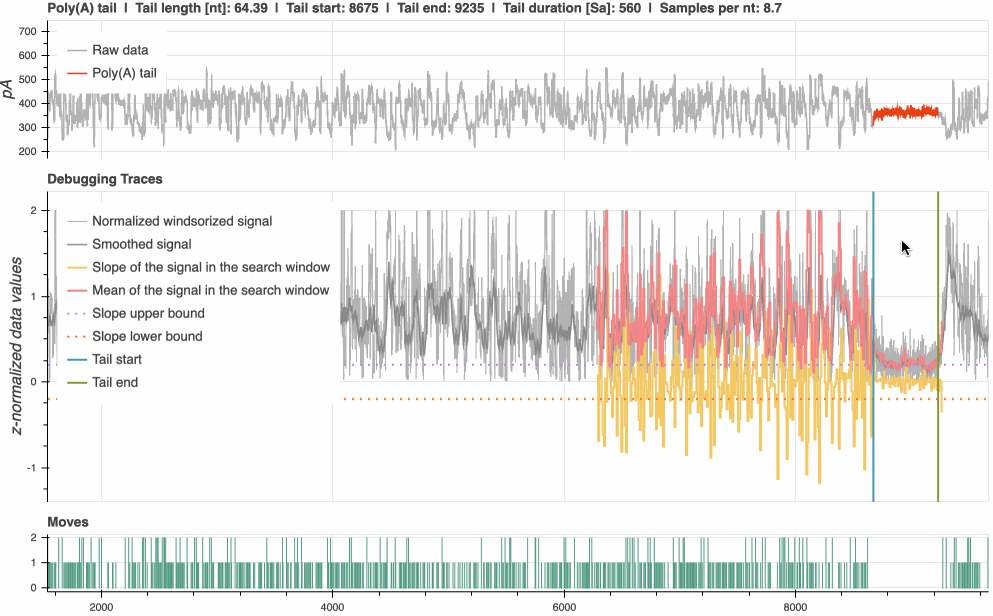

### poly_t_without_debug.gif

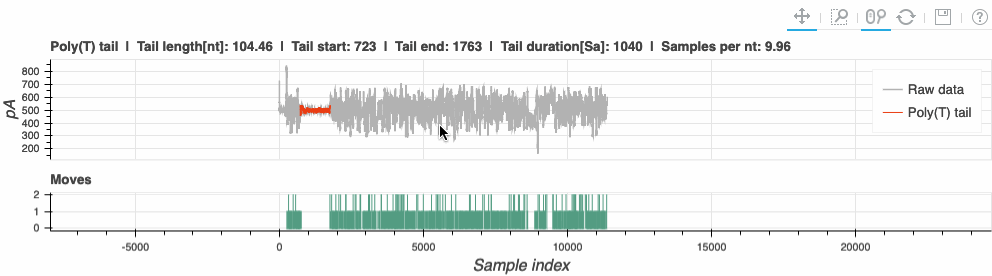

### tailfindr-logo.png

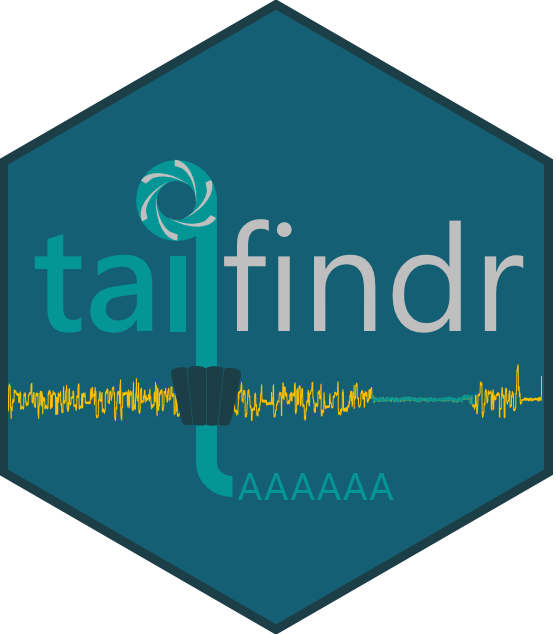
